# Tissue-ABPP enables high-resolution confocal fluorescence imaging of serine hydrolase activity in cryosections – Application to glioma brain unveils activity hotspots originating from tumor-associated neutrophils

**DOI:** 10.1101/783704

**Authors:** Niina Aaltonen, Prosanta K. Singha, Hermina Jakupović, Thomas Wirth, Haritha Samaranayake, Sanna Pasonen-Seppänen, Kirsi Rilla, Markku Varjosalo, Laura E. Edgington-Mitchell, Paulina Kasperkiewicz, Marcin Drag, Sara Kälvälä, Eemeli Moisio, Juha R. Savinainen, Jarmo T. Laitinen

## Abstract

Serine hydrolases (SHs) are a functionally diverse family of enzymes playing pivotal roles in health and disease and have emerged as important therapeutic targets in many clinical conditions. Activity-based protein profiling (ABPP) using fluorophosphonate (FP) probes has been a powerful chemoproteomic approach in studies unveiling roles of SHs in various biological systems. The ABPP approach utilizes cell/tissue proteomes and features the FP warhead, linked to a fluorescent reporter for in-gel fluorescence imaging or a biotin tag for streptavidin enrichment and LC-MS/MS-based target identification. Here, we advance the ABPP methodology to glioma brain cryosections, enabling high-resolution confocal fluorescence imaging of SH activity in different cell types of the tumor microenvironment, identified by using extensive immunohistochemistry on activity probe labeled sections. We name this technique tissue-ABPP to distinguish it from conventional gel-based ABPP. We show heightened SH activity in glioma vs. normal brain and unveil activity hotspots originating from tumor-associated neutrophils. Thorough optimization and validation is provided by parallel gel-based ABPP combined with LC-MS/MS-based target verification. Tissue-ABPP enables a wide range of applications for confocal imaging of SH activity in any type of tissue or animal species.

## Introduction

The serine hydrolases (SHs) form a diverse family of enzymes with a predicted number of ∼240 in humans, falling into two subfamilies: the serine proteases (∼125 members) and the metabolic SHs (mSHs ∼115 members) (Long and Cravatt, 2011; Simon and Cravatt, 2010). The mSHs include small-molecule hydrolases, such as lipases, esterases and amidases and utilize a conserved serine nucleophile to hydrolyze amide, ester, and thioester bonds in different types of substrates including metabolites, lipids and peptides. Importantly, SHs have emerged as therapeutic targets in diseases such as cancer, obesity, diabetes, and neurological diseases (Bachovchin and Cravatt, 2012).

The advent of chemoproteomic techniques some 20 years ago, activity-based protein profiling (ABPP) in particular, allowed for the first time proteome-wide profiling of SH activity in cells and tissue homogenates (Liu et al., 1999). The prototype activity probe for SHs features the active site-targeted warhead, typically a fluorophosphonate (FP), linked to a fluorescent reporter allowing in-gel imaging of SH activity in proteomes after SDS-PAGE separation. An advanced platform combining ABPP and multidimensional protein identification techniques (ABPP-MudPIT) was introduced to facilitate high-content functional proteomics discovery of potential new markers of human diseases (Jessani et al., 2005).

In the comparative mode, ABPP enables comparison of SH activity pattern between different proteomes, e.g. aggressive vs. non-aggressive cancer cells (Nomura et al., 2010). Indeed, such studies have highlighted previously unrecognized importance of SH family members as metabolic nodes orchestrating the availability of lipids involved in oncogenic signaling.

In the competitive mode, ABPP has proven its power in the discovery and selectivity testing of novel inhibitors targeting individual SHs (Niphakis and Cravatt, 2014). In this approach, the proteome is first treated with the inhibitor (commonly a serine-nucleophile targeting covalent inhibitor), binding of the inhibitor masks the active site, preventing subsequent labeling with the activity probe. EnPlex, an advanced high-throughput platform based on ABPP principles has been introduced as a feasible approach for SH superfamily-wide selectivity profiling that could be incorporated into the early stage of drug discovery (Bachovchin et al., 2014).

ABPP can also serve a powerful platform for mass-spectrometry-based target identification from complex proteomes. In this case, the FP-warhead is linked to a biotin tag, enabling streptavidin enrichment and subsequent LC/MS-MS-based target identification.

Thus, ABPP has emerged as a powerful method for enzymology and drug discovery, enabling studies of the biological activities of SHs in native biological systems, as well as the discovery of selective inhibitors. Current ABPP approaches characterize SHs based on mobility in gel or MS-based target identification and cannot reveal the identity of the cell-types responsible for an individual SH activity originating from complex proteomes, suggesting that the full potential of this technology may not have yet been harnessed. We reasoned that ABPP-based imaging of SH activity should be feasible at high resolution in complex native proteomes such as brain tissue while preserving the anatomical details of this delicate organ. Brain cryosections serve as fascinating premise to directly explore this issue as functional responses such as receptor-stimulated G protein activity can be readily monitored in brain cryosections without compromising the anatomical integrity (Aaltonen et al., 2013; Sim et al., 1995).

To date, quenched activity probe-based live cell fluorescence imaging has been described for a handful of proteases, such as cysteine cathepsins (Edgington-Mitchell et al., 2017). The quenched fluorescent activity probes were applied for labeling of active proteases in fresh-frozen cancer tissues (Withana et al., 2016), for macrophage detection in atherosclerotic plaques (Abd-Elrahman et al., 2016), and as a diagnostic tool to image skin tumor margins (Liu et al., 2019). Recently, a small-molecule chemical toolbox was constructed for parallel imaging of human neutrophil serine proteases (NSPs). The toolbox comprised activity-probes with different fluorophores showing minimal wavelength overlap and highly selective natural and unnatural amino acid recognition sequences tailored for the four individual NSPs (Kasperkiewicz et al., 2017). This elegant approach enabled for the first time simultaneous imaging of the four NSPs in living neutrophils by fluorescence microscopy.

Here, we advance the ABPP methodology for brain cryosections, enabling high-resolution confocal fluorescence imaging of SH activity in glioma brain sections with well-preserved cyto-architecture. We name this approach tissue-ABPP. We unveil heightened SH activity in the tumor, as compared to healthy brain. Heterogeneous distribution pattern of SH activity within the tumor microenvironment was explored in detail by identification of the glioma-associated cell types using immunohistochemical markers. Cross-validation was provided by classical gel-based ABPP using homogenates of healthy brain and glioma, as well as by LC/MS-MS-based target identification. These studied revealed quite unexpectedly that SH activity hotspots in glioma originate from tumor-associated neutrophils (TANs), rather than tumor-associated macrophages (TAMs). We anticipate that tissue-ABPP enables a wide range of applications for high-resolution confocal imaging of SH activity in practically any type of tissue and animal species, opening new avenues for cellular and subcellular localization of SH activity in complex proteomes with preserved anatomy.

## Results

The principal motivation for this work was to extend the utility of current chemoproteomic methodology, ABPP in particular, for applications aiming at high-resolution imaging of SH activity in glioma brain sections retaining delicate cyto-architecture of the tumor microenvironment. The tissue- and gel-based ABPP approaches that were utilized in this study are illustrated in Figure S1.

### The glioma model

Glioblastoma multiforme (GBM) is the most malignant and most frequent brain tumor accounting for more than 65 % of all cases (Ohgaki and Kleihues, 2009). As the name implies, GBMs have a wide spectrum of histological morphologies ranging from small-cell type to very pleomorphic giant-cell forms with poor differentiation and gliosarcomas (Claes et al., 2007; Ohgaki and Kleihues, 2009). We used the rat syngeneic BT4C gliosarcoma model (Barth and Kaur, 2009) in which the glioma grows in a sarcomatous pattern and is typically composed of a pleomorphic population of tumor cells (Figure S2). We imaged tumors in vivo 22-24 days after implantation using MRI (Figure S3) and animals were sacrificed 5-11 days later. Brain tissue was excised from saline-perfused animals and used for cryosectioning (tissue-ABPP) or preparation of homogenates that were used in cross-validation experiments by gel-ABPP. Animals of both sexes were used.

### Optimization and validation of ABPP protocol for brain cryosections

As our ultimate goal was to achieve high-resolution imaging of SH activity in cryosections without significantly compromising anatomical integrity, we considered fixation necessary to preserve delicate cyto-architecture. On the other hand, fixation should be sufficiently mild in order to preserve enzymatic activity, a prerequisite for the probe to covalently label catalytically competent SHs. We found that in contrast to acetone or methanol, fixation with paraformaldehyde preserved tissue integrity and inhibitor sensitivity of TAMRA-FP labeling (Figure S4).

We chose 0.5 µM probe concentration as a compromise between signal intensity and cost-affordable probe amount (Figure S5). Concerning assay buffer, we found that for applications where maximal TAMRA-FP signal is the desired final readout, Tris or phosphate buffer (pH 7.4) without BSA supplementation is optimal. However, for applications where TAMRA-FP labeling step is followed by immunohistochemistry, BSA is included to block tissue prior to antibody addition (Figure S6).

### Distribution and overall characteristics of TAMRA-FP signal in glioma and control brain

As shown in Figure 1, relatively intense and heterogeneously distributed TAMRA-FP labeling was evident over the glioma. In general, TAMRA-FP fluorescence was more intense over the glioma as compared to most regions of the healthy brain. Nuclear DAPI staining indicated dense cell population in glioma as compared to most regions of the healthy brain. This likely partly accounts for the heightened TAMRA-FP signal in glioma. Intense TAMRA-FP labeling was evident also in cell-dense structures of the healthy brain, such as hippocampal pyramidal cell layer and granular layer of dentate gyrus (Figure 1a) and cerebellar Purkinje cell layer (Figure S6). On the other hand, white matter tracts (Figure 1a) showed low TAMRA-FP labeling.

**Figure 1.**
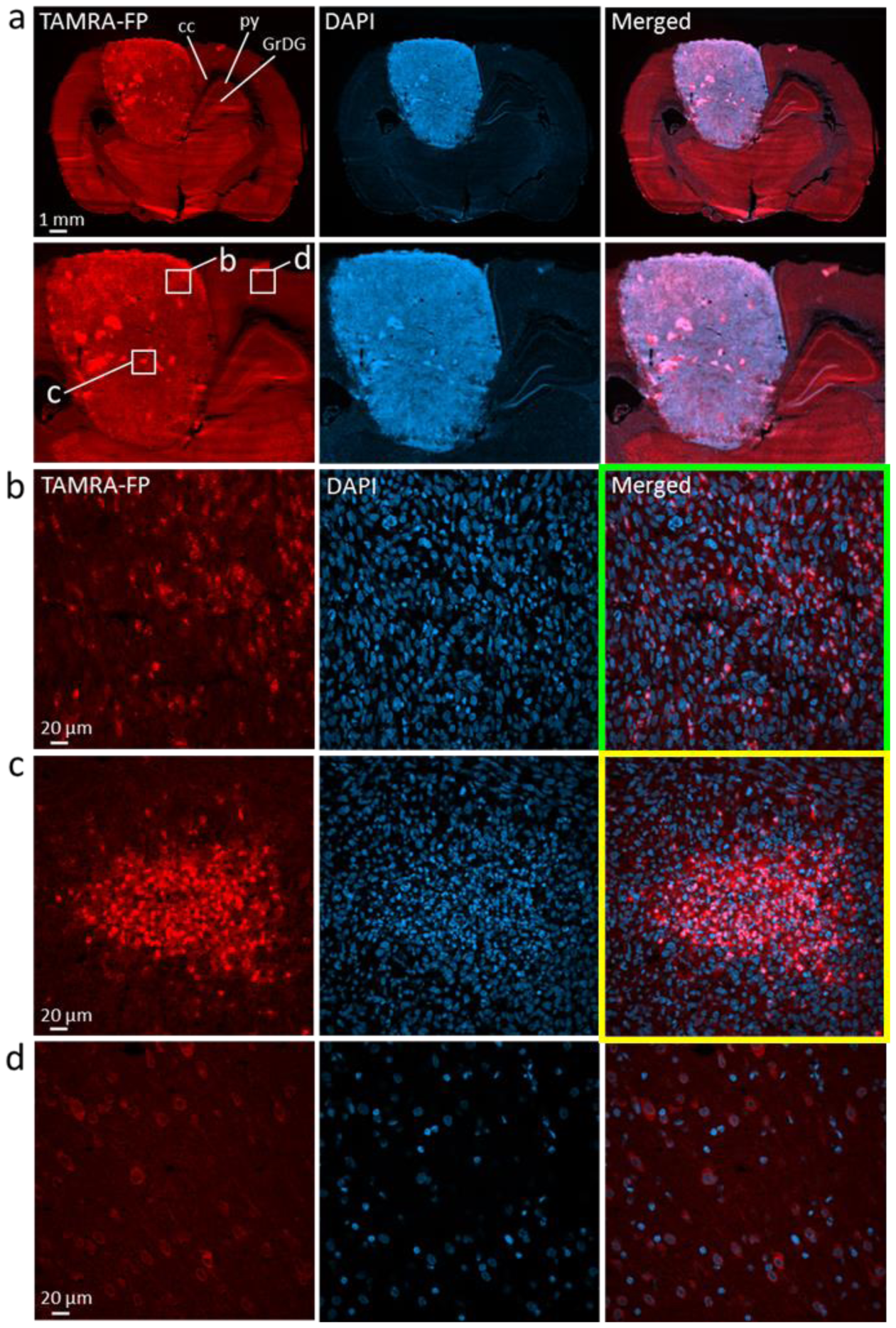
Distribution and overall characteristics of TAMRA-FP signal in glioma and control brain. Note relatively intense and heterogeneously distributed TAMRA-FP fluorescence over the glioma as compared to most regions of the healthy brain. Nuclear DAPI staining indicates dense cell population in glioma as compared to most regions of the healthy brain. Note relatively intense TAMRA-FP and DAPI signal also in cell-dense hippocampal pyramidal cell layer (py) and granular layer of dentate gyrus (GrDG) as well as relatively weak TAMRA-FP signal in white matter tracts of corpus callosum (cc) (**a**). Building on tissue-ABPP images of gliomas from different animals, a common pattern of TAMRA-FP labeling was established: **TAMRA-FP hotspots (b)**, characterized by intense and widely-distributed non-nuclear TAMRA-FP signal over the glioma, originating from evenly distributed individual cells. We define the second pattern as **TAMRA-FP hotspot clusters (c)**, characterized by intense non-nuclear TAMRA-FP signal originating from cell clusters. In healthy cortical region shown for comparison (**d**), TAMRA-FP signal is less intense and localizes mainly to cytosol and plasma membrane. The section illustrated here for TAMRA-FP and DAPI staining was further immunostained for the phagocyte marker CD11b/c and is presented again (Figure S18). 3D-animation of merged TAMRA-FP-DAPI fluorescence throughout the section thickness in TAMRA-FP hotspots (green lining) and TAMRA-FP hotspot clusters (yellow lining) is shown in Supplementary Video 1.

Building on tissue-ABPP images of gliomas from different animals, a common pattern of TAMRA-FP labeling emerged and the following classification will be used to facilitate signal interpretation in forthcoming immunohistochemistry. The most common TAMRA-FP labeling pattern with widest distribution over the glioma was characterized by non-nuclear intense fluorescence originating from evenly distributed individual cells (Figure 1b, Video S1). In what follows, we define these cells “TAMRA-FP hotspots”. However, the most intense TAMRA-FP labeling pattern was non-nuclear and originated from cell clusters with variable size (Figure 1c, Video S1). We define this labeling pattern “TAMRA-FP hotspot clusters”. In healthy cortex shown for comparison, TAMRA-FP fluorescence was less intense and mainly localized to cytosol and plasma membrane (Figure 1d).

### Probe labeling of tumor and healthy brain SHs is differentially sensitive to SH inhibitors

To assure that TAMRA-FP specifically reports SH activity under the conditions of tissue-ABPP, we used the competitive approach testing a panel of serine-nucleophile targeting broad-spectrum inhibitors, containing either FP (desthiobiotin-FP, MAFP, IDFP) or sulfonylfluoride (AEBSF and PMSF) as the warhead. Inhibitors were used at maximally effective concentrations to ensure comprehensive blockade of SHs activity prior to TAMRA-FP labeling. As evident from Figure 2, desthiobiotin-FP sharing the warhead-linker moiety with TAMRA-FP, effectively inhibited TAMRA-FP binding throughout the brain section, although weak residual labeling was evident as sparse spots over the glioma. Further, treatment with PMSF and IDFP efficiently prevented probe labeling in most brain regions, although some residual signal persisted over the tumor. Intriguingly, while the mSH-inhibitor MAFP efficiently blocked probe binding in most regions of the healthy brain, it was less effective in preventing TAMRA-FP labeling of the tumor, indicating that while MAFP-sensitive SH activity predominated in normal brain, SH activity originating from glioma was rather resistant to this inhibitor. This is a trailblazing finding, as MAFP acts as a pan-SH inhibitor potently targeting the vast majority of mSHs (Bachovchin et al., 2014).

**Figure 2.**
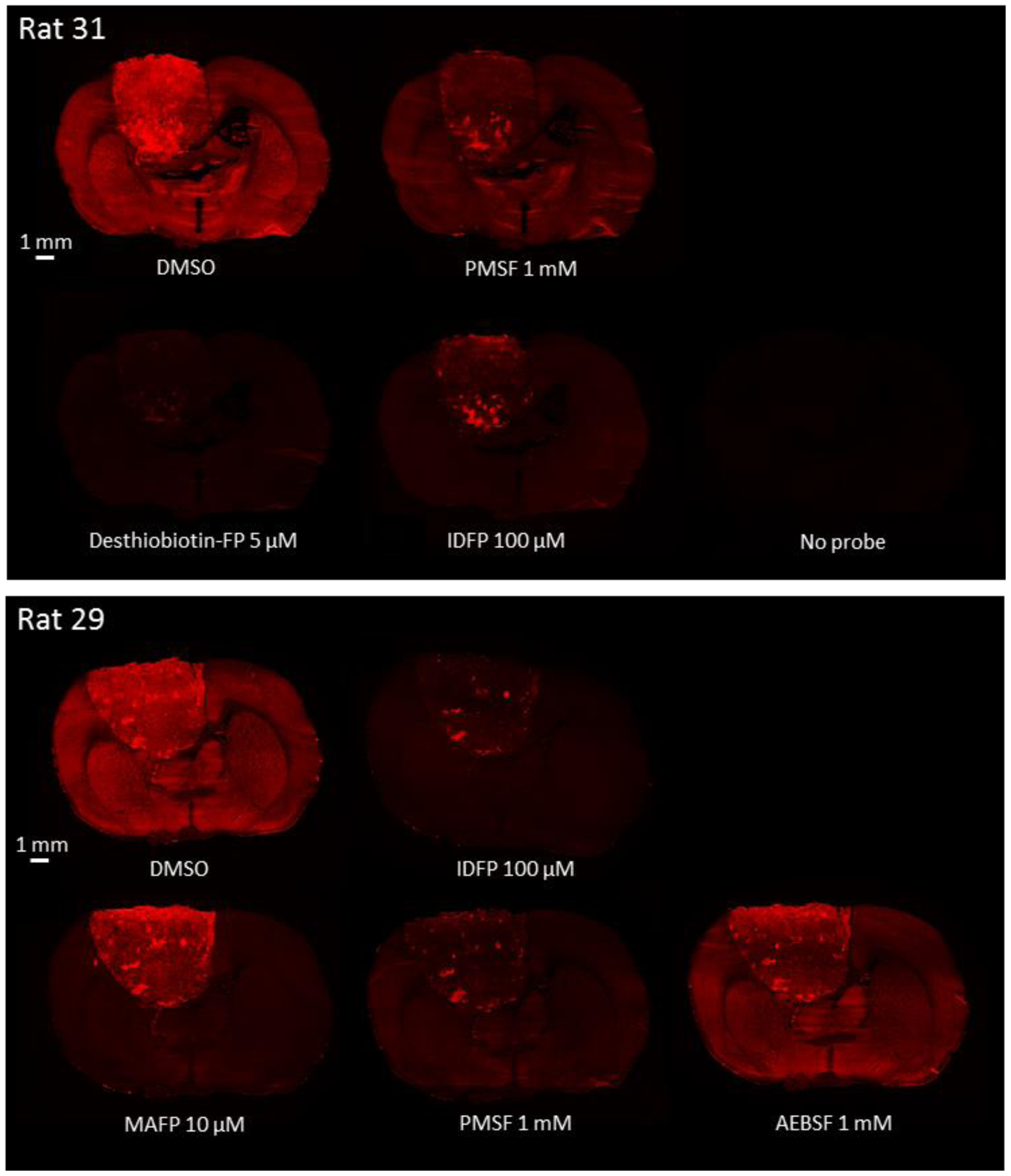
Competitive tissue-ABPP demonstrating that TAMRA-FP reports SH activity in glioma brain sections. Sections were pretreated for 1h with the indicated concentrations of various serine-nucleophile targeting inhibitors and processed for TAMRA-FP labeling and confocal fluorescence imaging as detailed in online Methods. Sections from two individual rats bearing the tumor were used and images from two independent experiments are shown as separate panels with DMSO control included in both experiment. Desthiobiotin-FP (5 µM) efficiently inhibits TAMRA-FP labeling throughout the brain section. However, a weak residual signal persists as spots over the glioma tissue. Note absence of fluorescence in the section processed without TAMRA-FP, indicating no detectable autofluorescence in the Cy3-window used for TAMRA-FP imaging. Note that PMSF (1 mM), in contrast to AEBSF (1 mM), and IDFP (100 µM) both efficiently inhibit TAMRA-FP labeling throughout the brain sections, yet leaving some residual activity over the tumor. Note in particular that MAFP (10 µM) efficiently inhibits probe labeling throughout the healthy brain regions but only marginally inhibits TAMRA-FP labeling of the tumor tissue. The scale bars represent 1 mm. Images were adjusted for brightness and contrast.

### Comparative gel-based ABPP of SH activity in homogenates of glioma and control brain

We cross-validated the tissue-ABPP findings in gel-based ABPP using homogenates of glioma and control brain. In addition, we used rat cerebellar membranes to facilitate comparison, as previous gel-based ABPP has frequently relied on rodent brain membrane proteomes. These studies revealed that while many of the SH bands were common to both proteomes (Figure 3), brain-resident SHs, including MAGL (∼35 kDa) and KIAA1363 (∼50 kDa) showed variable activity in glioma. Both were proposed to play protumorigenic role in various cancers (Nomura et al., 2010). While comparable activity was evident for KIAA1363, MAGL showed low-to-non-detectable activity in glioma, suggesting that it might not play a major role in this glioma model. In addition, two distinct mSHs, namely LYPLA1/2 (∼25 kDa) showed prominent, yet comparable activity in control brain and glioma. LYPLA1/2 are depalmitoylases regulating membrane-association and oncogenic signaling of Ras, and are inhibited by palmostatin B (Lin et al., 2017 and references therein).

**Figure 3.**
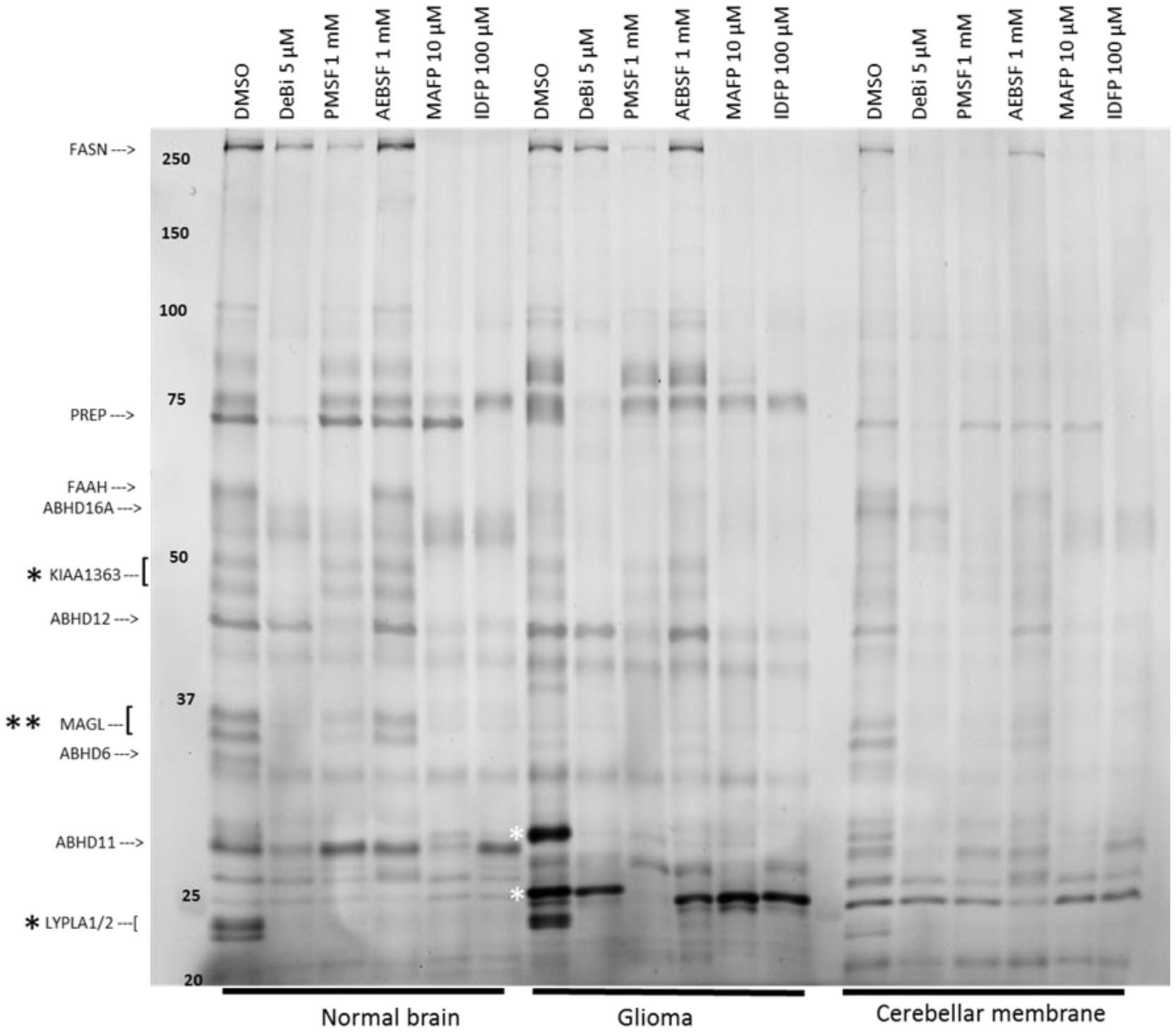
Comparative and competitive gel-based ABPP reveals distinct SH activity profiles of glioma and healthy brain. Rat cerebellar membranes were included as an additional control to facilitate SH band comparison and identification. Proteomes (1 mg/ml) were treated for 1h with DMSO or the indicated concentrations of the SH inhibitors, after which TAMRA-FP labelling was conducted for 1h as detailed in online Methods. The reaction was quenched and 10 µg protein was loaded per lane and separated by SDS-PAGE. TAMRA-FP labeled bands appear as black after in-gel imaging. Position of molecular weight markers (in kDa) are indicated on the gel. Based on previous studies from this and other laboratories, some healthy brain-resident SHs were identified, as indicated at left. Note comparable activities of KIAA1363 (also known as AADACL1 or NCEH1) and LYPLA1/2 doublets (black asterisk) in homogenates of control brain and glioma. Note also prominent activity of MAGL doublet (double black asterisk) in control brain as opposed to hardly detectable activity in glioma. Note that the glioma homogenate contains two prominent SH bands, migrating at ∼25 and ∼30 kDa (white asterisks). It is noteworthy that the respective SH bands are either absent or show low activity in control brain homogenate. Inhibitor profiling reveals that the glioma 30 kDa SH band is sensitive to all tested inhibitor whereas the ∼25 kDa band is fully sensitive to PMSF and less so to deshiobiotin-FP (DeBi), but resistant to other tested inhibitors, including MAFP. The gel is representative of two independent ABPP runs with similar outcome. ABHD1, ABHD6, ABHD11, ABHD12, ABHD16A, α,β-hydrolase domain-containing 1, 6, 11, 12 and 16A; FAAH, fatty acid amide hydrolase; FASN, fatty acid synthase; KIAA1363, also known as AADACL1 or NCEH1; LYPLA1/2, lysophospholipase A1/A2 (also known as acyl-protein thioesterases 1/2 or APT1/APT2); MAGL, monoacylglycerol lipase; PREP, prolyl oligopeptidase, also known as POP. Image was adjusted for brightness and contrast.

Of note, the glioma possessed two prominent SH bands migrating at ∼25 and ∼30 kDa (Figure 3). It is noteworthy that the respective SH bands showed low-to-non-detectable activity in control brain. We verified that the profile differences were present in homogenates from seven glioma rats (Figure S7). Inhibitor profiling revealed that the ∼30 kDa band was sensitive to all tested inhibitors whereas the ∼25 kDa band was sensitive to PMSF and deshiobiotin-FP, being resistant to other inhibitors, including MAFP. These findings not only cross-validate findings from tissue-ABPP but additionally indicate that the SH activity profile in glioma is distinct from that in normal brain and must therefore reflect activity originating from cell types not normally residing in brain.

### Tissue-ABPP combined with immunohistochemistry enables subcellular localization of SH activity within the tumor microenvironment

The glioma is a highly heterogeneous cancer and consists of several cell types, including tumor cells, endothelial cells from angiogenic vessels, neurons, scar-forming astroglia, as well as infiltrating monocytes/macrophages that are recruited from the periphery (Morisse et al., 2018; Quail and Joyce, 2017; Wirth et al., 2012). We approached this cellular complexity by immunohistochemistry using various cell-type markers that were applied on TAMRA-FP labeled sections.

To visualize tumor cells, sections were stained for Ki67, a nuclear marker of proliferating cell populations. Proliferating scattered cell populations were detected in glioma but as expected, were absent from healthy brain (Figure S8). Ki67-positive cells showed poor co-localization with TAMRA-FP hotspots or TAMRA-FP hotspot clusters.

GFAP-positive astrocytes appeared as a dense cell population encircling the tumor (Figure S9). Although scattered GFAP-positive cells were detected within the tumor, the bulk of tumor was devoid of astrocytes, in line with previous findings with this model (Wirth et al., 2012). The sparse presence of astrocytes is consistent with the gliosarcoma character of this model (Barth and Kaur, 2009). Although astrocytes showed moderate SH activity, TAMRA-FP hotspots or TAMRA-FP hotspot clusters showed poor co-localization with GFAP-positive cells. Throughout the healthy brain, cells with star-shaped morphology (i.e. astrocytes) were visible.

Tumor vasculature, visualized using the endothelial markers von Willebrand factor and CD34, revealed relatively low TAMRA-FP signal originating from endothelial cells (Figure S10). However, TAMRA-FP hotspot clusters resided often in the vicinity of the vessels. This glioma model shows leaky blood vessels with compromised blood-brain-barrier (BBB) function, especially at the tumor edge (Figure S1). Sections were stained for the redox-sensitive marker heme oxygenase-1 (HO-1) to visualize areas prone to redox stress (Figure S11). A heterogeneous pattern of HO-1 staining was evident in the tumor with no detectable signal in control brain. However, although HO-1 positive cells localized to regions of intense SH activity, neither TAMRA-FP hotspots nor TAMRA-FP hotspot clusters stained strongly for HO-1.

In the absence of truly selective microglial marker, we stained sections for Iba1 (Figure S12), being aware of that this marker also labels activated monocytes and macrophages. We observed heterogeneous presence of Iba1-positive cells throughout the tumor with intense staining in tumor margins, and noted partial overlap of Iba1 staining with TAMRA-FP hotspots. However, no Iba1-positive cells were evident in tumor regions of TAMRA-FP hotspot clusters. As expected, Iba1-positive cells with characteristic microglial morphology were detected in the healthy brain (Figure S12).

### SH activity in relation to hyaluronan (HA) – CD44 axis and markers of tumor stiffness

Stromal HA accumulation is considered as a protumorigenic factor in several solid tumors (McCarthy et al., 2018; Tammi et al., 2019). Extensive modeling of the extracellular matrix associated with biomechanical changes, often called tumor “stiffness”, is a hallmark of “cancerized” fibrotic stroma (McCarthy et al., 2018). The HA receptor CD44 promotes cancer cell motility, tumor growth, angiogenesis as well as resistance to chemo- and radiotherapy, and is overexpressed in various tumors, including GBM (Mooney et al., 2016). We clarified whether TAMRA-FP hotspots localize to tumor regions undergoing matrix remodeling using the stiffness markers pMLC2 and tenascin C along with HA and CD44. HA was detected in the extracellular matrix throughout the brain with more intense staining in glioma, especially the glioma edges (Figure S13). HA was abundant in areas of TAMRA-FP hotspots. In contrast, HA was sparse in regions of TAMRA-FP hotspot clusters, whereas these clusters were surrounded by HA-enriched tissue. Like HA, CD44 was enriched in the glioma with low expression in normal brain (Figure S14). Like HA, CD44 was abundant in glioma regions of TAMRA-FP hotspots but was undetectable in regions of TAMRA-FP hotspot clusters, yet these clusters were surrounded by CD44-positive cells. pMLC2 showed intense staining over the glioma but was absent from healthy brain (Figure S15). Interestingly, the stiffness marker frequently co-localized with TAMRA-FP hotspots. In contrast, pMLC2-positive cells were rare in the central part of TAMRA-FP hotspot clusters whereas TAMRA-FP-positive cells at the edge of these clusters expressed pMLC2 and were also surrounded by pMLC2-positive cells. The outcome with tenascin C (Figure S16) was similar to what was observed with pMLC2.

### SH activity in relation to cells of myeloid and lymphoid origin

Bone-marrow-derived cell populations are abundant in tumors, including GBM (Morisse et al., 2018; Quail and Joyce, 2017). In human and rodent gliomas, tumor-associated macrophages (TAMs) are the major class of tumor-promoting immune cells and were also previously detected in this model (Wirth et al., 2012). We pursued to identify TAMs and other bone-marrow-derived cells in glioma. Interestingly, both TAMRA-FP hotspots and hot spot clusters expressed the hematopoietic marker CD45 (Figure S17) or the phagocyte marker CD11b/c (Figure S18). Although macrophages were abundant in glioma regions showing prominent SH activity, neither TAMRA-FP hotspots nor TAMRA-FP hotspot clusters stained for the macrophage markers CD68 (Figure S19), CD163 (Figure S20) or CD169 (Figure S21). Lymphoid cells were present in glioma regions showing prominent SH activity and TAMRA-FP hotspots partially overlapped with CD4- and CD8-positive cells, whereas TAMRA-FP hotspot clusters sparsely expressed the T-cell markers (Figures S22 and S23).

### Tumor-associated mast cells partly account for the high SH activity

We found low expression of the mast cell marker FcεRIγ within TAMRA-FP hotspots whereas FcεRIγ-positive cells were enriched in TAMRA-FP hotspot clusters, showing marked co-localization with TAMRA-FP (Figure S24). Besides mast cells, FcεRIγ labels eosinophils, basophils and monocytes, so this marker alone cannot verify the presence of mast cells. Mast cell-specific serine proteases include tryptases and chymases. We stained the sections for chymase using CMA1 antibody. Faint expression of CMA1 was evident in glioma where CMA1-positive cells co-localized with TAMRA-FP hotspots, whereas CMA1-positive cells were not detected in TAMRA-FP hotspot clusters (Figure S25). Unfortunately, we found no additional antibodies that qualified for further verification of mast cell presence. Further, attempts to detect mast cells by enzymo-histochemical staining of serine protease activity using peptide substrates tailored for human tryptase and chymase (Harvima et al., 1990) were unsuccessful (data not shown).

### Tumor-associated neutrophils (TANs) account for tumor SH activity hotspots

Neutrophils encompass four serine proteases (NSPs), namely neutrophil elastase (NE), cathepsin G (CATG), proteinase 3 (PR3), and neutrophil proteinase 4 (NSP4, also known as PRSS57) (Benarafa and Simon, 2017; Kasperkiewicz et al., 2017). We stained glioma sections for the neutrophil marker myeloperoxidase (MPO) and found that MPO-positive cells localized to TAMRA-FP hotspots (Figure S26). Further, MPO-positive cells were abundant in TAMRA-FP hotspot clusters (Figure S26). The outcome was essentially the same in sections stained for elastase (Figure S27). Further, high-resolution imaging revealed that TAMRA-FP hotspots had the morphological characteristics of neutrophil-type multi-nucleated cells and that TAMRA-FP labeling of these cells was sensitive to PMSF and desthiobiotin-FP (Figure 4). In sham-operated animals, TAMRA-FP hotspots with TAN-like morphology were rare at the site of injection (Figure S28), indicating that the surgical operation *per se* did not cause TAN accumulation. Collectively these findings indicated that TANs are the principal cell-type accounting for TAMRA-FP hotspots and hotspot clusters.

**Figure 4.**
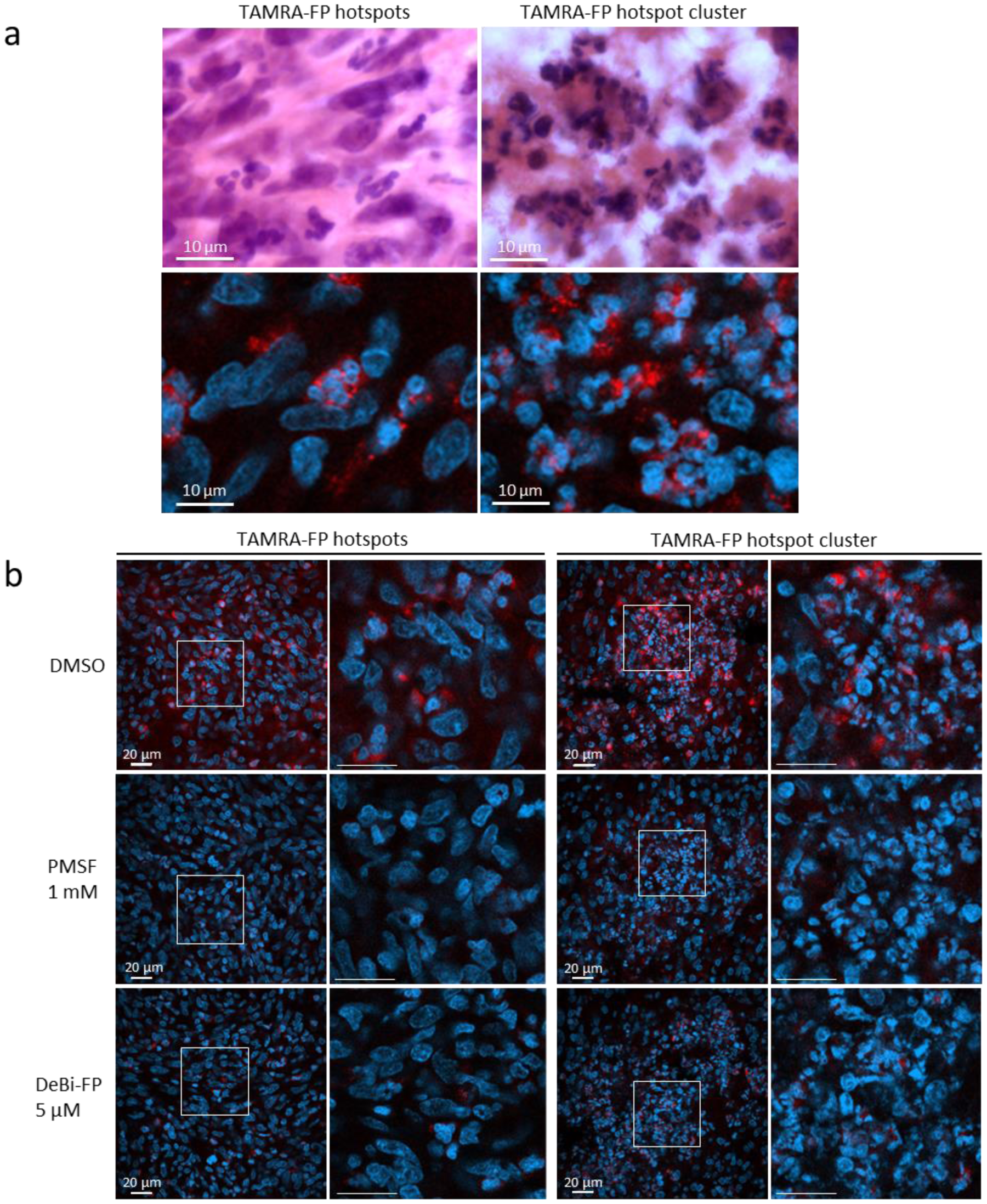
High-resolution imaging of TAMRA-FP hotspots and TAMRA-FP hotspot clusters and their inhibitor sensitivity in glioma. In **a**, the upper panel shows hematoxin-eosin (H&E) stained section. For images in the lower panel, sections went through the tissue-ABPP protocol to label SHs (red), followed by DAPI staining to visualize nuclei (blue). Note presence of multi-nucleated cells with TAN-like morphology in regions of TAMRA-FP hotspots and TAMRA-FP hotspot clusters. In **b**, sections were pretreated with DMSO or with the SH inhibitors PMSF (1 mM) or deshiobiotin-FP (DeBi-FP, 5 µM) for 1h at RT, after which they went through the tissue-ABPP protocol to label SHs (red), followed by DAPI staining to visualize nuclei (blue). Note that throughout the examined regions, TAMRA-FP labeling is sensitive to the inhibitors. Scale bars 10 µm (**a**) and 20 µm (**b**). Images were adjusted for brightness and contrast.

### ABPP of glioma proteome using novel activity probes designed for human serine proteases

In further efforts to characterize glioma serine protease activity, we utilized newly introduced activity probes with preference for trypsin and elastase (Edgington-Mitchell et al., 2017) using both gel- and tissue-based ABPP approaches. These probes bear a diphenylphosphonate (DPP) warhead, which targets serine proteases more specifically than the FP warhead which broadly targets the mSH family (Bachovchin et al., 2010). Besides glioma, we profiled tumor cells and bone-marrow-derived mononuclear cells (rBM). Interestingly, the elastase-preferring probe V-DPP detected 3-4 proteins (∼25 kDa) in rBM and weakly two bands of similar size in glioma (Figure S29). Probe binding was sensitive to SH inhibitors, indicating that V-DPP specifically reported elastase-type serine protease activity. The trypsin-preferring probe PK-DPP gave no detectable signal in gel-ABPP (Figure S29). When tested in tissue-ABPP in place of TAMRA-FP, probe binding was hardly visible and neither probe labeled inhibitor-sensitive targets in glioma (Figure S30).

We evaluated also novel activity probes bearing the DPP warhead that were tailored to report individually activities of the four human NSPs (Kasperkiewicz et al., 2017). Unfortunately, the NSP-probes did not recognize any TAMRA-FP sensitive bands in gel-ABPP of rat proteomes (Figure S31). When tested in tissue-ABPP in place of TAMRA-FP, the NSP-probes readily bound to sections, yet in non-specific manner, as probe binding was not prevented by TAMRA-FP (Figure S31).

As a final step, we used gel-ABPP to assess inhibitor sensitivity of TAMRA-FP labeled proteins in neutrophils and rBMs (Figure S32). Only few prominent bands migrating at ∼25-30 kDa were present in these proteomes and TAMRA-FP labeling of these bands was inhibited by PMSF, desthiobiotin-FP and AEBSF, but was largely resistant to MAFP and IDFP, consistent with the pharmacology of the ∼25 kDa glioma band. Compound 22, designed to potently target human CTSG (Swedberg et al., 2017), was inactive in the rat proteomes whereas another CTSG inhibitor (GTSG-I) blocked TAMRA-FP labeling of the ∼25 kDa band in both samples. In tissue-ABPP, the inhibitor did not prevent TAMRA-FP labeling of hotspots or hotspot clusters (Figure S32).

### LC-MS/MS analysis of the glioma SH bands migrating at ∼25-30 kDa

Gel-pieces encompassing the prominent SH bands in glioma, rBMs and tumor cells were cut after in-gel fluorescence scanning and subjected to LC-MS/MS-based target identification. To facilitate SDS-PAGE separation of proteins with similar size, proteomes were deglycosylated prior to TAMRA-FP labeling. The complete LC-MS/MS data with all identified proteins is available in Supplementary File 1. Interestingly, neutrophil serine proteases GTSG, NE and PR3 were found from the ∼25 kDa gel-pieces of glioma and bone marrow samples (Figure S33), confirming findings from tissue-ABPP. Interestingly, the ∼25-30 kDa gel-pieces encompassed also two MAFP-sensitive SHs (Bachovchin et al., 2014), namely PAF-acetylhydrolases 1b2 and 1b3 (PAFAH1b3 and PAFAH1b2).

## Discussion

Our study widens the applicability of the chemoproteomic technology to tissue sections, enabling for the first time high-resolution confocal fluorescence imaging of SH activity in anatomically preserved complex native cellular environment. We named this technique tissue-ABPP to distinguish it from conventional gel-based ABPP. Thorough optimization and validation was provided by parallel gel-based ABPP, extensive immunohistochemical mapping of TAMRA-FP signal origin, as well as LC-MS/MS-based target verification. Collectively all the evidence demonstrates convincingly that tissue-ABPP faithfully reports SH activity in tissue sections at subcellular resolution. We emphasize that it is TAMRA-FP labeling combined with immunohistochemistry and DAPI staining, each step utilizing different fluorophores with minimal wavelength overlap that makes tissue-ABPP a remarkably powerful new approach to explore SH activity.

The focus here was to achieve high-resolution imaging of SH activity throughout the glioma microenvironment, necessitating systematic immunohistochemical mapping of origins of SH activity and its localization with cell-type and tissue stiffness markers (Figures S8-28). Together with inhibitor-sensitivity of TAMRA-FP labeling, these studies provide the key validation for tissue-ABPP, ruling out the possibility that TAMRA-FP labeling is nonspecific. Besides illuminating the complexity of tumor microenvironment, our study unveiled heightened SH activity in glioma vs. normal brain and excluded major tumor-associated cells-types such as TAMs as a source of this activity. Unexpectedly, we pinpointed TANs as the principal cell-type generating SH activity hotspots in glioma.

The activity probe that we used is commercially available and similar FP-probes have been previously used and extensively characterized for gel-based ABPP applications (Niphakis and Cravatt, 2014; Nomura et al., 2010; Simon and Cravatt, 2010). Furthermore, the coverage of mSH-family members recognized by the FP-probes is notable (>80 %) (Bachovchin et al., 2010). SHs are key regulators of metabolic pathways in diseases including cancer, neurodegenerative diseases, metabolic syndrome and atherosclerosis. Tissue-ABPP offers a new tool to explore the myriad roles of SHs likely play in physiology and disease. We anticipate that tissue-ABPP should find broad immediate applications. We applied tissue-ABPP also successfully to spleen sections, mainly to provide additional validation in native immune tissue for TAMRA-FP labeling in relation to some of the antibodies (Figures S34 and S35). Previously, a single study has presented low-resolution TAMRA-FP imaging of AEBSF-sensitive SH activity in prostate cancer specimen in supplementary figure (Dudani et al., 2018). Although preliminary, that work extends the applicability of tissue-ABPP to human proteomes. As ABPP requires no *a priori* knowledge of the identity of the target, we envisage no imaginable reason why tissue-ABPP would not work for sections regardless of species and tissue source.

Our study did not definitely reveal identity of the individual SHs generating TAMRA-FP hotspots or hotspot clusters in glioma. However, both cell morphology and immunohistochemical evidence revealing close localization of neutrophil markers to TAMRA-FP hotspots, together with LC-MS/MS-based verification of NSPs in the gel-pieces encompassing the ∼25 kDa TAMRA-FP bands, allowed to conclude that NSPs account for the bulk of this activity. In further support are findings showing that the elastase-preferring V-DPP-probe detected inhibitor-sensitive ∼25 kDa bands in glioma and that neutrophil and glioma proteomes shared similarly migrating intense SH bands showing similar pharmacology. Collectively these findings indicate that NSPs rather than mSHs accounted for the TAMRA-FP hots spots and hotspot clusters.

Interestingly, the ∼30 kDa SH was enriched in glioma and was not detected in healthy brain or neutrophils. LC-MS/MS disclosed PAFAH1b3 and PAFAH1b2, among others, in the gel-pieces encompassing the ∼25-30 kDa TAMRA-FP bands. Noteworthy, these mSHs are MAFP-sensitive (Bachovchin et al., 2014) and were proposed to play protumorigenic role in various cancers (Chang et al., 2015; Kohnz et al., 2015; Mulvihill et al., 2014). Future work should elucidate the role of PAFAH1b3 and PAFAH1b2 in glioma.

The newly developed activity probes (Kasperkiewicz et al., 2017) and CTSG inhibitor Compound 22 (Swedberg et al., 2017) were designed to target human NSPs. In our study, the compounds behaved as expected when tested using human proteomes (Figure S31 and data not shown). However, our cross-validation experiments clearly indicated that the Cy5-probes did not recognize mutual targets with TAMRA-FP in neutrophils or glioma. This was not totally unexpected, as both probe and inhibitor design exploited unique substrate recognition sequences around the catalytic sites which are not necessarily shared between human and rat NSPs. It is likely that the different warhead (DPP vs. FP) could also account for major differences in the reactivity and sensitivity of these probes to the inhibitors.

We recognize that rat gliosarcoma model is not widely used. Nevertheless, it has been useful to test novel chemotherapeutic targeting strategies, antitumor effects of gene therapy, anti-angiogenic agents alone or in combination with radiation and chemotherapy (Barth and Kaur, 2009 and references therein). We chose this model for convenience as it was previously used by researchers at the UEF (Wirth et al., 2012) and the expertise to execute such studies was readily available.

Our study did not address the role(s) that TANs might play in glioma. TANs are newcomers at the mainstage of cancer research (Shaul and Fridlender, 2018). In GBM, TANs seem to play mainly a tumorigenic role (Massara et al., 2017). A recent study suggested involvement of TANs in GBM recurrence after radiotherapy by showing that neutrophils promoted cancer cells stemness (Jeon et al., 2019). In clinical samples, a positive correlation existed between neutrophils and patients diagnosed with recurrent tumor (Jeon et al., 2019). Future work could elucidate the role of TANs and NSPs in more detail.

We did not provide detailed characterization of SH activity in different regions or cell types of healthy brain. Gel- and MS-based ABPP has shown that isolated mouse brain primary cells such as neurons, astrocytes and microglia each show characteristic SH activity profiles (Viader et al., 2016). Tissue-ABPP offers a powerful tool for detailed investigation of this issue in situ with all normal cell types present. Such studies should be of wide interest to increase knowledge on the myriad roles that SHs likely play in the CNS. A straightforward starting point could be comparative tissue-ABPP of an individual SH, e.g. the principal endocannabinoid hydrolase MAGL using brain sections of wild-type and MAGL-deficient mice, as MAGL knockout does not lead to compensatory changes in the activity of other SHs, as evidenced by gel-based ABPP (Navia-Paldanius et al., 2015). Competitive tissue-ABPP would offer a complementary approach for sections where MAGL is chemically inactivated with highly selective and ultra-potent inhibitors, such as JJKK-048 and KML29 (Aaltonen et al., 2013; Bachovchin and Cravatt, 2012). We envision that in addition to TAMRA-FP, tissue-ABPP could possibly utilize newly-developed activity probes tailored to target only a subset of mSHs (van Rooden et al., 2018).

## Supporting information

Supplementary Video related to Fig. 1

## Acknowledgements

J.T.L. was supported by the Finnish Academy (grant no. 278212). L.E.M. was supported by a Grimwade Fellowship from the Russell and Mab Grimwade Miegunyah Fund at The University of Melbourne and a DECRA Fellowship from the Australian Research Council (ARC, DE180100418). Drag laboratory is supported by Foundation for Polish Science and National Science Centre in Poland. PK. is the beneficiary of a L’Oreal Poland and the Polish’ Ministry of Science and Higher Education scholarships. We thank Tanja Kosunen and Dina Navia-Paldanius for contributing to the pilot tissue-ABPP experiments and Miika Martikainen for practical guidance with glioma animals. We thank Mikko Kettunen and Kimmo Jokivarsi for help in MRI and Taina Vihavainen, Taija Hukkanen, Satu Marttila, Eija Rahunen, Shabana Azmy, Juha Niskanen for skillful technical help. We thank Anne Koivisto and Ilkka Harvima for enzymo-histochemical staining of tryptase and chymase. We thank Sini Miettinen and Tiina Öhman for help in analyzing and processing the LC-MS/MS data. We thank Joakim Swedberg (at that time in the Institute for Molecular Bioscience, University of Queensland, Brisbane, Australia) for providing compound 22. We are grateful for the UEF Cell and Tissue Imaging Unit and for Kuopio Biomedical Imaging Unit (Kuopio-BIU) for providing facilities for histology, confocal microscopy and MRI.

## Author Contributions

J.T.L., T.W. and J.R.S. conceptualized the study. N.A., H.J., P.K.S., E.M. and S.K. carried out the in vitro experiments. N.A., P.K.S., T.W. and H.S. generated the glioma rats. P.K.S. and N.A. carried out MRI imaging of tumors, and harvesting of tissue from the animals. P.K.S. isolated bone-marrow cells and neutrophils, and performed cell culture and gel-based ABPP experiments. P.K.S. prepared samples and M.V. supervised the mass-spectrometry experiments. P.K.S. and J.R.S. analyzed the MS data. L.E.M. synthesized tryptase and elastase probes, P.K. and M.D. synthesized NSP probes. N.A. and H.J. carried out tissue-ABPP experiments. N.A., H.J., J.R.S. and J.T.L. designed tissue-ABPP experiments; N.A. and H.J. performed confocal fluorescence imaging with help and expertise provided by S.P-S. and K.R.. J.T.L., N.A. and P.K.S. wrote the manuscript. All others edited the manuscript. J.T.L. acquired the funding.

## Competing interests

The authors declare no competing interests.

## Materials and Methods

Detailed information for all Materials and Methods can be found in the Supporting information.

## Supporting information

The following supplemental files accompany this manuscript: Supporting information: Supplementary Figures S1-S35 and Materials and Methods; Supplementary Video 1: 3D-animation of merged TAMRA-FP-DAPI fluorescence throughout the section thickness in TAMRA-FP hotspots and TAMRA-FP hotspot clusters (related to Figure 1); Supplementary File 1_LC-MS-MS data: Complete LC-MS/MS data of all proteins identified from the ABPP gel-pieces.

## Materials and Data Availability

Correspondence and request for materials should be addressed to J.T.L. or J.R.S., to P.K and M.D. (NSP probes) or to L.E.M. (elastase/tryptase probes).

