## Supplementary Figures & Materials and Methods for "Tissue-ABPP enables high-resolution confocal fluorescence imaging of serine hydrolase activity in cryosections – Application to glioma brain unveils activity hotspots originating from tumor-associated neutrophils"

### Inventory

#### Supplementary Figures

Figure S1. Activity-based protein profiling (ABPP) and the power of this approach to unveil SH activity in glioma.

Figure S2. Characteristics of the rat BT4C gliosarcoma model.

Figure S3. Coronal plane MRI images of glioma and control brains.

Figure S4. Testing various fixation method for rodent brain sections.

Figure S5. Effect of TAMRA-FP concentration on fluorescence signal in mouse brain sections.

Figure S6. Effects of pH and buffer composition on TAMRA-FP signal and its inhibitor sensitivity.

Figure S7. Comparative gel-based ABPP of seven animals confirming distinct SH activity profiles between glioma and control brain.

Figure S8. Confocal imaging of SH activity in relation to proliferating tumor cells.

Figure S9. Confocal imaging of SH activity in relation to astrocytes.

Figure S10. Confocal imaging of SH activity in relation to blood vessels.

Figure S11. Confocal imaging of SH activity in relation to heme oxygenase 1 (HO-1).

Figure S12. Confocal imaging of SH activity in relation to microglial marker Iba1.

Figure S13. Confocal imaging of SH activity in relation to hyaluronan (HA).

Figure S14. Confocal imaging of SH activity in relation to HA receptor CD44.

Figure S15. Confocal imaging of SH activity in relation to the stiffness marker pMLC2.

Figure S16. Confocal imaging of SH activity in relation to the stiffness marker tenascin C.

Figure S17. Confocal imaging of SH activity in relation to CD45, a marker for nucleated hematopoietic cells.

Figure S18. Confocal imaging of SH activity in relation to CD11b/c, a marker for phagocytes.

Figure S19. Confocal imaging of SH activity in relation to CD68, a marker for monocytes and macrophages.

Figure S20. Confocal imaging of SH activity in relation to CD163, a marker for monocytes and macrophages.

Figure S21. Confocal imaging of SH activity in relation to CD169, a marker for macrophages.

Figure S22. Confocal imaging of SH activity in relation to T cell marker CD4.

Figure S23. Confocal imaging of SH activity in relation to T cell marker CD8.

Figure S24. Confocal imaging of SH activity in relation to FcεRIγ, a marker for mast cells, eosinophils, basophils and monocytes.

Figure S25. Confocal imaging of SH activity in relation to chymase (CMA1), a marker for mast cells.

Figure S26. Confocal imaging of SH activity in relation to myeloperoxidase (MPO), a marker for neutrophils.

Figure S27. Confocal imaging of SH activity in relation to neutrophil elastase (NE).

Figure S28. TAMRA-FP signal at the site of injection in sham-operated animals.

Figure S29. Gel-ABPP of rat glioma proteomes using Cy5-labeled serine protease activity probes PK-DPP and V-DPP.

Figure S30. Tissue-ABPP of glioma sections using Cy5-labeled activity probes PK-DPP and V-DPP.

Figure S31. ABPP of rat neutrophil and glioma samples using Cy5-labeled neutrophil serine protease (NSP) probes in combination with TAMRA-FP.

Figure S32. Inhibitor profiles of human cathepsin G (hCTSG) and the prominent 25-30 kDa SH bands in rat bone-marrow-derived mononuclear cells and neutrophils.

Figure S33. LC-MS/MS-based identification of the ~25-30 kDa SH bands in rat glioma and bone marrow-derived mononuclear cells.

Figure S34. High-resolution imaging of TAMRA-FP hotspots and their inhibitor sensitivity in rat spleen.

Figure S35. Confocal imaging of SH activity in rat spleen sections in relation to selected immunomarkers.

**Materials and Methods**

**Ethical issues**

**Chemicals and Reagents**

**Cell culture**

**Animals**

*Malignant rat glioma model*

*Tissue harvesting from glioma animals*

*Control rats used for harvesting immune cells and tissue*

*Mice used in preliminary tissue-ABPP experiments*

**MRI**

**Cryosectioning**

**Tissue-ABPP**

*Protocol for imaging SH activity*

*Tissue-ABPP combined with immunohistochemistry and nuclear staining*

**Histology**

**Fluorescence imaging**

*Fluorescence scanning*

*Confocal microscopy*

**Rat Bone Marrow Cell Isolation**

**Human and rat blood neutrophil isolation**

**Preparation of proteomes for gel-based ABPP**

**Competitive gel-based ABPP**

**Sample preparation for LC-MS/MS**

**LC-MS/MS analysis**

**References for the Materials and Methods section**

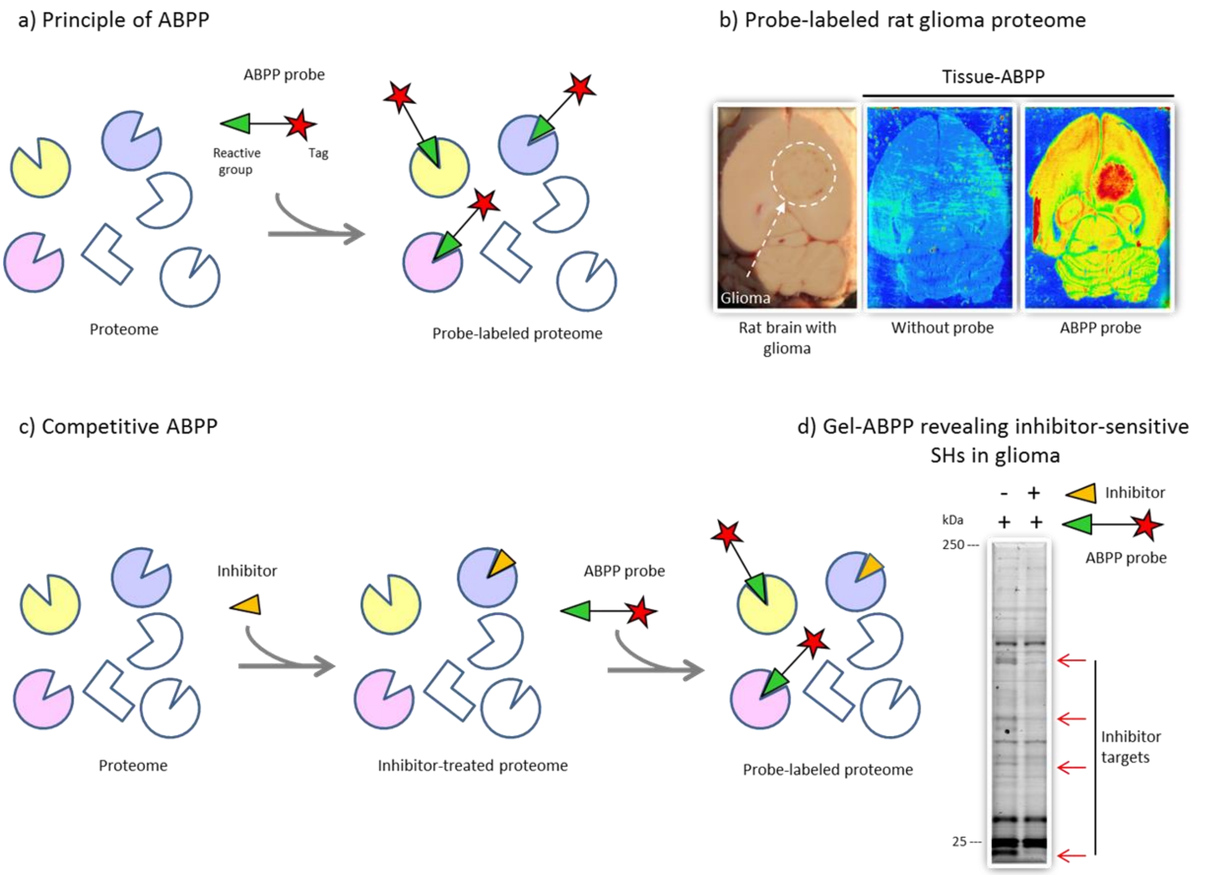


**Figure S1. Activity-based protein profiling (ABPP) and the power of this approach to unveil SH activity in glioma.** **a)** An active site-targeted covalent probe comprises a reactive fluorophosphonate (FP) group (warhead) that is coupled via a linker to a reporter tag (e.g. fluorescent dye or biotin) to covalently label, detect, enrich and identify active SHs (colored objects in this scheme). **b)** Tissue-ABPP unveils heightened SH activity in rat glioma. As compared to healthy brain tissue, the tumor lights up in red due to high SH activity, evident in the brain section treated with the ABPP probe TAMRA-FP (panel at right). Due to leaky blood vessels, the glioma borders appear as a brownish halo, as illustrated for the brain used in cryosectioning (panel at left). In competitive ABPP (**c**), the proteome is first treated with an inhibitor that covalently binds to the catalytic serine, inactivating the target enzyme. Thereafter, the proteome is reacted with the activity probe. Probe labeling of the inhibitor-targeted enzyme is prevented (a single target is illustrated here). **d)** Competitive gel-based ABPP revealing molecular size, relative abundance and inhibitor-sensitivity of the glioma-enriched SHs. The proteins are resolved using SDS-PAGE and visualized after in-gel fluorescence scanning. Treatment with a broadly-acting lipase inhibitor prevents probe labeling of several SH bands (red arrows). Images in panels A and C are modified from Bachovchin and Cravatt (2012).

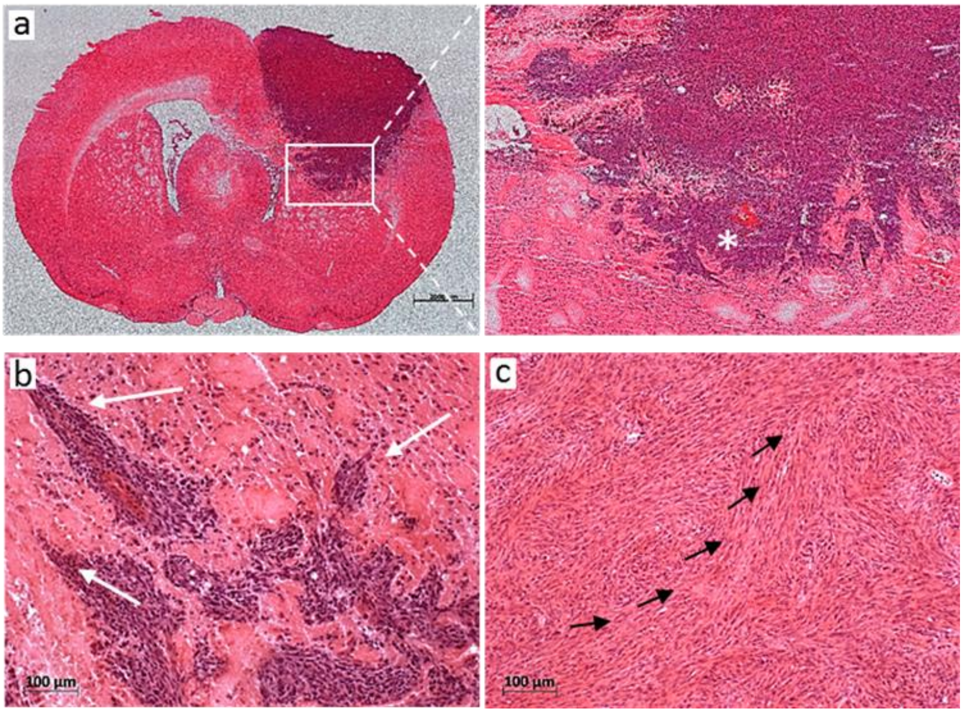


**Figure S2. Characteristics of the rat BT4C gliosarcoma model.** Tumors were generated as detailed in Materials and Methods and their presence verified by MRI imaging in vivo 22-24 days after implantation of tumor cells (Figure S3). Animals were sacrificed 5-11 days after MRI imaging, i.e. 4-5 weeks after implantation of tumor cells. 20 µm thick coronal brain cryosections were fixed with 4% PFA supplemented with 0.01% glutaraldehyde and stained with hematoxylin-eosin (H&E). The tumor grows expansively, invading the surrounding normal brain. Microhemorrhages are seen in tumor periphery (white asterisk) (**a**). White arrows point leading edges of migrating tumor cells (**b**). In tumor bed, H&E staining reveals sarcomatous pattern of glioma growth (black arrows) with elongated, spindle shaped cells (**c**). Scale bars 2 mm (a) and 100 µm (b and c).

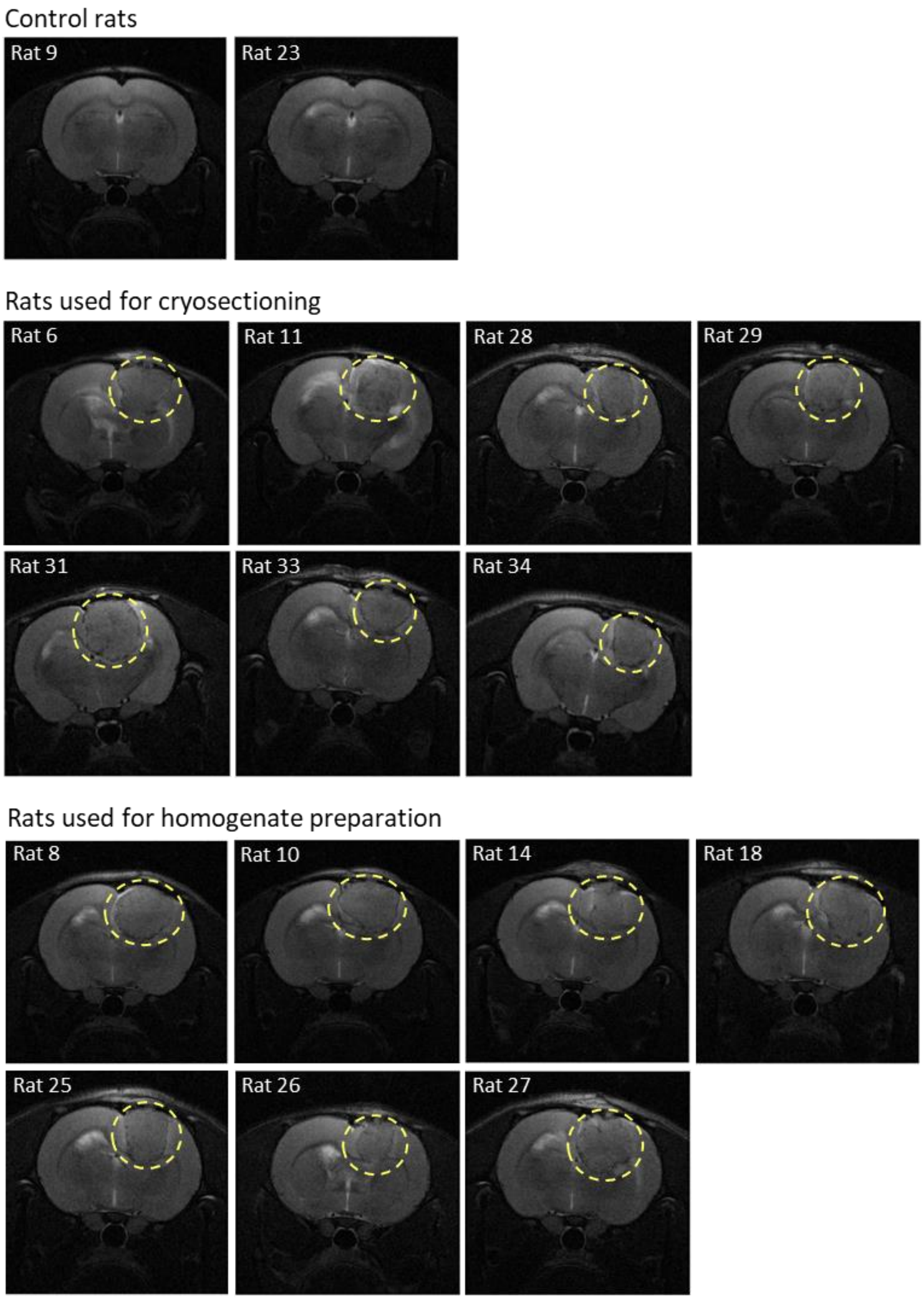


**Figure S3. Coronal plane MRI images of glioma and control brains.** Brains were used for cryosectioning (tissue-ABPP) or for preparation of homogenates (gel-based ABPP for validation experiments). Rats 9 and 23 were injected with the tumor cell medium only and served as controls to provide sections from the site of tumor implantation. Gliomas are visible inside the dashed circles. Images were adjusted for brightness and contrast.

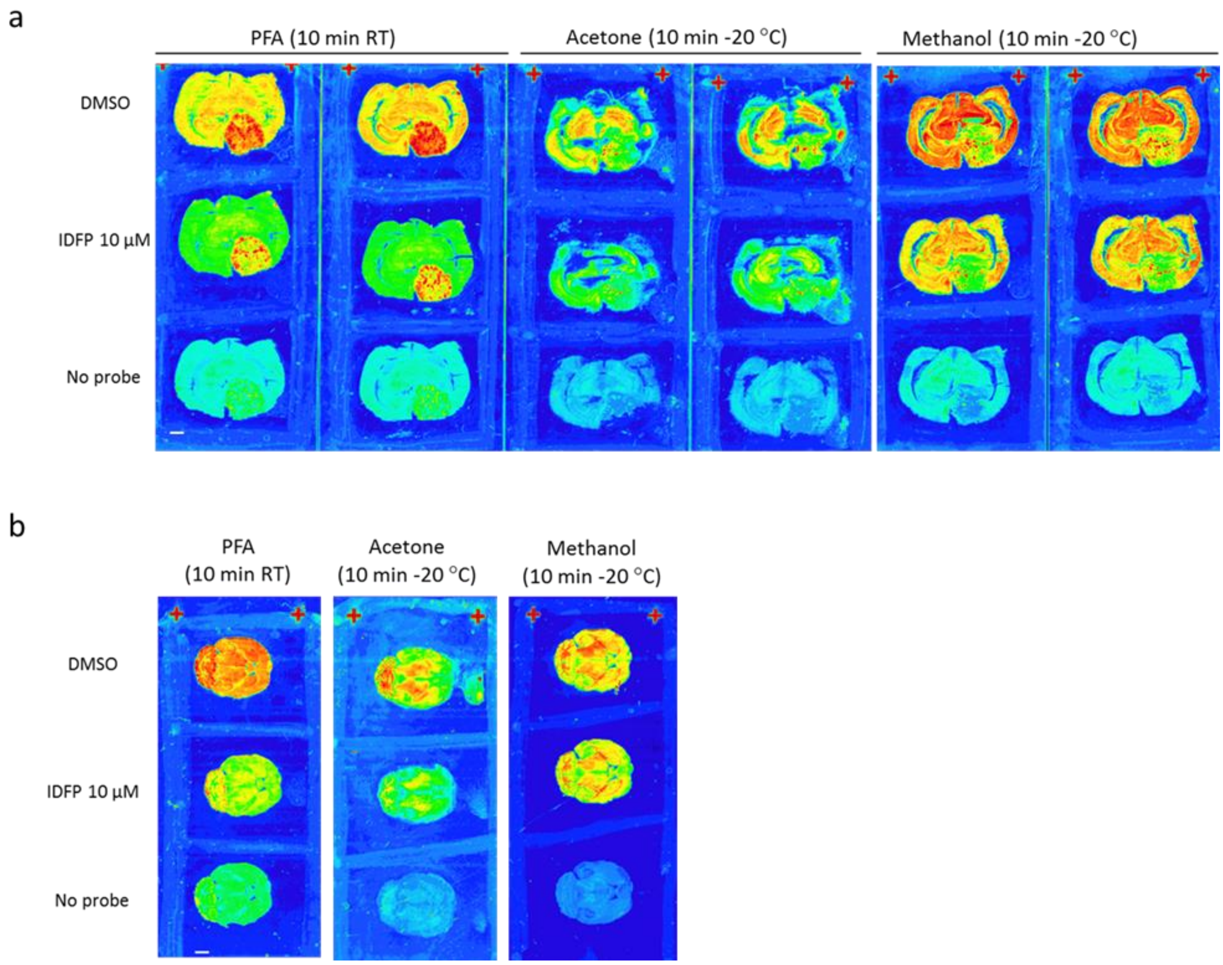


**Figure S4. Testing various fixation method for rodent brain sections.** Coronal sections (20 µm thick) of glioma brain (**a**) or horizontal sections of mouse brain (**b**) were fixed using either 4% paraformaldehyde supplemented with 0.01 % glutaraldehyde (PFA, 10 min at RT), or acetone (10 min at -20 °C) or methanol (10 min at -20 °C), washed for 2x5 min in 0.1 M PBS and processed thereafter for tissue-ABPP as detailed in Materials and Methods. Sections were pretreated with DMSO (top and bottom rows) or with the broad-spectrum SH inhibitor isopropyl dodecylfluorophosphonate (IDFP, 10 µM) for 1h at RT, after which they were incubated for 1h at RT with (top and middle rows) or without (bottom row) TAMRA-FP (0.5 µM). After washes, sections were imaged using Fuji gel scanner ($\lambda_{ex}$ 552 nm/$\lambda_{em}$ 575 nm) for visual assessment of tissue integrity and inhibitor sensitivity of TAMRA-FP labeling. Fluorescence intensity is shown in arbitrary color-scale where red and blue denotes high and low fluorescence, respectively. The images in **a** and **b** come from separate experiments and fluorescence colors are not comparable between the experiments. Note that while PFA fixation preserves both tissue integrity and inhibitor sensitivity of TAMRA-FP labeling, both parameters are compromised in sections fixed with acetone or methanol. Images in **a** are from duplicate slides with three consecutive coronal sections in each, cut from glioma rat 2 or from slides with three consecutive horizontal sections of normal mouse brain in each (**b**). Scale bar (white) 2 mm.

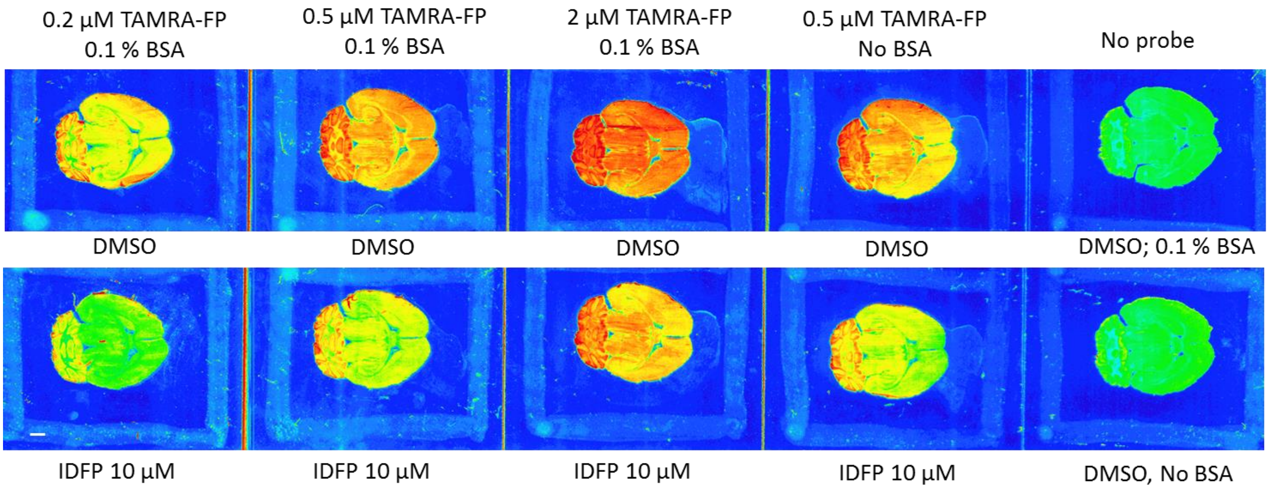


**Figure S5. Effect of TAMRA-FP concentration on fluorescence signal in mouse brain sections.** Horizontal mouse brain sections (20 µm thick) were fixed using PFA (10 min at RT), washed for 2x5 min in 0.1 M PBS and processed for tissue-ABPP as detailed in Materials and Methods. Sections were pretreated with DMSO or with 10 µM IDFP for 1h at RT, after which they were incubated for 1h at RT with or without increasing concentrations of TAMRA-FP (0.2, 0.5 or 2 µM). After washes, sections were imaged using Fuji gel scanner ($\lambda_{ex}$ 552 nm/$\lambda_{em}$ 575 nm) for visual assessment of fluorescence intensity and inhibitor sensitivity of probe labeling. Intensity of the fluorescence signal is shown as arbitrary colors where red and blue denotes high and low fluorescence, respectively. Note increasing signal intensity with increasing probe concentration. Note also IDFP-sensitivity of TAMRA-FP labeling regardless of probe concentration. Note also that omission of BSA (0.1 % w/v) only marginally affects signal intensity or inhibitor sensitivity. Scale bar (white) 2 mm.

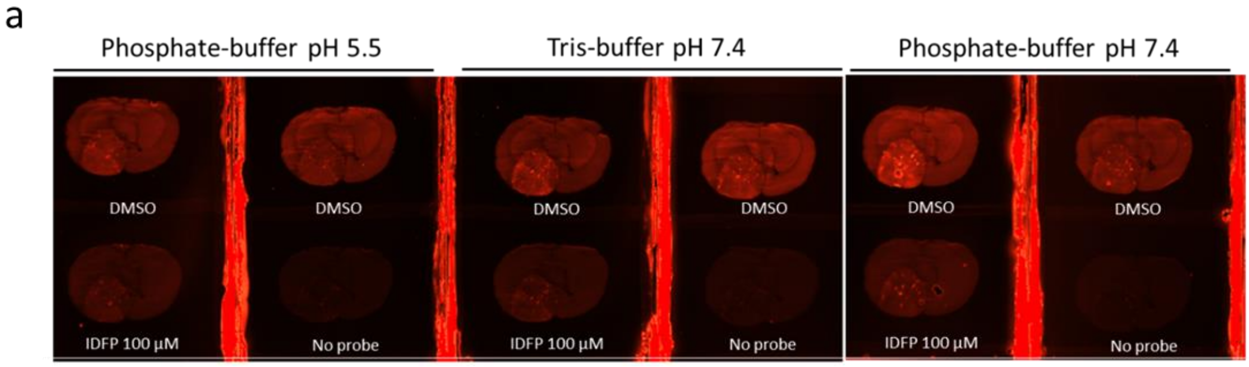


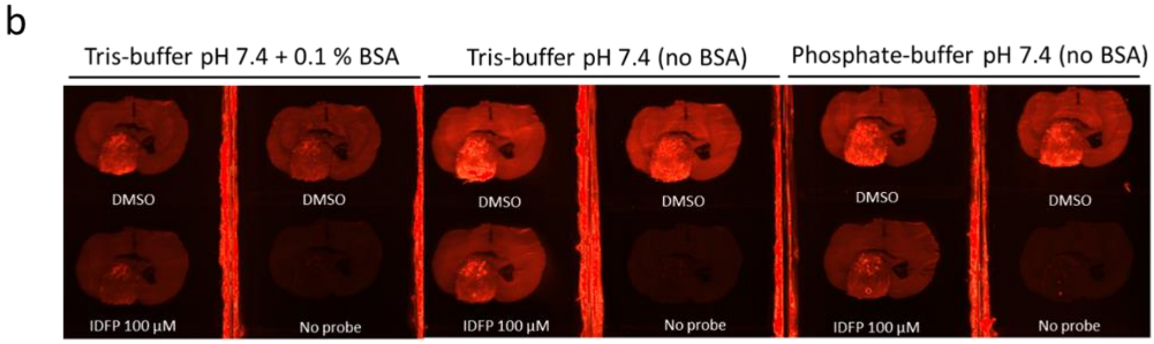


**Figure S6. Effects of pH and buffer composition on TAMRA-FP signal and its inhibitor sensitivity.** Coronal sections (20 µm thick) of rat glioma brain were fixed using PFA (10 min at RT), washed for 2x5 min in 0.1 M PBS and processed for tissue-ABPP, as detailed in Materials and Methods. Sections were pretreated with DMSO or with 100 µM IDFP for 1h at RT, followed by labeling step for 1h at RT with or without TAMRA-FP (0.5 µM) using the indicated buffer systems with varying pH (**a**) or buffer composition at pH 7.4 (**b**). After washes, sections were imaged using BioRad gel scanner (Cy3-window) for visual assessment of fluorescence intensity and inhibitor sensitivity of probe labeling. TAMRA-FP fluorescence is in red. **a.** Phosphate- vs. Tris-buffer (Tris-$MgCl_2$-NaCl-EDTA, pH 7.4), both containing 0.1 % BSA (w/v). Note that TAMRA-FP fluorescence is more intense at pH 7.4, being somewhat weaker in Tris-buffer at the same pH, and clearly blunted in the phosphate-buffer at pH 5.5. **b.** TAMRA-FP signal is comparable in Tris- vs. phosphate-buffer, inclusion of 0.1 % BSA clearly blunts TAMRA-FP fluorescence. Note that inhibitor sensitivity of TAMRA-FP labeling is retained in all conditions. For applications where imaging of maximal TAMRA-FP labeling is the desired endpoint readout, optimal signal is obtained in Tris- or phosphate buffer in the absence of BSA. For applications where TAMRA-FP labeling is followed by immunohistochemistry applied on the same section, inclusion of BSA is justified as a preceding facilitatory step to block tissue before application of primary antibody. Images were adjusted for brightness and contrast.

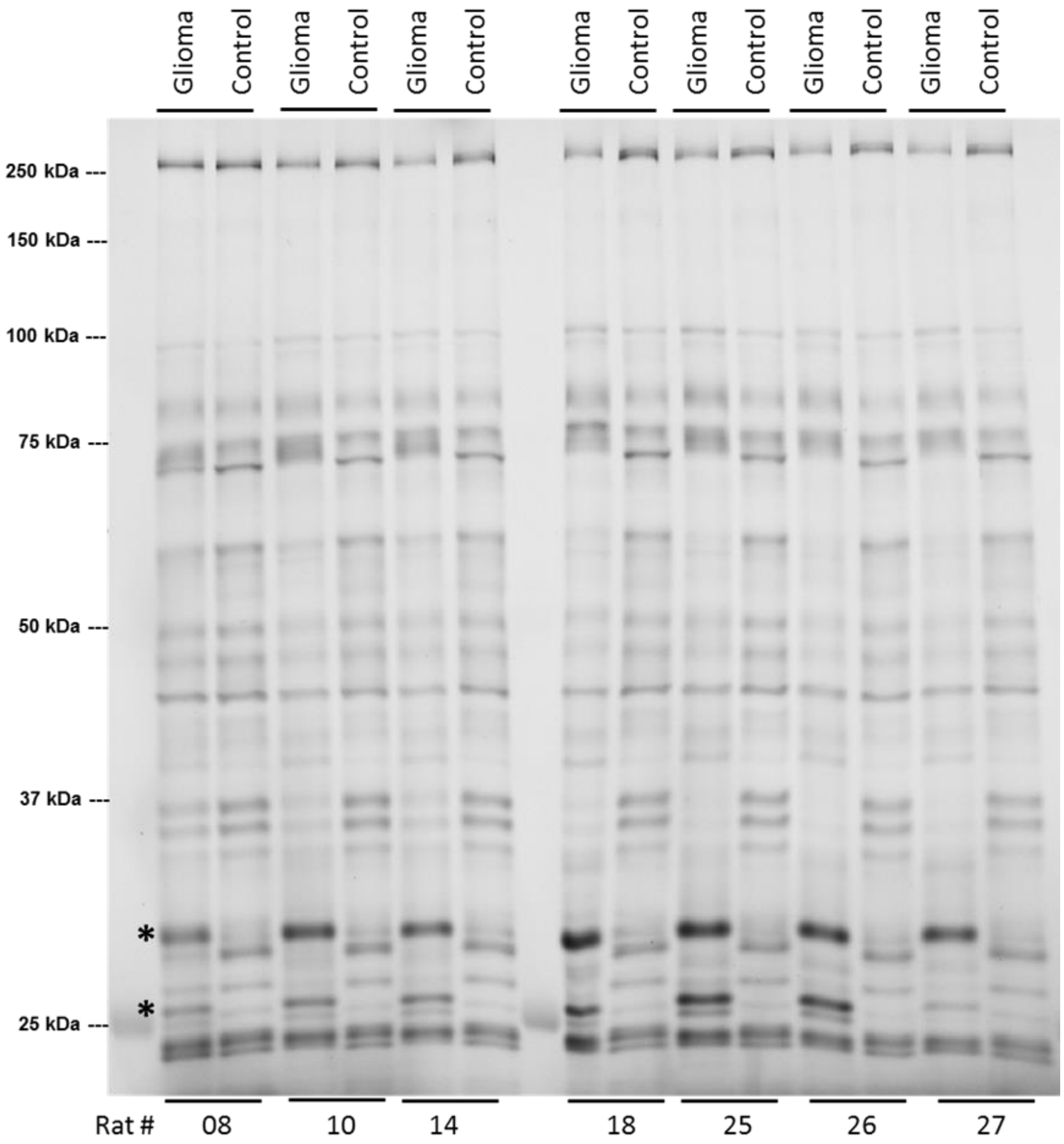


**Figure S7. Comparative gel-based ABPP of seven animals confirming distinct SH activity profiles between glioma and control brain.** Proteomes (1 mg/ml) were treated with TAMRA-FP (1 µM) for 1h at RT as detailed in Materials and Methods. The reaction was quenched and 12.5 µg protein was loaded per lane, followed by protein separation using SDS-PAGE. Shown are TAMRA-FP labeled bands (dark) after in-gel imaging. Position of molecular weight markers is indicated at left. Note similar activity profile between the seven gliomas; glioma-enriched SH bands (black asterisks) migrate at ~25 and ~30 kDa. Note also similar activity profiles between the seven control brains and particularly that the SH activity profiles of glioma and control brain are distinct. Image was adjusted for brightness and contrast.

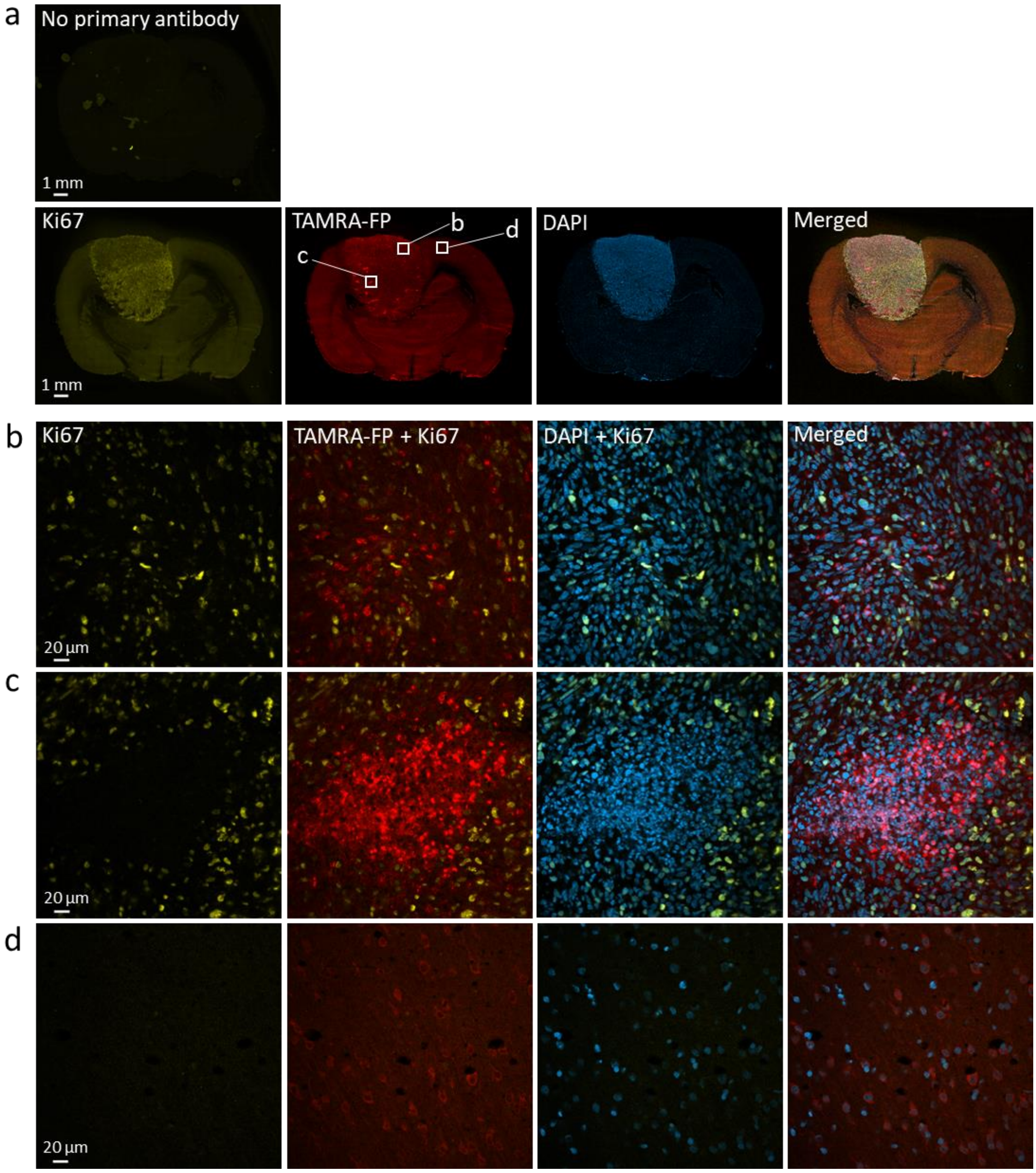


**Figure S8. Confocal imaging of SH activity in relation to proliferating tumor cells.** Sections went through the tissue-ABPP protocol to label SHs (red) and were thereafter immunostained for Ki67 to visualize proliferating cells (yellow), followed by DAPI staining to visualize nuclei (blue). Panel **a** shows overall staining pattern throughout the coronal section plane. A control section undergoing identical staining protocol with no primary antibody is illustrated at top. Panel **b** shows staining pattern in glioma region characterized by intense SH activity originating from individual cells (TAMRA-FP hotspots). Panel **c** shows staining pattern in glioma region characterized by intense SH activity originating from cell clusters (TAMRA-FP hotspot clusters). Panel **d** shows staining pattern in healthy brain (cortex). Note scattered population of proliferating cells with large nuclei and with distinct morphology in region of TAMRA-FP hotspots (**b**). Note proliferating cells around the region with TAMRA-FP hotspot clusters and lack of co-localization with TAMPA-FP signal (**c**). Note absence of Ki67 immunostaining from the healthy brain (**d**). Magnification 10x in **a**, 40x in **b-d**. Primary antibody rabbit anti-Ki67 (Abcam, cat# ab15580), dilution 1:500, secondary antibody goat anti-rabbit IgG-Alexa Fluor 647 conjugate, dilution 1:100. Sections were from male rat 29. Scale bars: 1 mm in a, 20 µm in b-d. Images were adjusted for brightness and contrast.

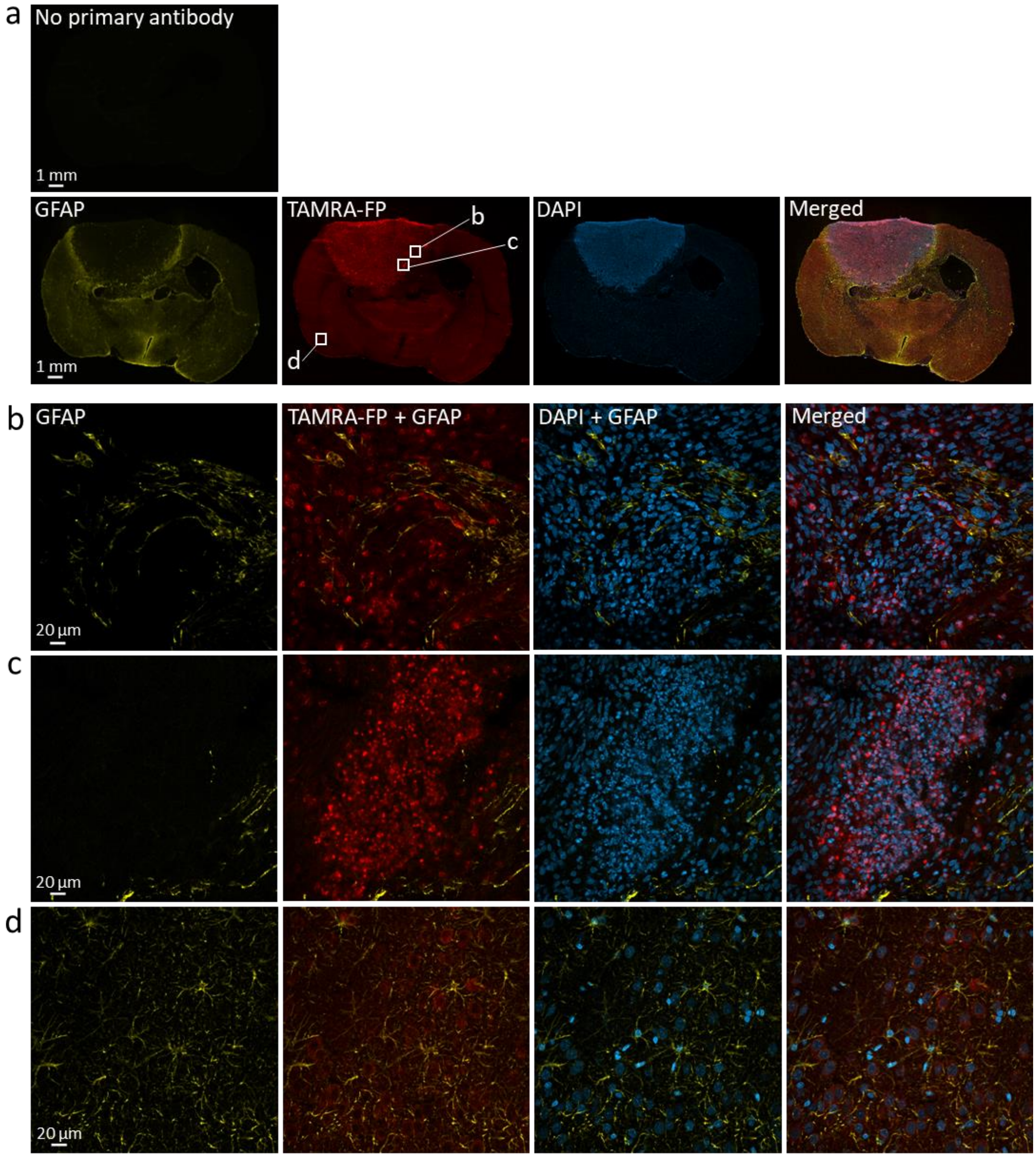


**Figure S9. Confocal imaging of SH activity in relation to astrocytes.** Sections went through the tissue-ABPP protocol to label SHs (red) and were thereafter immunostained for GFAP to visualize astrocytes (yellow), followed by DAPI staining to visualize nuclei (blue). Panel **a** shows overall staining pattern throughout the coronal section plane. A control section undergoing identical staining protocol with no primary antibody is illustrated at top. Panel **b** shows staining pattern in glioma region characterized by intense SH activity originating from individual cells (TAMRA-FP hotspots). Panel **c** shows staining pattern in glioma region characterized by intense SH activity originating from cell clusters (TAMRA-FP hotspot clusters). Panel **d** shows staining pattern in control region (basal forebrain). Note dense astrocyte population encircling the tumor (**a**). Note sparse overall presence of astrocytes in the tumor (**a**) and their presence in the tumor margin (**b**). Note poor co-localization of GFAP-positive cells with TAMRA-FP hotspots (**b**). Note also lack of astrocytes from TAMRA-FP hotspot clusters (**c**). In healthy brain, star-shaped cells (i.e. astrocytes) are clearly visible (**d**). Magnification 10x in **a**, 40x in **b-d**. Primary antibody goat anti-GFAP (Abcam, cat# ab53554), dilution 1:500, secondary antibody donkey anti-goat IgG-Alexa Fluor 647 conjugate, dilution 1:500. Sections were from male rat 33. Scale bars: 1 mm in a, 20 µm in b-d. Images were adjusted for brightness and contrast.

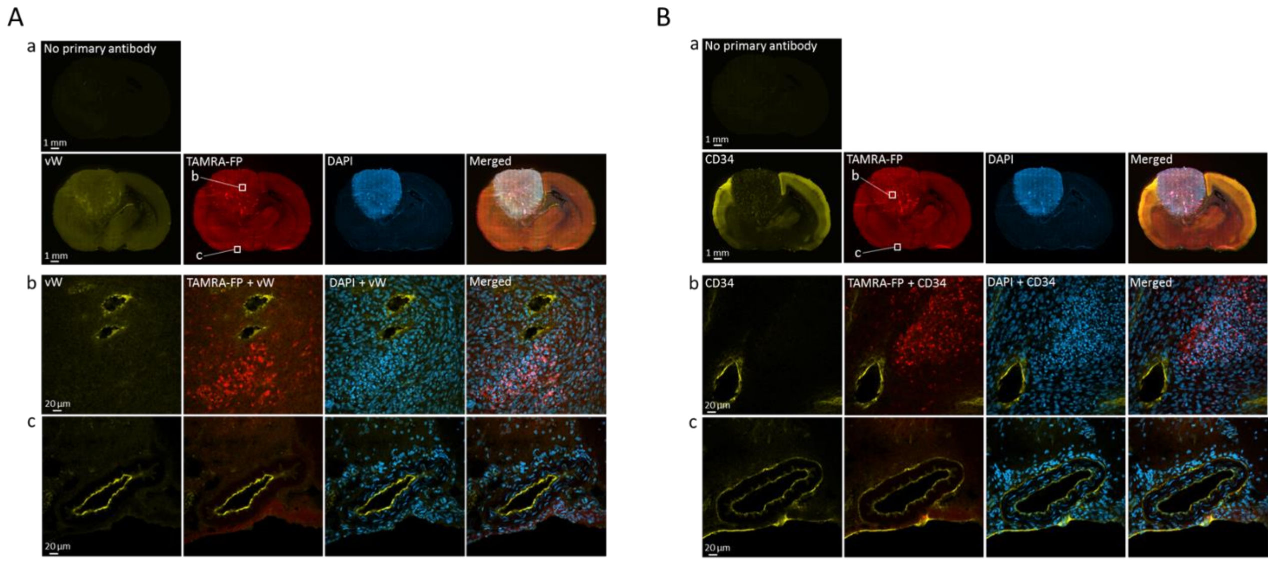


**Figure S10. Confocal imaging of SH activity in relation to blood vessels.** Vasculature was visualized using the endothelial markers von Willebrand factor (vW) and CD34. Sections went through the tissue-ABPP protocol to label SHs (red) and were thereafter immunostained for vW (**A**) or CD34 (**B**) to visualize endothelial cells (yellow), followed by DAPI staining to visualize nuclei (blue). Panel **a** shows overall staining pattern throughout the coronal section plane. A control section undergoing identical staining protocol with no primary antibody is illustrated at top. Panel **b** shows staining pattern in glioma region with intense SH activity (TAMRA-FP hotspot clusters). Panel **c** shows staining pattern in control brain (hypotalamus). Note intense labeling of endothelial cells lining the vessels by both markers (**b**). Note TAMRA-FP hotspot clusters in close vicinity of the vessels (**b**). In healthy brain (**c**), endothelial cells lining the vessels are labeled by both markers. Magnification 10x in **a**, 40x in **b-c**. In **A**, primary antibody rabbit anti-von Willebrand Factor (Abcam, cat# ab6994), dilution 1:2000, secondary antibody goat anti-rabbit IgG-Alexa Fluor 647 conjugate, dilution 1:500. Sections were from female rat 11. In **B**, primary antibody goat anti-CD34 (R&D systems, cat# AF4117), dilution 1:500, secondary antibody donkey anti-goat IgG-Alexa Fluor 647 conjugate, dilution 1:100. Sections were from female rat 11. Scale bars: 1 mm in a, 20 µm in b-c. Images were adjusted for brightness and contrast.

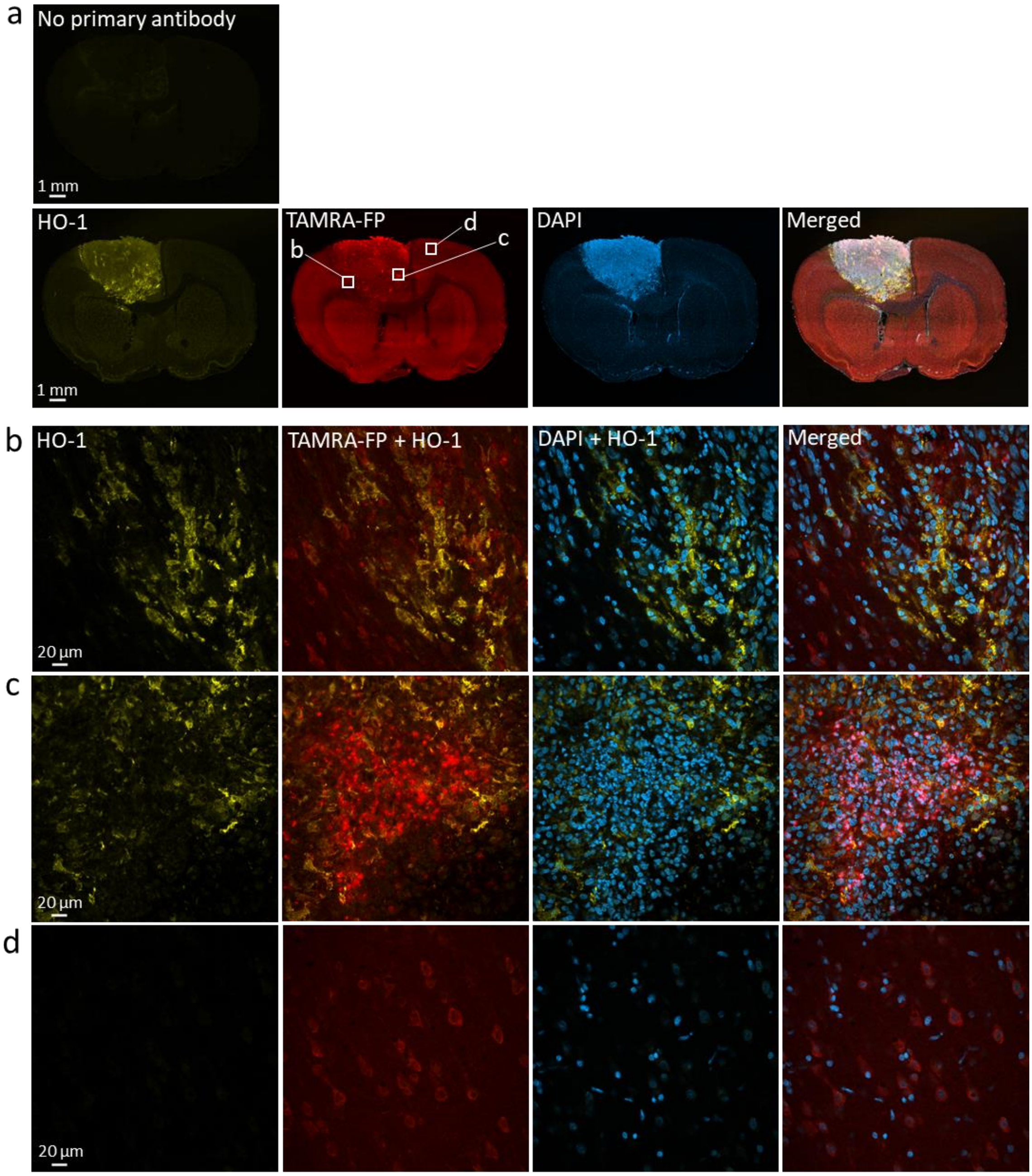


**Figure S11. Confocal imaging of SH activity in relation to heme oxygenase 1 (HO-1).** Sections went through the tissue-ABPP protocol to label SHs (red) and were thereafter immunostained for HO-1 to visualize redox-sensitive areas (yellow), followed by DAPI staining to visualize nuclei (blue). Panel **a** shows overall staining pattern throughout the coronal section plane. A control section undergoing identical staining protocol with no primary antibody is illustrated at top. Panel **b** shows staining pattern in glioma region characterized by intense SH activity originating from individual cells (TAMRA-FP hotspots). Panel **c** shows staining pattern in glioma region characterized by intense SH activity originating from cell clusters (TAMRA-FP hotspot clusters). Panel **d** shows staining pattern in healthy brain (cortex). Note heterogeneous pattern of HO-1 expression over the tumor and its absence from healthy brain. Note poor co-localization of TAMRA-FP labeling with the stress marker. Primary antibody mouse anti-heme oxygenase 1 (GTS-1, Abcam, cat# ab12220), dilution 1:500, secondary antibody donkey anti-mouse IgG-Alexa Fluor 647 conjugate, dilution 1:100. Sections were from male rat 29. Scale bars: 1 mm in a, 20 µm in b-d. Images were adjusted for brightness and contrast.

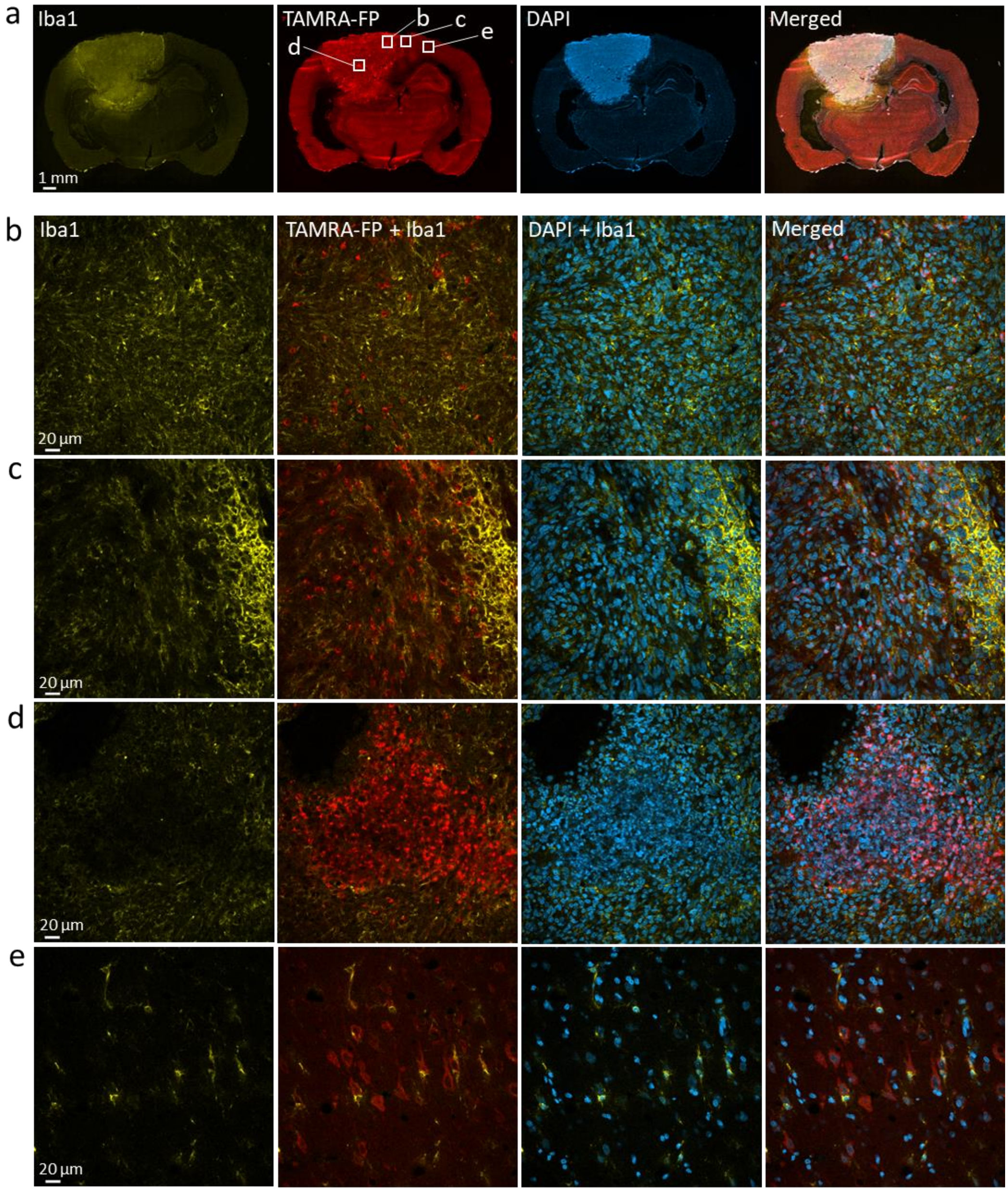


**Figure S12. Confocal imaging of SH activity in relation to microglial marker Iba1.** Sections went through the tissue-ABPP protocol to label SHs (red) and were thereafter immunostained for Iba1 to visualize microglia (yellow), followed by DAPI staining to visualize nuclei (blue). Panel **a** shows overall staining pattern throughout the coronal section plane. Due to technical problem, a control section without primary antibody is not available for this experiment. Panel **b** shows staining pattern in glioma region characterized by intense SH activity originating from individual cells (TAMRA-FP hotspots). Panel **c** shows staining pattern in glioma border with more Iba1-positive cells residing on the healthy side. Panel **d** shows staining pattern in glioma region characterized by intense SH activity originating from cell clusters (TAMRA-FP hotspot clusters). Panel **e** shows staining pattern in control brain (cortex). Note heterogeneous presence of Iba1-positive cells throughout the tumor (**a**) and especially its border (**c**). Note partial overlap of Iba1 staining with TAMRA-FP hotspots (**b**). In contrast, no Iba1-positive cells are visible in region of TAMRA-FP hotspot clusters (**d**). In control brain, Iba1-positive cells with characteristic microglia morphology and low TAMRA-FP fluorescence are visible. Primary antibody goat anti-Iba1 (Abcam, cat# ab5076), dilution 1:1000, secondary antibody donkey anti-goat IgG-Alexa Fluor 647 conjugate, dilution 1:100. Sections were from female rat 11. Scale bars: 1 mm in a, 20 µm in b-d. Images were adjusted for brightness and contrast.

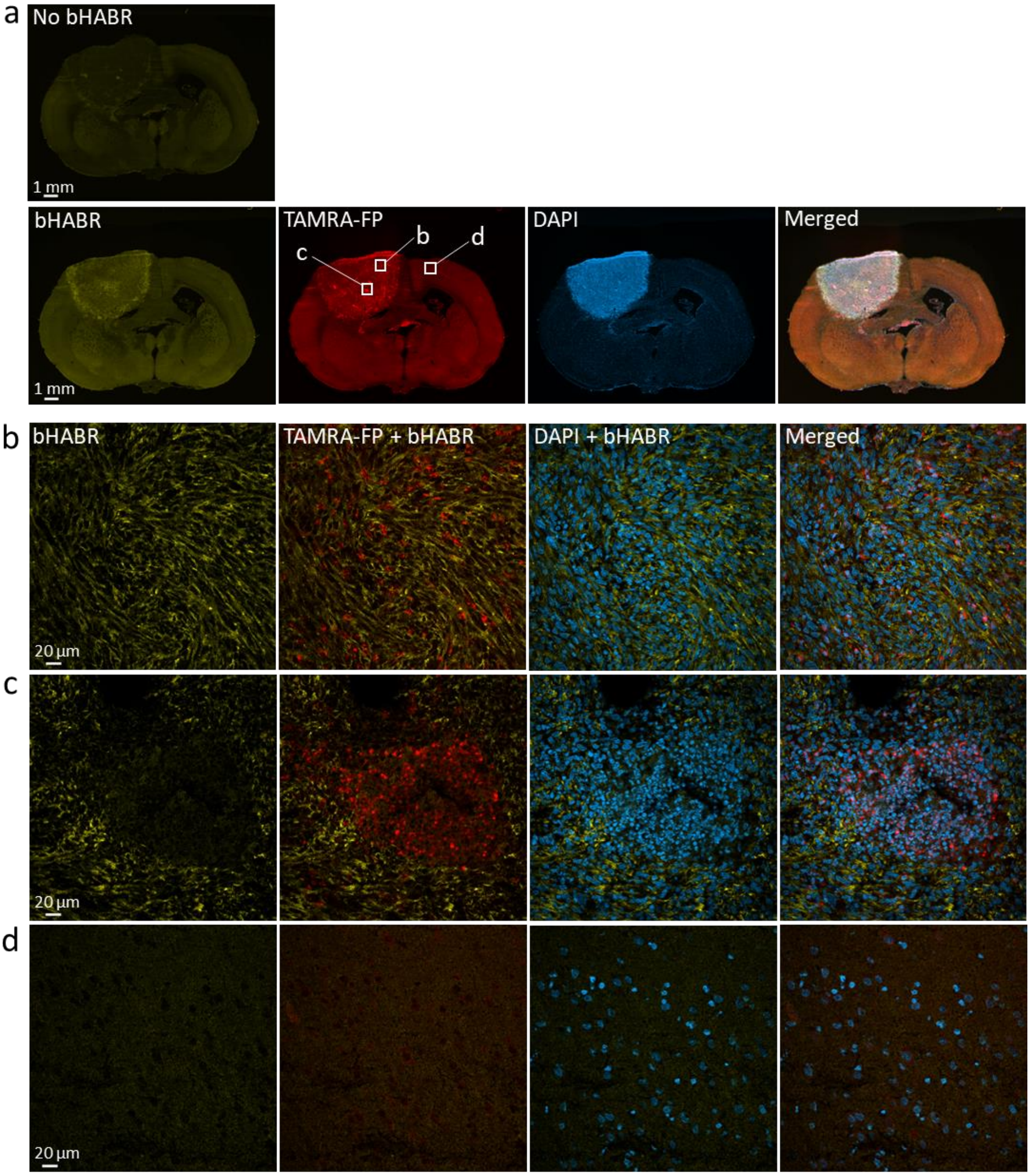


**Figure S13. Confocal imaging of SH activity in relation to hyaluronan (HA).** Sections went through the tissue-ABPP protocol to label SHs (red) and were thereafter stained for HA (yellow), followed by DAPI staining to visualize nuclei (blue). Panel **a** shows overall staining pattern throughout the coronal section plane. A control section undergoing identical staining protocol with no bHABR is illustrated at top. Panel **b** shows staining pattern in glioma region characterized by intense SH activity originating from individual cells (TAMRA-FP hotspots). Panel **c** shows staining pattern in glioma region characterized by intense SH activity originating from cell clusters (TAMRA-FP hotspot clusters). Panel **d** shows staining pattern in healthy brain (cortex). Note abundance of HA in the extracellular matrix throughout the brain with more intense staining over glioma, notably glioma edges. Note presence of HA in region of TAMRA-FP hotspots (**b**). Note also absence of HA from TAMRA-FP hotspot clusters, as well as encircling of these clusters by HA-enriched matrix (**c**). Relatively low abundance of HA is evident in healthy brain (**d**). HA-binding probe (bHABR, prepared in-house), dilution 1:30, secondary antibody DyLight488 streptavidin, dilution 1:900. Sections were from male rat 34. Scale bars: 1 mm in a, 20 µm in b-d. Images were adjusted for brightness and contrast.

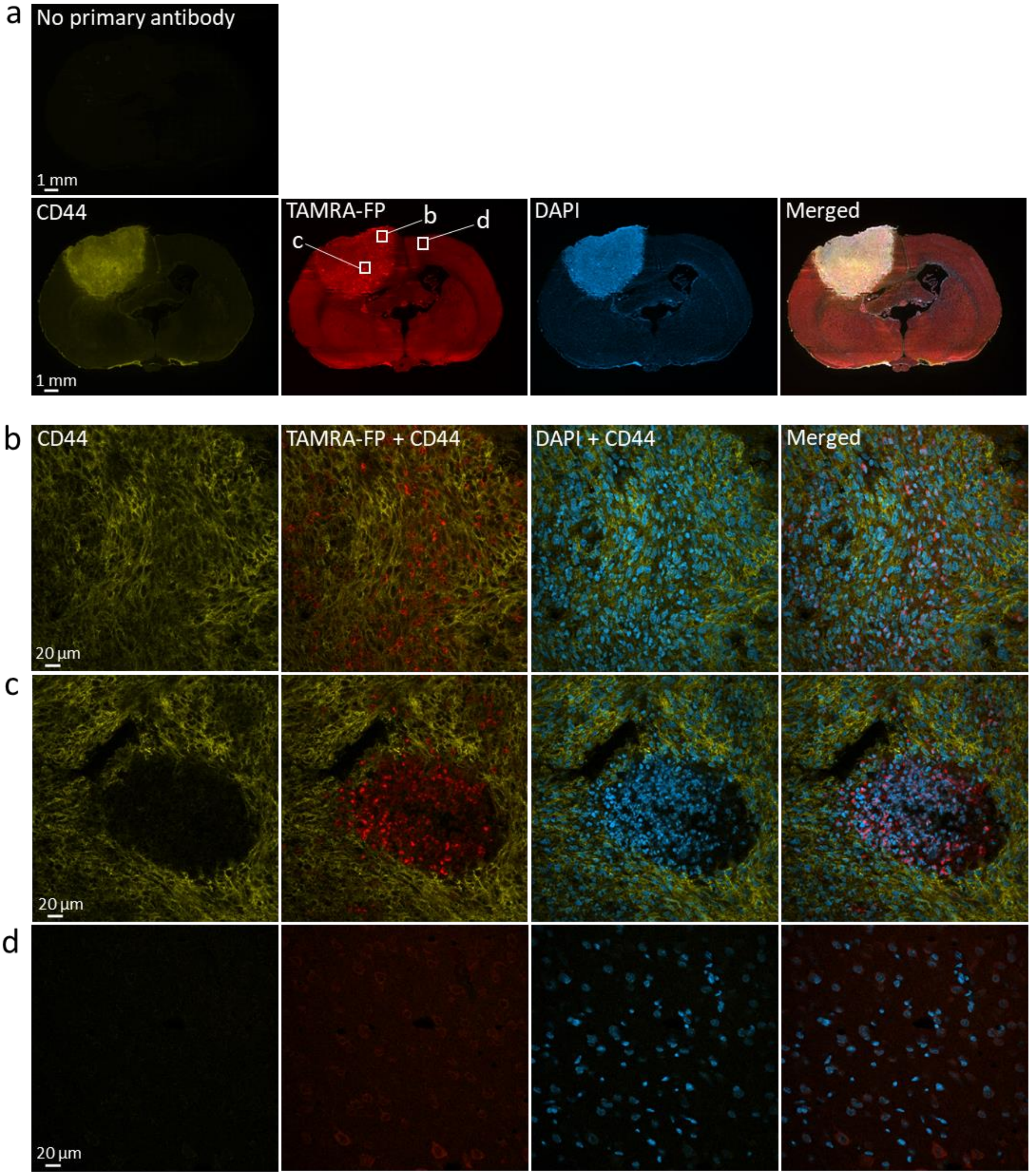


**Figure S14. Confocal imaging of SH activity in relation to HA receptor CD44.** Sections went through the tissue-ABPP protocol to label SHs (red) and were thereafter immunostained for CD44 (yellow), followed by DAPI staining to visualize nuclei (blue). Panel **a** shows overall staining pattern throughout the coronal section plane. A control section undergoing identical staining protocol with no primary antibody is illustrated at top. Panel **b** shows staining pattern in glioma region characterized by intense SH activity originating from individual cells (TAMRA-FP hotspots). Panel **c** shows staining pattern in glioma region characterized by intense SH activity originating from cell clusters (TAMRA-FP hotspot clusters). Panel **d** shows staining pattern in healthy brain (cortex). Note wide, heterogeneous distribution of CD44 in glioma. Note CD44-positive cells in region of TAMRA-FP hotspots (**b**). Note absence of CD44 from TAMRA-FP hotspot clusters (**c**). Note also that these clusters are surrounded by CD44-positive cells (**c**). CD44 is undetectable in healthy brain (**d**). Primary antibody rabbit anti-CD44 (Abcam, cat# ab157107), dilution 1:2000, secondary antibody goat anti-rabbit IgG-Alexa Fluor 647 conjugate, dilution 1:100. Sections were from male rat 34. Scale bars: 1 mm in a, 20 µm in b-d. Images were adjusted for brightness and contrast.

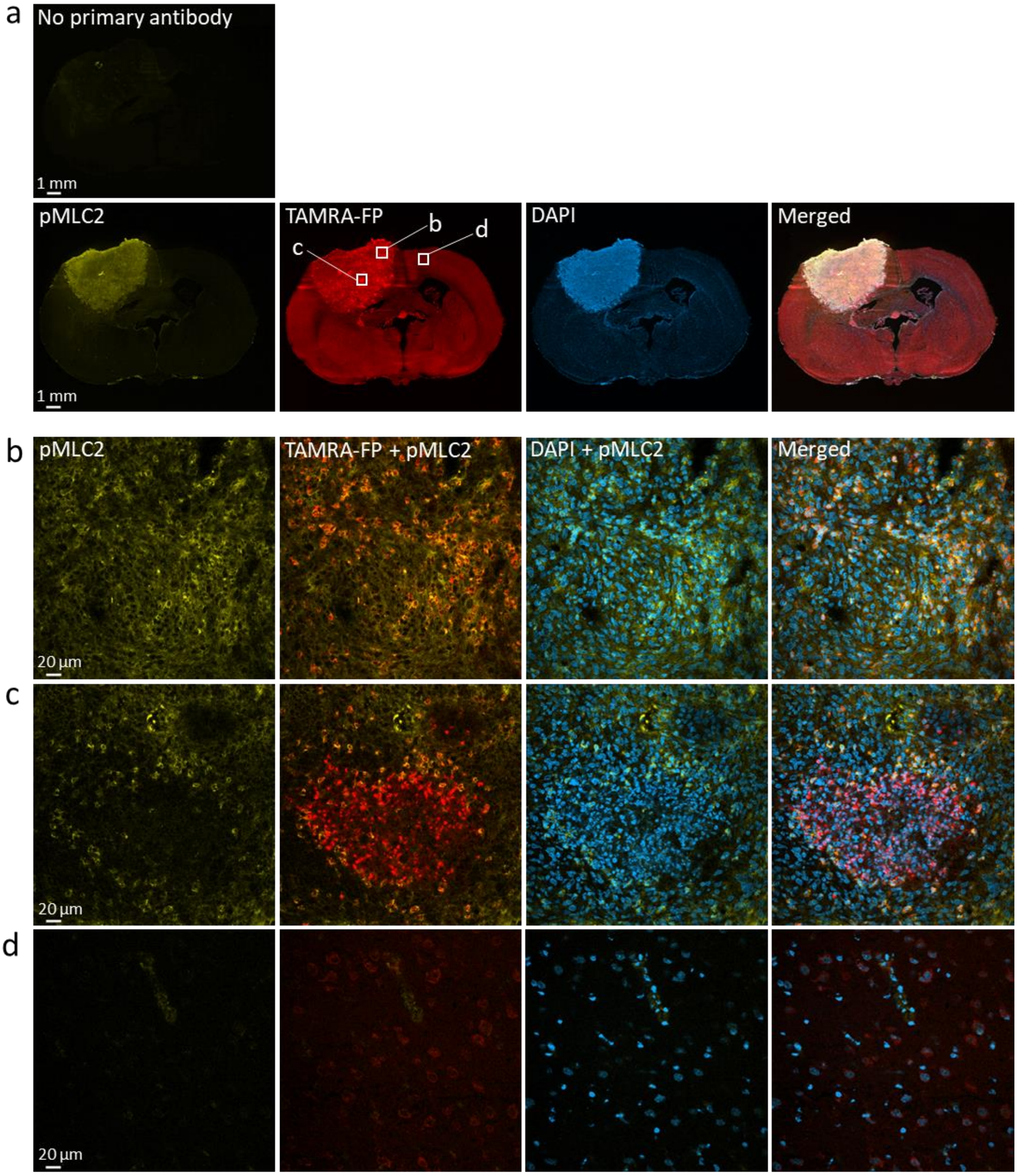


**Figure S15. Confocal imaging of SH activity in relation to the stiffness marker pMLC2.** Sections went through the tissue-ABPP protocol to label SHs (red) and were thereafter immunostained for pMLC2 (yellow), followed by DAPI staining to visualize nuclei (blue). Panel **a** shows overall staining pattern throughout the coronal section plane. A control section undergoing identical staining protocol with no primary antibody is illustrated at top. Panel **b** shows staining pattern in glioma region characterized by intense SH activity originating from individual cells (TAMRA-FP hotspots). Panel **c** shows staining pattern in glioma region characterized by intense SH activity originating from cell clusters (TAMRA-FP hotspot clusters). Panel **d** shows staining pattern in healthy brain (cortex). Note abundance of pMLC2 throughout the glioma (**a**) and that the stiffness marker co-localizes with TAMRA-FP hotspots (**b**). Note low pMLC2 expression within the TAMRA-FP hotspot clusters (**c**). Note also that the TAMRA-FP positive cluster rim and cells surrounded by the cluster contain pMLC2-positive cells (**c**). pMLC2 is undetectable in cortical control region (**d**). Primary antibody mouse anti-Phospho-Myosin Light Chain 2 (pMLC2, Cell Signaling Technology, cat# 3675s), dilution 1:50, secondary antibody donkey anti-mouse IgG-Alexa Fluor 647 conjugate, dilution 1:100. Sections were from male rat 34. Scale bars: 1 mm in a, 20 µm in b-d. Images were adjusted for brightness and contrast.

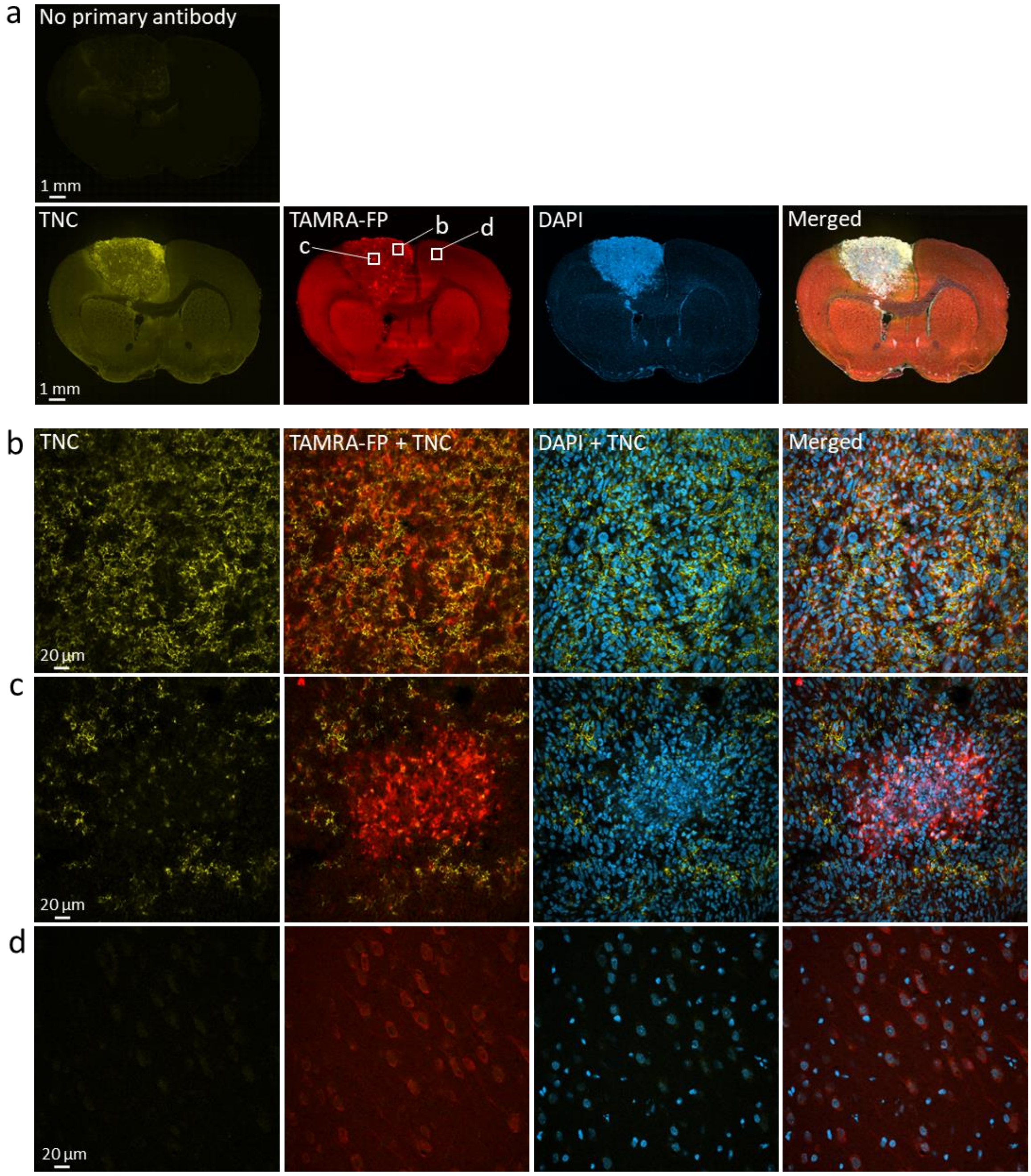


**Figure S16. Confocal imaging of SH activity in relation to the stiffness marker tenascin C.** Sections went through the tissue-ABPP protocol to label SHs (red) and were thereafter immunostained for tenascin C (TNC, yellow), followed by DAPI staining to visualize nuclei (blue). Panel **a** shows overall staining pattern throughout the coronal section plane. A control section undergoing identical staining protocol with no primary antibody is illustrated at top. Panel **b** shows staining pattern in glioma region characterized by intense SH activity originating from individual cells (TAMRA-FP hotspots). Panel **c** shows staining pattern in glioma region characterized by intense SH activity originating from cell clusters (TAMRA-FP hotspot clusters). Panel **d** shows staining pattern in healthy brain (cortex). Note heterogeneous expression of tenascin C in the glioma (**a**) and that tenascin C co-localizes with TAMRA-FP hotspots (**b**). Note low tenascin C expression in TAMRA-FP hotspot clusters and that the cells around these clusters frequently express tenascin C (**c**). Tenascin C is undetectable in cortical control region (**d**). Primary antibody mouse anti-Tenascin C (Thermo Scientific, cat# MA5-16086), dilution 1:100, secondary antibody donkey anti-mouse IgG-Alexa Fluor 647 conjugate, dilution 1:100. Sections were from male rat 29. Scale bars: 1 mm in a, 20 µm in b-d. Images were adjusted for brightness and contrast.

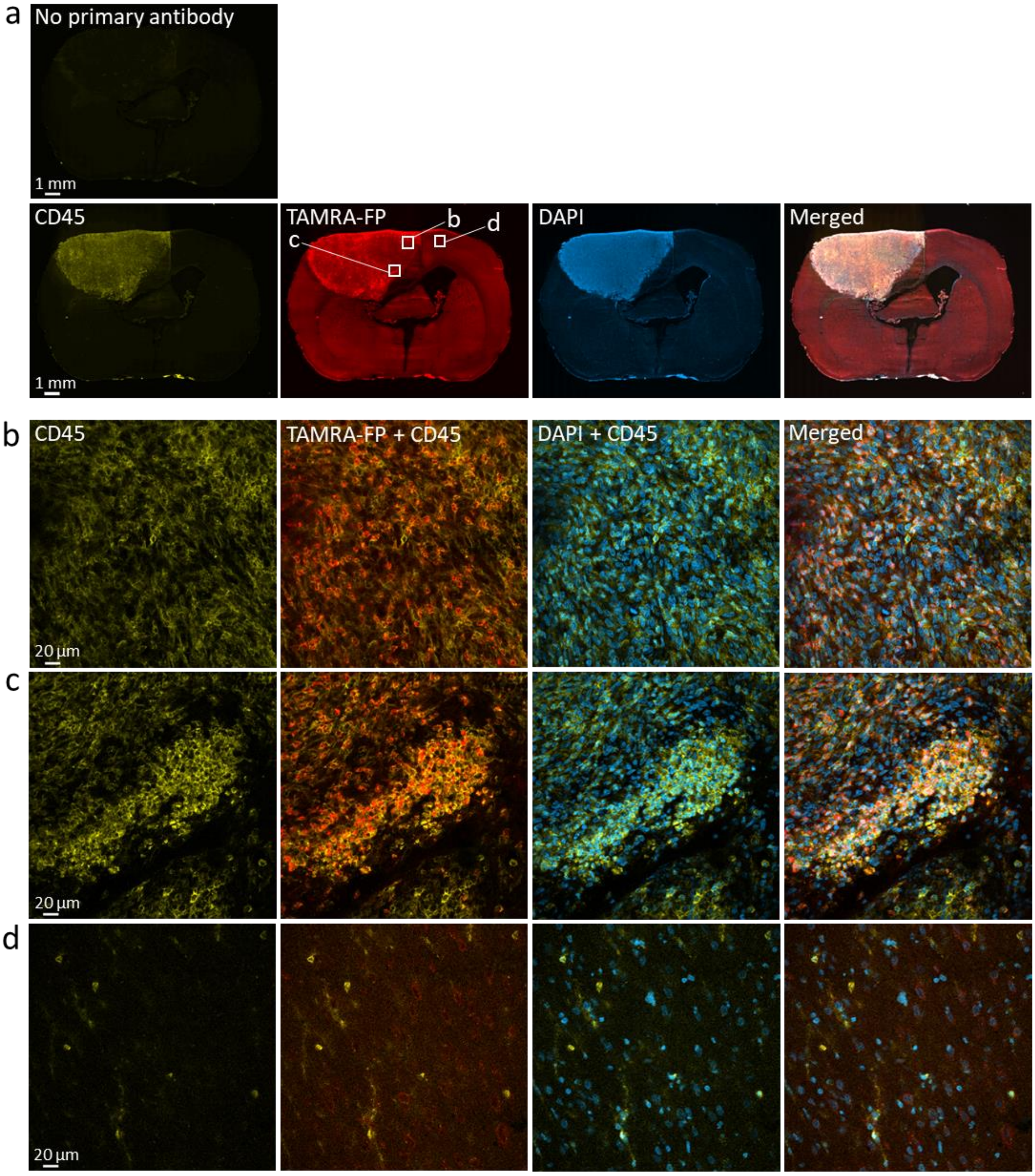


**Figure S17. Confocal imaging of SH activity in relation to CD45, a marker for nucleated hematopoietic cells.** Sections went through the tissue-ABPP protocol to label SHs (red) and were thereafter immunostained for CD45 (yellow), followed by DAPI staining to visualize nuclei (blue). Panel **a** shows overall staining pattern throughout the coronal section plane. A control section undergoing identical staining protocol with no primary antibody is illustrated at top. Panel **b** shows staining pattern in glioma region characterized by intense SH activity originating from individual cells (TAMRA-FP hotspots). Panel **c** shows staining pattern in glioma region characterized by intense SH activity originating from cell clusters (TAMRA-FP hotspot clusters). Panel **d** shows staining pattern in healthy brain (cortex). Note ample expression of CD45 in the glioma (**a**) and that this marker shows co-localization with TAMRA-FP hotspots (**b**). Note also abundance of CD45 and its co-localization with TAMRA-FP hotspot clusters (**c**). Sparse immunostaining is visible in control cortical region (**d**). Primary antibody mouse anti-CD45 (MRC-OX1, Abcam, ab33923), dilution 1:500, secondary antibody donkey anti-mouse IgG-Alexa Fluor 647 conjugate, dilution 1:100. Sections were from male rat 33. Scale bars: 1 mm in a, 20 µm in b-d. Images were adjusted for brightness and contrast.

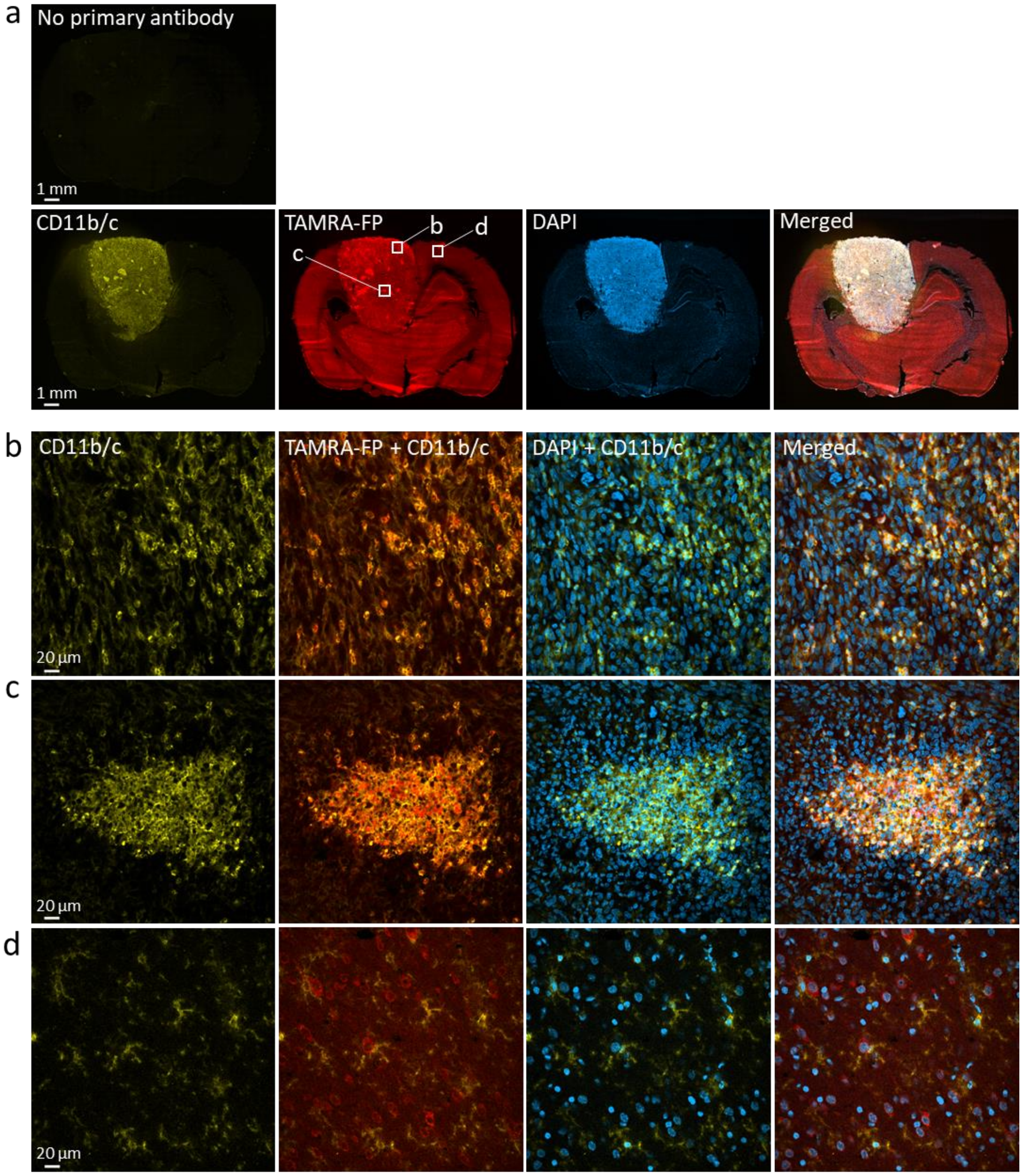


**Figure S18. Confocal imaging of SH activity in relation to CD11b/c, a marker for phagocytes.** Sections went through the tissue-ABPP protocol to label SHs (red) and were thereafter immunostained for CD11b/c (yellow), followed by DAPI staining to visualize nuclei (blue). Panel **a** shows overall staining pattern throughout the coronal section plane. A control section undergoing identical staining protocol with no primary antibody is illustrated at top. Panel **b** shows staining pattern in glioma region characterized by intense SH activity originating from individual cells (TAMRA-FP hotspots). Panel **c** shows staining pattern in glioma region characterized by intense SH activity originating from cell clusters (TAMRA-FP hotspot clusters). Panel **d** shows staining pattern in healthy brain (cortex). Note ample expression of CD11b/c in the glioma (**a**) and that this marker co-localizes with TAMRA-FP hotspots (**b**). Note also high CD11b/c expression and co-localization with TAMRA-FP hotspot clusters (**c**). CD11b/c-positive cells with characteristic microglial morphology are visible in cortical control region (**d**). Primary antibody mouse anti-CD11b/c (OX42, Abcam, ab1211), dilution 1:1000, secondary antibody donkey anti-mouse IgG-Alexa Fluor 647 conjugate, dilution 1:100. Sections were from male rat 29. Scale bars: 1 mm in a, 20 µm in b-d. Images were adjusted for brightness and contrast.

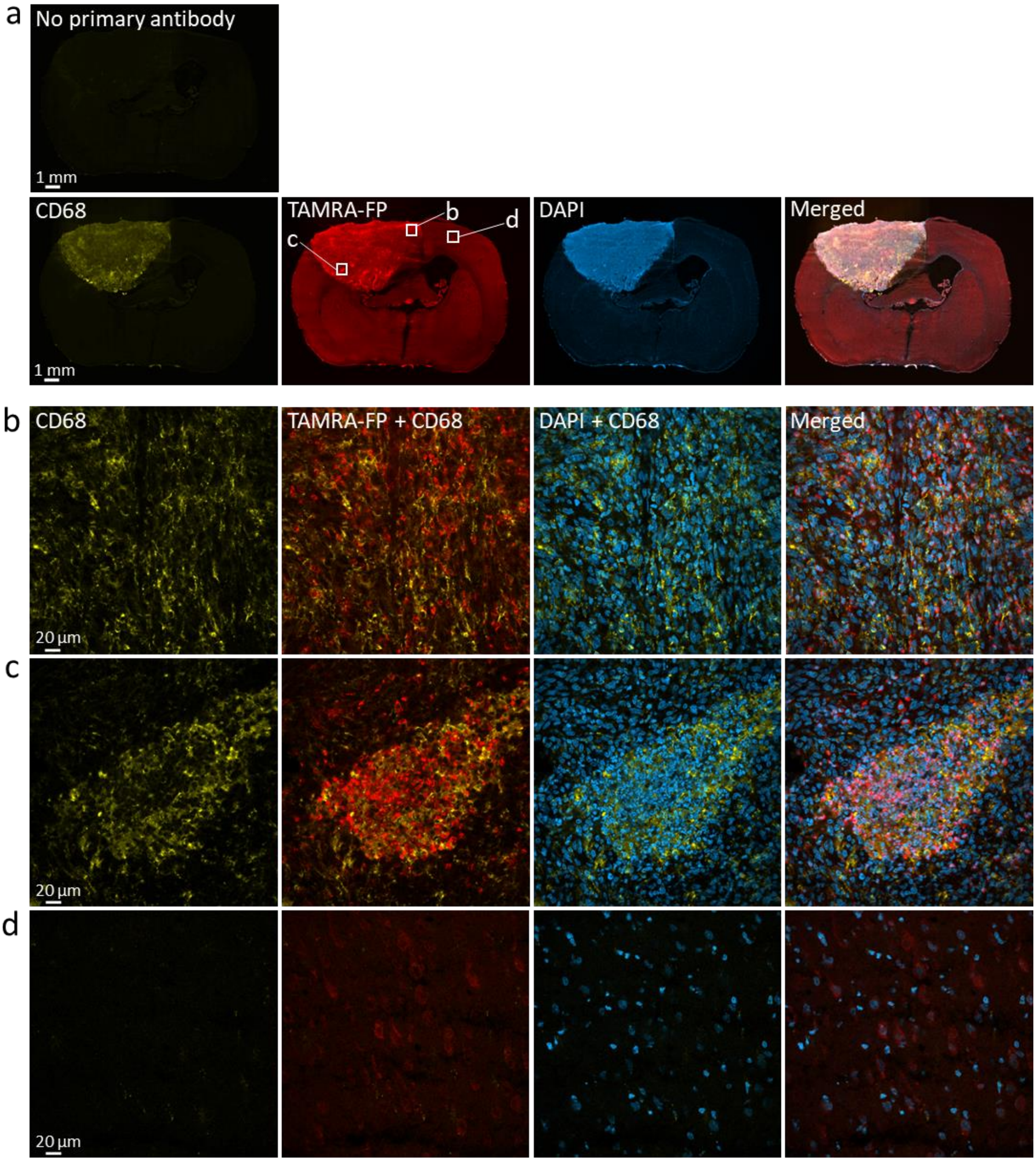


**Figure S19. Confocal imaging of SH activity in relation to CD68, a marker for monocytes and macrophages.** Sections went through the tissue-ABPP protocol to label SHs (red) and were thereafter immunostained for CD68 (yellow), followed by DAPI staining to visualize nuclei (blue). Panel **a** shows overall staining pattern throughout the coronal section plane. A control section undergoing identical staining protocol with no primary antibody is illustrated at top. Panel **b** shows staining pattern in glioma region characterized by intense SH activity originating from individual cells (TAMRA-FP hotspots). Panel **c** shows staining pattern in glioma region characterized by intense SH activity originating from cell clusters (TAMRA-FP hotspot clusters). Panel **d** shows staining pattern in healthy brain (cortex). Note intense expression of CD68 in the glioma (**a**) and its partial co-localization with TAMRA-FP hotspots (**b**). Note enrichment of CD68 and its partial co-localization with TAMRA-FP hotspot clusters (**c**). CD68 is undetectable in control cortical region (**d**). Primary antibody mouse anti-CD68 (ED1, Bio-Rad, cat# MCA341R), dilution 1:1000, secondary antibody donkey anti-mouse IgG-Alexa Fluor 647 conjugate, dilution 1:100. Sections were from male rat 33. Scale bars: 1 mm in a, 20 µm in b-d. Images were adjusted for brightness and contrast.

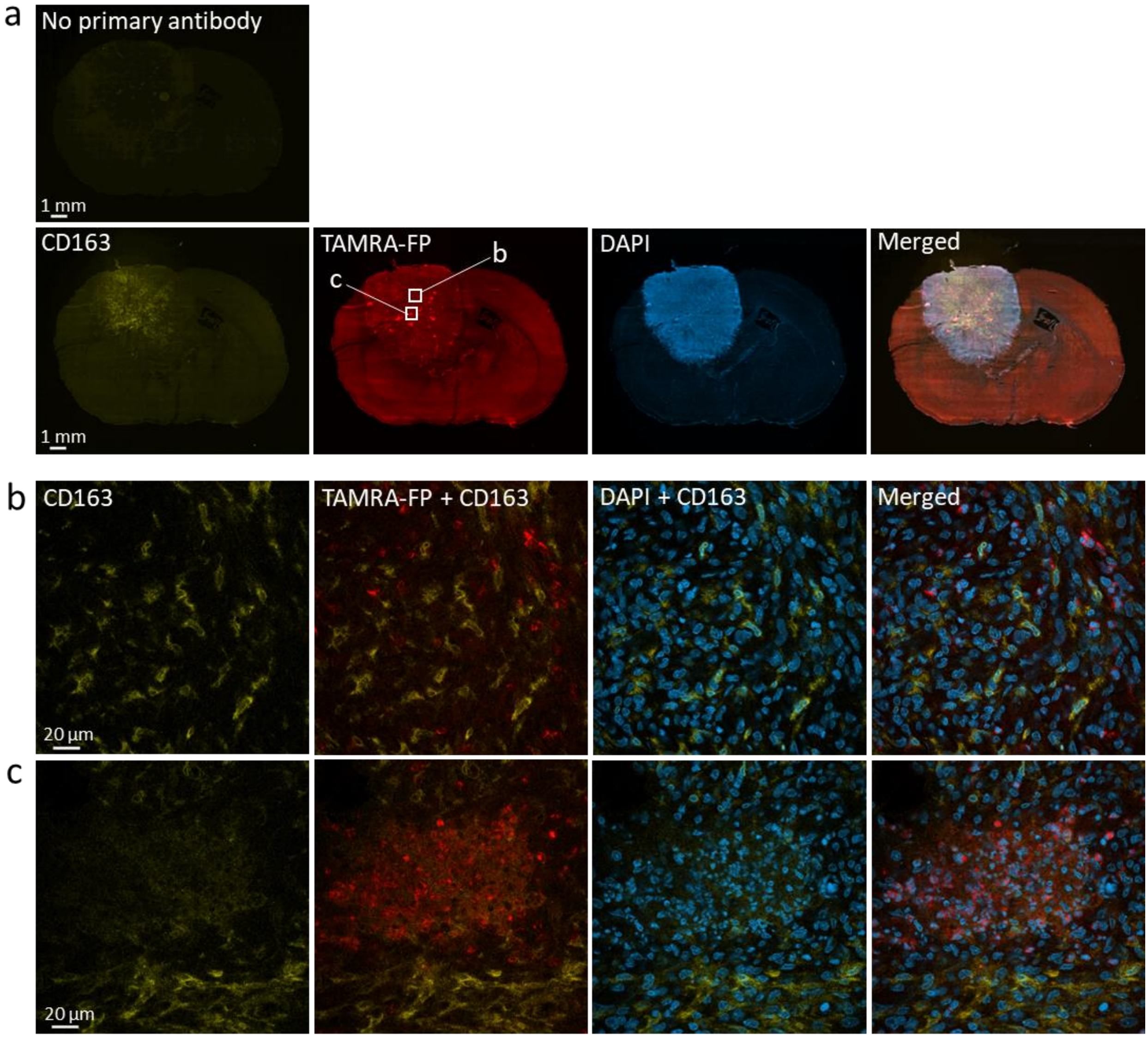


**Figure S20. Confocal imaging of SH activity in relation to CD163, a marker for monocytes and macrophages.** Sections went through the tissue-ABPP protocol to label SHs (red) and were thereafter immunostained for CD163 (yellow), followed by DAPI staining to visualize nuclei (blue). Panel **a** shows overall staining pattern throughout the coronal section plane. A control section undergoing identical staining protocol with no primary antibody is illustrated at top. Panel **b** shows staining pattern in glioma region characterized by intense SH activity originating from individual cells (TAMRA-FP hotspots). Panel **c** shows staining pattern in glioma region characterized by intense SH activity originating from cell clusters (TAMRA-FP hotspot clusters). Note detectable expression of CD68 in the glioma and its absence from other brain regions (**a**). Note poor co-localization of CD68-positive cells with TAMRA-FP hotspots (**b**) or with TAMRA-FP hotspot clusters (**c**). Primary antibody rabbit anti-CD163 (Abcam, cat# ab182422), dilution 1:75, secondary antibody Goat anti-rabbit IgG-Alexa Fluor 647 conjugate, dilution 1:100. Sections were from female rat 11. Scale bars: 1 mm in a, 20 µm in b-c. Images were adjusted for brightness and contrast.

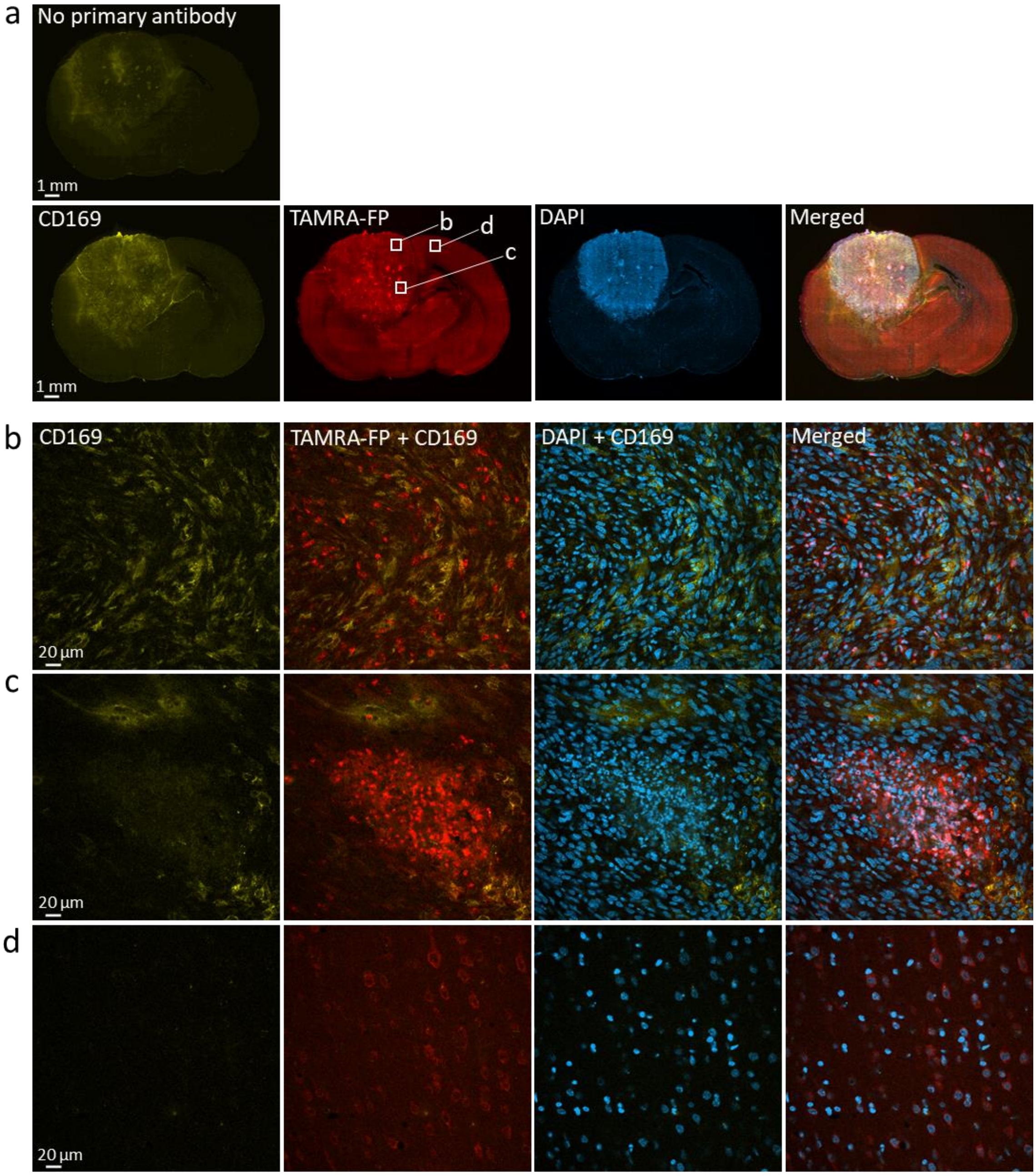


**Figure S21. Confocal imaging of SH activity in relation to CD169, a marker for macrophages.** Sections went through the tissue-ABPP protocol to label SHs (red) and were thereafter immunostained for CD169 (yellow), followed by DAPI staining to visualize nuclei (blue). Panel **a** shows overall staining pattern throughout the coronal section plane. A control section undergoing identical staining protocol with no primary antibody is illustrated at top. Panel **b** shows staining pattern in glioma region characterized by intense SH activity originating from individual cells (TAMRA-FP hotspots). Panel **c** shows staining pattern in glioma region characterized by intense SH activity originating from cell clusters (TAMRA-FP hotspot clusters). Panel **d** shows staining pattern in healthy brain (cortex). Note detectable expression of CD169 in the glioma (**a**) and its poor co-localization with TAMRA-FP hotspots (**b**). Note lack of CD169 from TAMRA-FP hotspot clusters (**c**). CD169 is undetectable in control cortical region (**d**). Primary antibody mouse anti-CD169 (Bio-Rad, cat# MCA343GA), dilution 1:25, secondary antibody donkey anti-mouse IgG-Alexa Fluor 647 conjugate, dilution 1:100. Sections were from female rat 11. Scale bars: 1 mm in a, 20 µm in b-d. Images were adjusted for brightness and contrast.

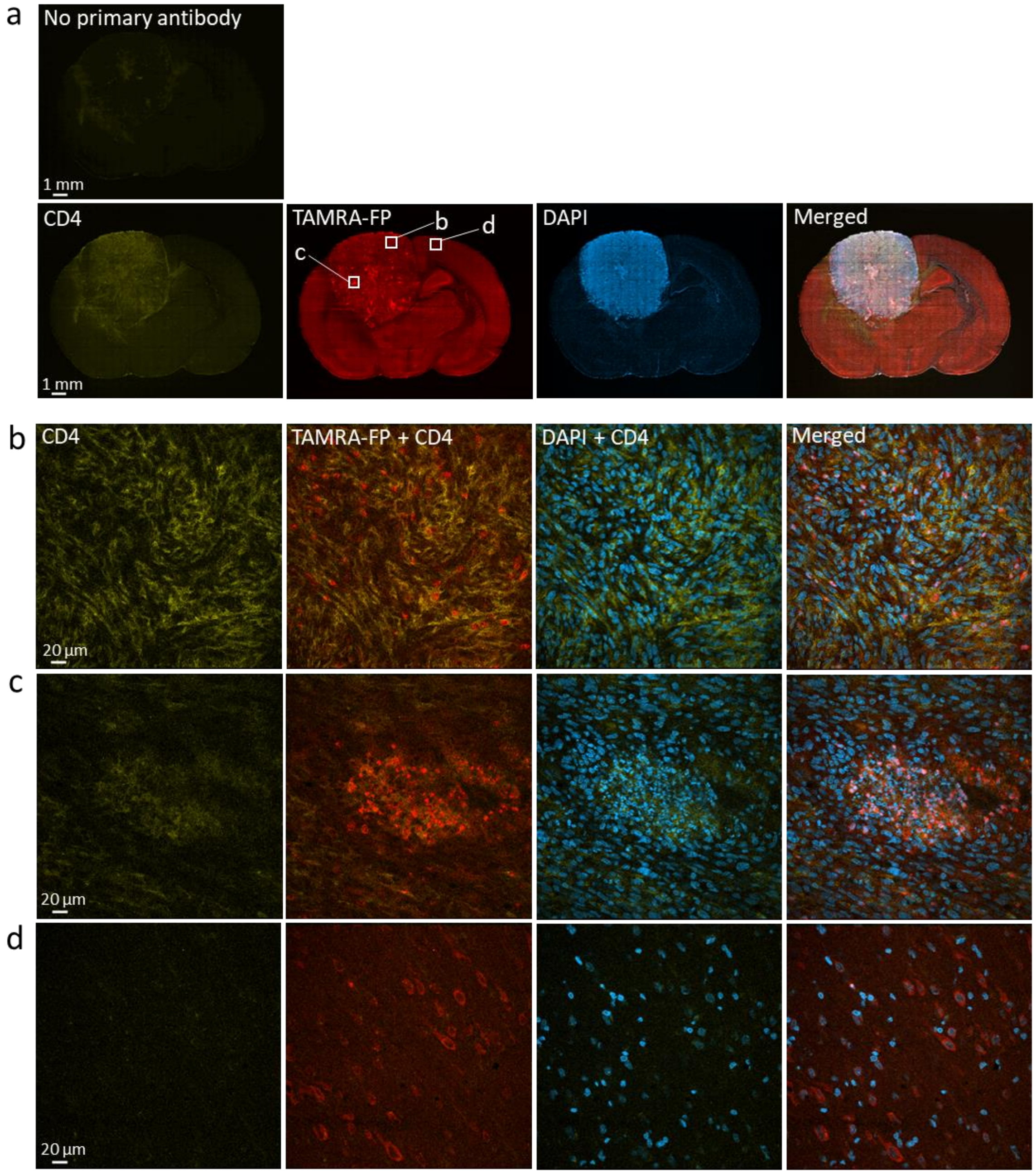


**Figure S22. Confocal imaging of SH activity in relation to T cell marker CD4.** Sections went through the tissue-ABPP protocol to label SHs (red) and were thereafter immunostained for CD4 (yellow), followed by DAPI staining to visualize nuclei (blue). Panel **a** shows overall staining pattern throughout the coronal section plane. A control section undergoing identical staining protocol with no primary antibody is illustrated at top. Panel **b** shows staining pattern in glioma region characterized by intense SH activity originating from individual cells (TAMRA-FP hotspots). Panel **c** shows staining pattern in glioma region characterized by intense SH activity originating from cell clusters (TAMRA-FP hotspot clusters). Panel **d** shows staining pattern in healthy brain (cortex). Note detectable expression of CD4 in the glioma (**a**) and its absence from other brain regions (**a, d**). Note robust CD4 expression and its partial co-localization with TAMRA-FP hotspots (**b**). Note also CD expression in area with TAMRA-FP hotspot clusters and its partial co-localization with the cluster (**c**). Primary antibody mouse anti-CD4 (OX-35, Abcam, cat# ab33775), dilution 1:100, secondary antibody donkey anti-mouse IgG-Alexa Fluor 647 conjugate, dilution 1:100. Sections were from female rat 11. Scale bars: 1 mm in a, 20 µm in b-d. Images were adjusted for brightness and contrast.

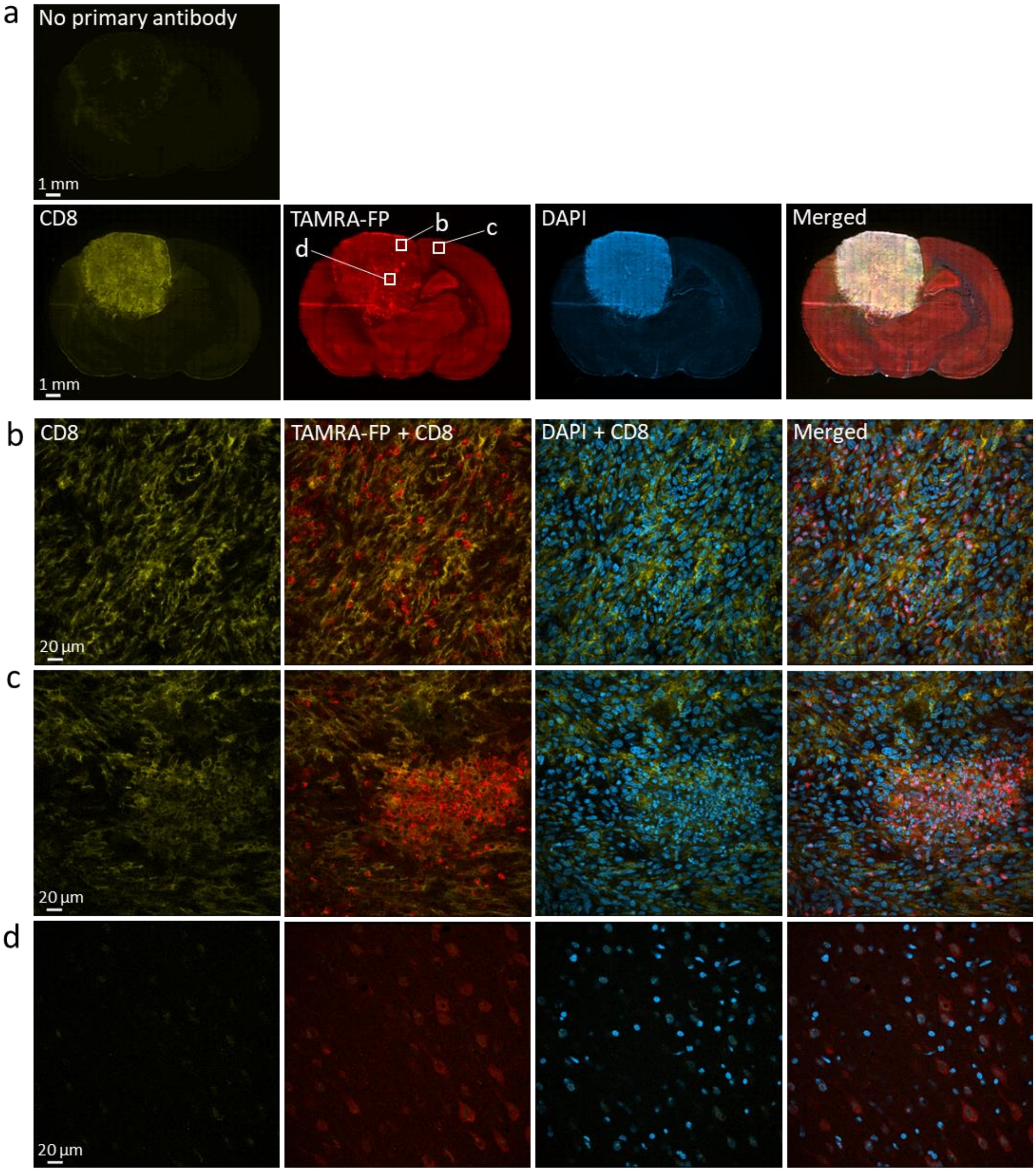


**Figure S23. Confocal imaging of SH activity in relation to T cell marker CD8.** Sections went through the tissue-ABPP protocol to label SHs (red) and were thereafter immunostained for CD8 (yellow), followed by DAPI staining to visualize nuclei (blue). Panel **a** shows overall staining pattern throughout the coronal section plane. A control section undergoing identical staining protocol with no primary antibody is illustrated at top. Panel **b** shows staining pattern in glioma region characterized by intense SH activity originating from individual cells (TAMRA-FP hotspots). Panel **c** shows staining pattern in glioma region characterized by intense SH activity originating from cell clusters (TAMRA-FP hotspot clusters). Panel **d** shows staining pattern in healthy brain (cortex). Note prominent expression of CD8 throughout the glioma (**a**) and its absence from healthy brain (**a, d**). Note CD8-positive cells and their partial localization to TAMRA-FP hotspots (**b**). Note abundance of CD8-positive cells in the region of TAMRA-FP hotspot clusters, showing partial co-localization with TAMRA-FP signal (**c**). Primary antibody mouse anti-CD8 alpha (OX-8, Abcam, cat# ab33786), dilution 1:500, secondary antibody donkey anti-mouse IgG-Alexa Fluor 647 conjugate, dilution 1:100. Sections were from female rat 11. Scale bars: 1 mm in a, 20 µm in b-d. Images were adjusted for brightness and contrast.

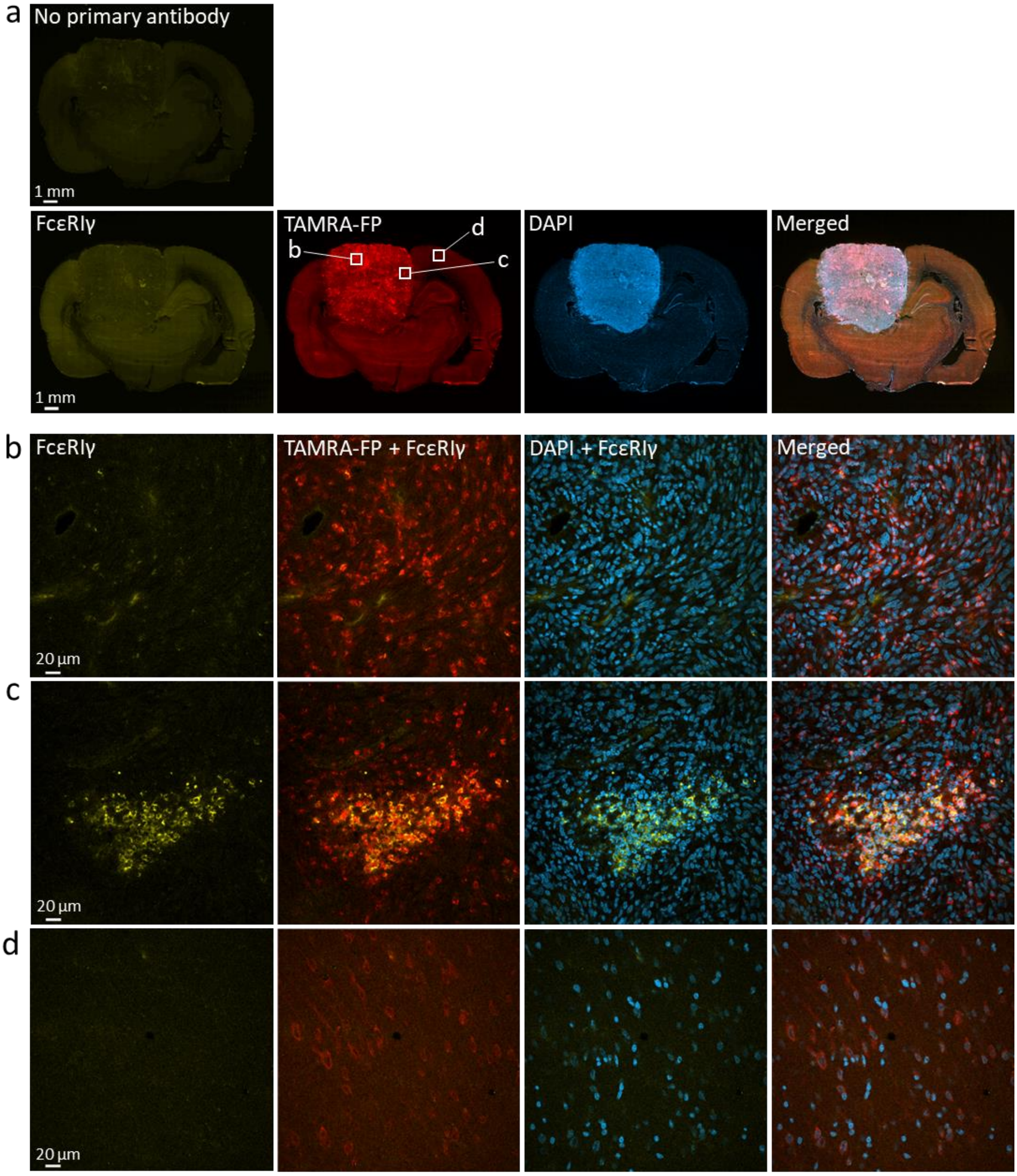


**Figure S24. Confocal imaging of SH activity in relation to FcεRIγ, a marker for mast cells, eosinophils, basophils and monocytes.** Sections went through the tissue-ABPP protocol to label SHs (red) and were thereafter immunostained for FcεRIγ (yellow), followed by DAPI staining to visualize nuclei (blue). Panel **a** shows overall staining pattern throughout the coronal section plane. A control section undergoing identical staining protocol with no primary antibody is illustrated at top. Panel **b** shows staining pattern in glioma region characterized by intense SH activity originating from individual cells (TAMRA-FP hotspots). Panel **c** shows staining pattern in glioma region characterized by intense SH activity originating from cell clusters (TAMRA-FP hotspot clusters). Panel **d** shows staining pattern in healthy brain (cortex). Note scarce expression of FcεRIγ in the glioma (**a**) and that this marker is practically absent from TAMRA-FP hotspots (**b**). Note enrichment of FcεRIγ and its co-localization with TAMRA-FP hotspot clusters (**c**). FcεRIγ is undetectable in control cortical region (**d**). Primary antibody mouse anti-FcεRIγ (F-1, Santa Cruz, cat# sc-390221), dilution 1:100, secondary antibody donkey anti-mouse IgG-Alexa Fluor 647 conjugate, dilution 1:100. Sections were from male rat 31. Scale bars: 1 mm in a, 20 µm in b-d. Images were adjusted for brightness and contrast.

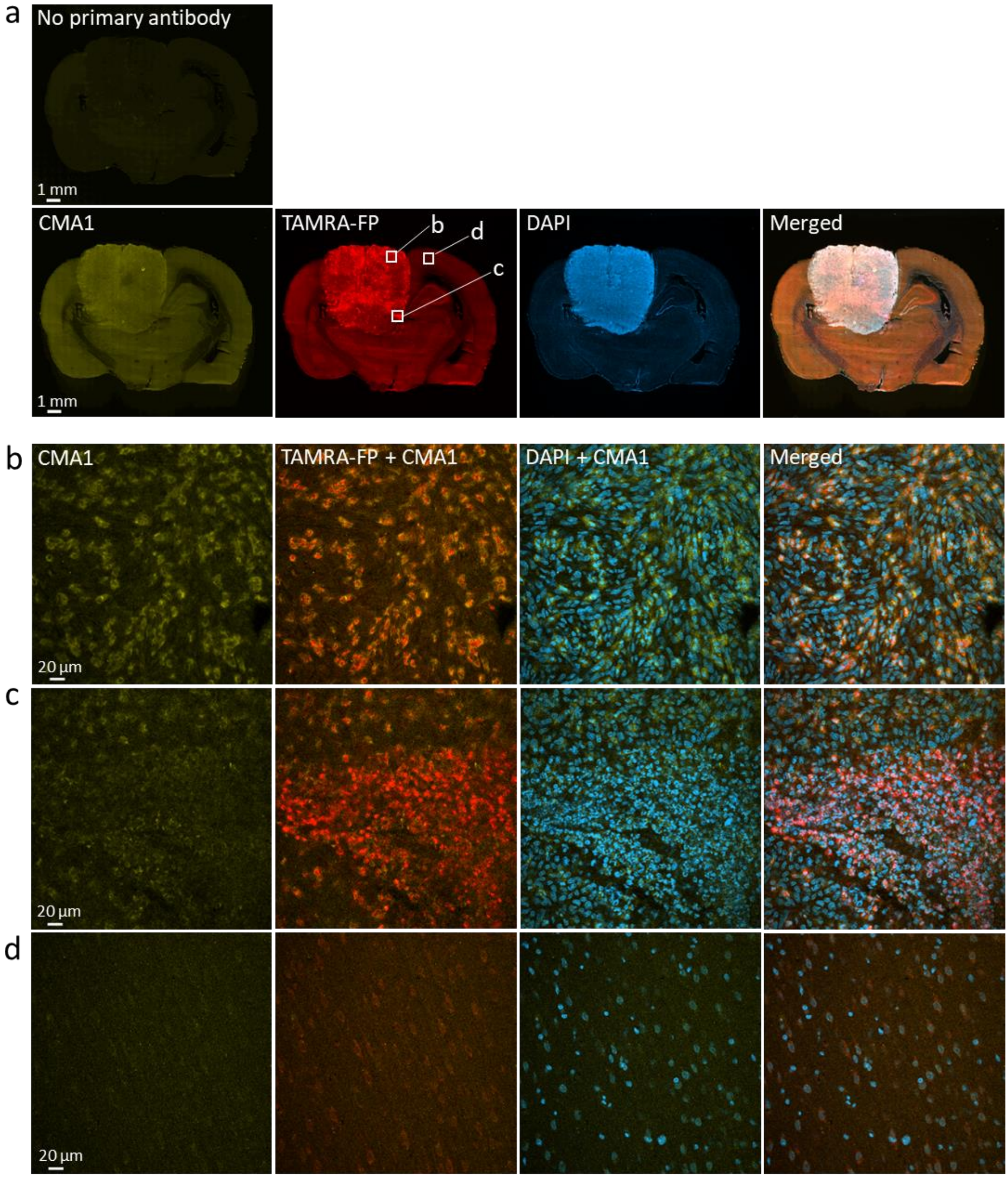


**Figure S25. Confocal imaging of SH activity in relation to chymase (CMA1), a marker for mast cells.** Sections went through the tissue-ABPP protocol to label SHs (red) and were thereafter immunostained for CMA1 (yellow), followed by DAPI staining to visualize nuclei (blue). Panel **a** shows overall staining pattern throughout the coronal section plane. A control section undergoing identical staining protocol with no primary antibody is illustrated at top. Panel **b** shows staining pattern in glioma region characterized by intense SH activity originating from individual cells (TAMRA-FP hotspots). Panel **c** shows staining pattern in glioma region characterized by intense SH activity originating from cell clusters (TAMRA-FP hotspot clusters). Panel **d** shows staining pattern in healthy brain (cortex). Note faint expression of CMA1 in the glioma (**a**). Note CMA1-positive cells in the region of TAMRA-FP hotspots with notable co-localization with the TAMRA-FP signal (**b**). Note absence of CMA1-positive cells from TAMRA-FP hotspot clusters (**c**). CMA1 is undetectable in control cortical region (**d**). Primary antibody rabbit anti-Chymase 1 (Biomatic, CAU22217), dilution 1:500, secondary antibody goat anti-rabbit IgG-Alexa Fluor 647 conjugate, dilution 1:100. Sections were from male rat 31. Scale bars: 1 mm in a, 20 µm in b-d. Images were adjusted for brightness and contrast.

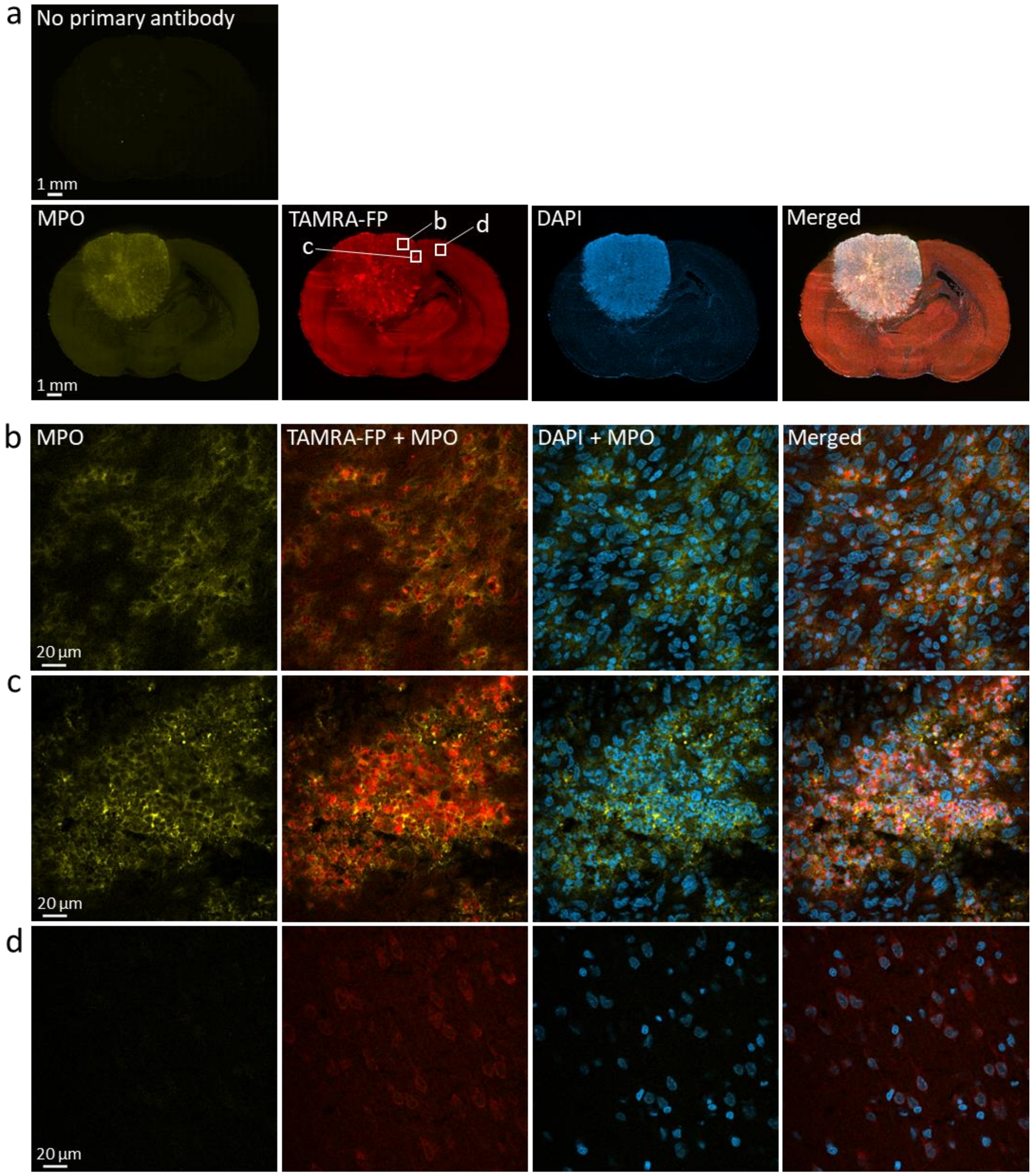


**Figure S26. Confocal imaging of SH activity in relation to myeloperoxidase (MPO), a marker for neutrophils.** Sections went through the tissue-ABPP protocol to label SHs (red) and were thereafter immunostained for MPO (yellow), followed by DAPI staining to visualize nuclei (blue). Panel **a** shows overall staining pattern throughout the coronal section plane. A control section undergoing identical staining protocol with no primary antibody is illustrated at top. Panel **b** shows staining pattern in glioma region characterized by intense SH activity originating from individual cells (TAMRA-FP hotspots). Panel **c** shows staining pattern in glioma region characterized by intense SH activity originating from cell clusters (TAMRA-FP hotspot clusters). Panel **d** shows staining pattern in healthy brain (cortex). Note detectable expression of MPO in the glioma (**a**). Note MPO-positive cells in the region of TAMRA-FP hotspots and marked co-localization of MPO with the TAMRA-FP signal (**b**). Note abundance of MPO-positive cells within TAMRA-FP hotspot clusters, as well as close match of MPO staining with TAMRA-FP hotspot clusters (**c**). MPO is not visible in control cortical region (**d**). Primary antibody rabbit anti-MPO (Abcam, cat# ab9535), dilution 1:25, secondary antibody Goat anti-rabbit IgG-Alexa Fluor 647 conjugate, dilution 1:100. Sections were from female rat 11. Scale bars: 1 mm in a, 20 µm in b-d. Images were adjusted for brightness and contrast.

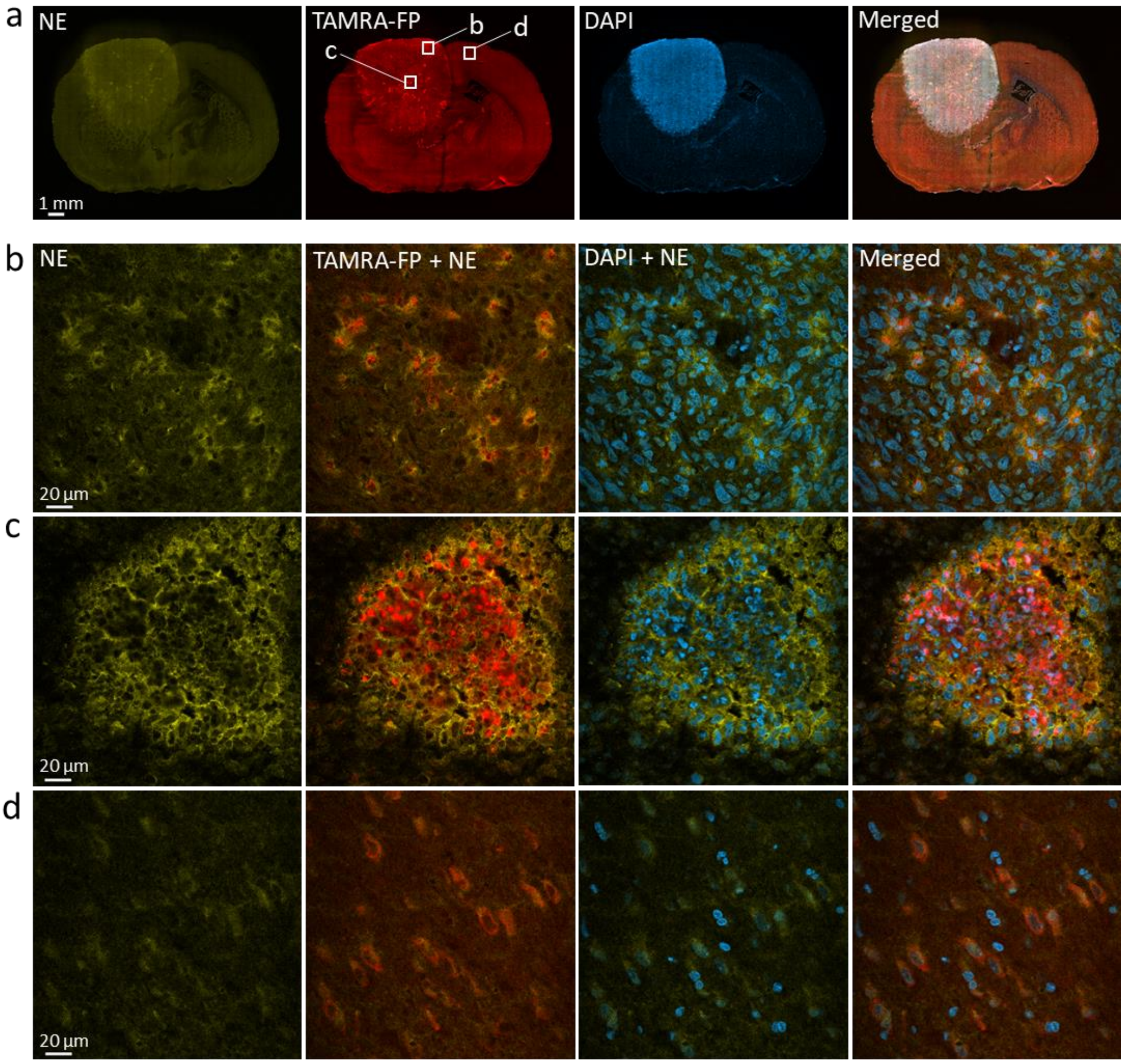


**Figure S27. Confocal imaging of SH activity in relation to neutrophil elastase (NE).** Sections went through the tissue-ABPP protocol to label SHs (red) and were thereafter immunostained for NE (yellow), followed by DAPI staining to visualize nuclei (blue). Panel **a** shows overall staining pattern throughout the coronal section plane. A control section undergoing identical staining protocol with no primary antibody was not available for this experiment. Panel **b** shows staining pattern in glioma region characterized by intense SH activity originating from individual cells (TAMRA-FP hotspots). Panel **c** shows staining pattern in glioma region characterized by intense SH activity originating from cell clusters (TAMRA-FP hotspot clusters). Panel **d** shows staining pattern in healthy brain (cortex). Note detectable expression of NE throughout the glioma (**a**). Note in particular NE-positive cells in the region of TAMRA-FP hotspots and close match of NE-positive cells with the TAMRA-FP signal (**b**). Note strong NE-immunostaining of the TAMRA-FP hotspot clusters and close match of NE staining within the TAMRA-FP hotspot cluster (**c**). NE is not visible in control cortical region (**d**). Primary antibody rabbit anti-NE (Abcam, cat# ab21595), dilution 1:500, secondary antibody Goat anti-rabbit IgG-Alexa Fluor 647 conjugate, dilution 1:100. Sections were from female rat 11. Scale bars: 1 mm in a, 20 µm in b-d. Images were adjusted for brightness and contrast.

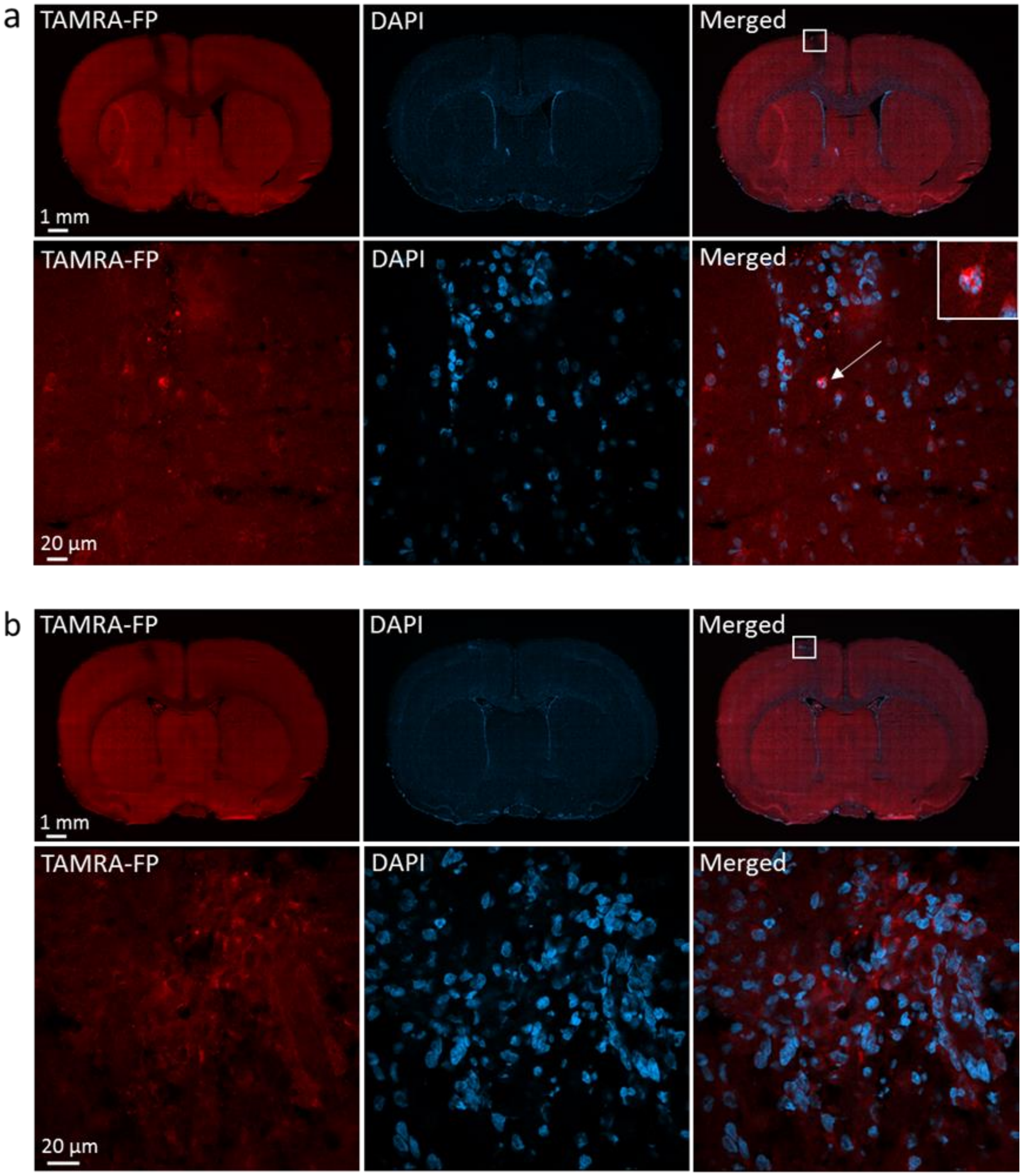


**Figure S28. TAMRA-FP signal at the site of injection in sham-operated animals.** Sections covering the site of sham-injection (tumor cell medium only) in male rat 23 (**a**) and female rat 9 (**b**) went through the tissue-ABPP protocol to label SHs (red), followed by DAPI staining to visualize nuclei (blue). As a rule, multi-nucleated cells with TAN-like morphology were not evident with the exception of one particular cell showing also TAMRA-FP hotspot characteristics. Scale bar 1 mm (upper panels) and 20 µm (lower panels). Scale bars: 1 mm in a, 20 µm in b-d. Images were adjusted for brightness and contrast.

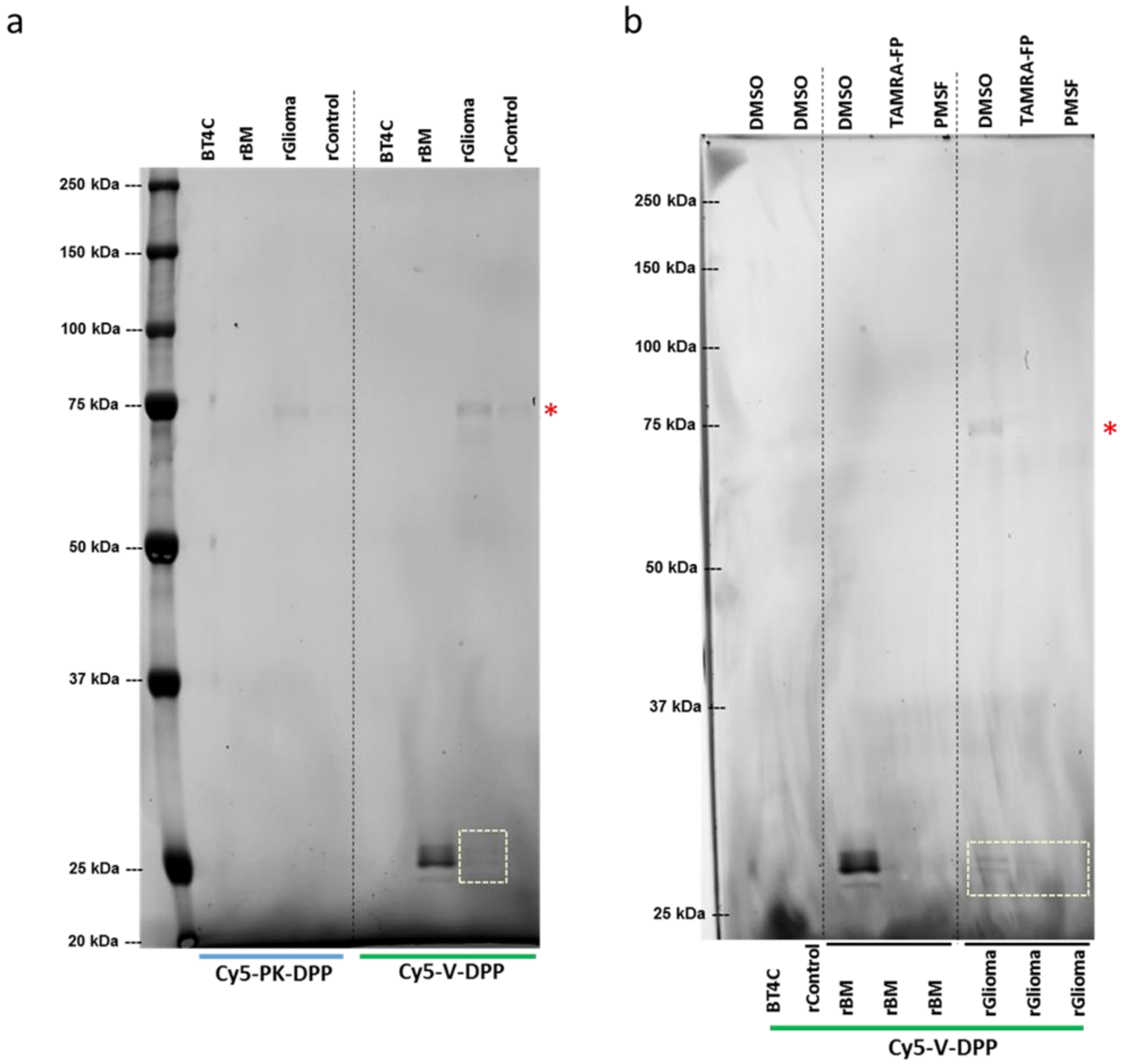


**Figure S29. Gel-ABPP of rat glioma proteomes using Cy5-labeled serine protease activity probes PK-DPP and V-DPP.** Lysates of BT4C tumor cells, bone-marrow-derived mononuclear cells (rBM) or homogenates of glioma and control brain (all 4 mg/ml) were directly labelled by the indicated Cy5-probes (**a**) or first pretreated for 1h with DMSO or with the SH inhibitors TAMRA-FP (1 µM) or PMSF (1 mM) (**b**), after which the proteomes were labelled by the indicated Cy5-probes for 1h (probe concentration 1 µM). The reaction was quenched, 30 µg protein was loaded per lane, proteins were separated by SDS-PAGE, followed by in-gel fluorescence imaging. MW markers are indicated at left. In **a**, note that the trypsin-preferring probe PK-DPP weakly detects a single ~75 kDa band with unknown identity (red asterisk) in samples of glioma and control brain and that a band of similar size is also weakly recognized by V-DPP. Note that the elastase-preferring probe V-DPP detects 3-4 bands at the ~25 kDa range in rBM and two bands of similar size faintly in the glioma sample (circled by the dotted line). In **b** note that V-DPP labeling of the ~25 kDa protein bands in rBM and glioma (circled by the dotted line) is sensitive to the inhibitors, indicating mutual SH targets for these compounds. Images were adjusted for brightness and contrast. Lanes not related to these probes were removed.

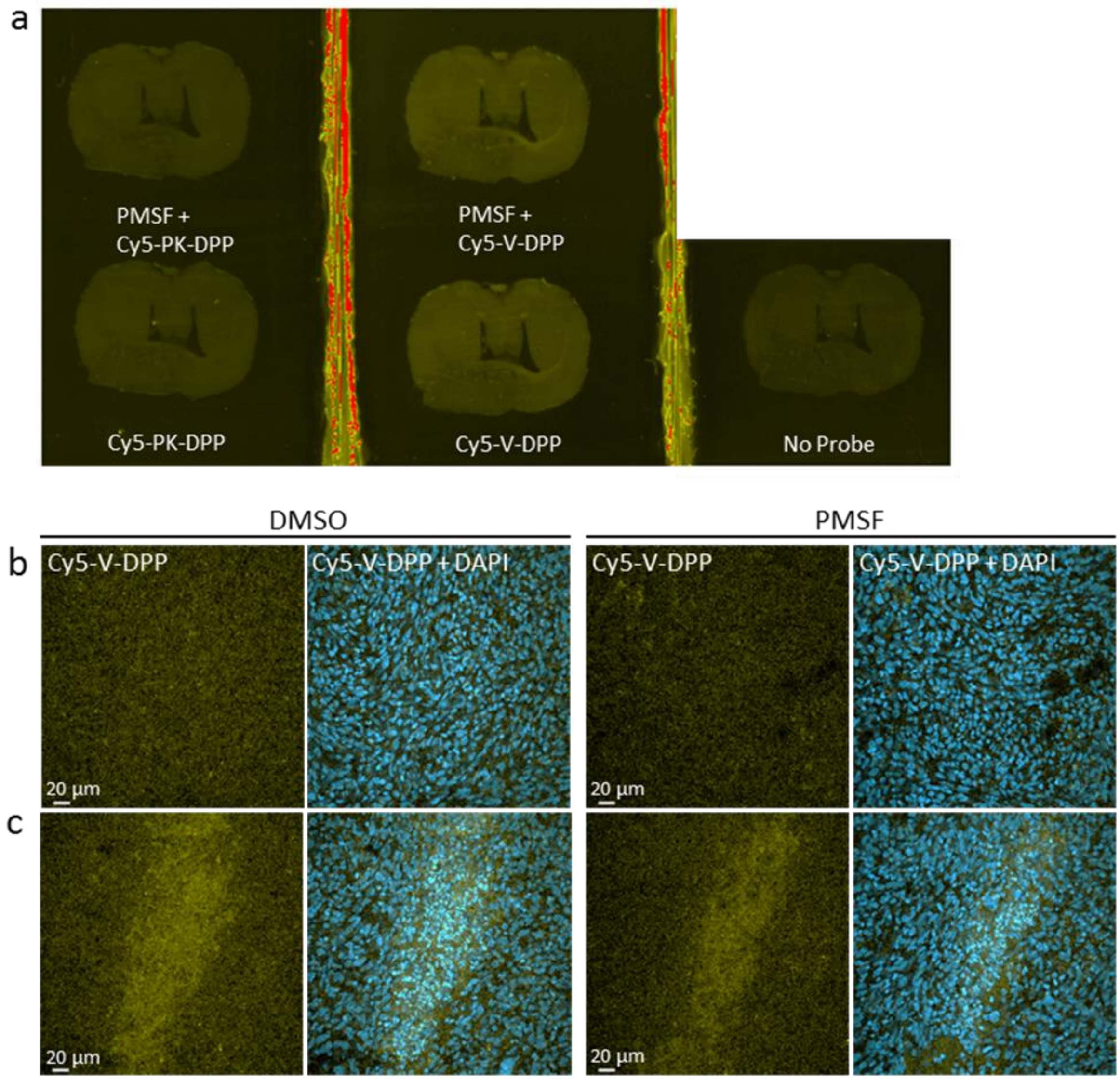


**Figure S30. Tissue-ABPP of glioma sections using Cy5-labeled activity probes PK-DPP and V-DPP.** Sections were pretreated with solvent (DMSO) or PMSF (1 mM) for 1h at RT, after which they went through the tissue-ABPP protocol using the indicated probes instead of TAMRA-FP (yellow, 0.5 µM final probe concentration), followed by DAPI staining to visualize nuclei (blue). Panel **a** shows overall staining pattern throughout the coronal section plane, imaged using BioRad gel scanner (Cy5 window). A control section undergoing identical protocol with no probe is shown for comparison. Note weak and largely PMSF-resistant signal for both probes. Panel **b** compares Cy5-probe staining pattern between DMSO- and PMSF-pretreated sections in glioma region of TAMRA-FP hotspots. Panel **c** compares Cy5-probe staining pattern between DMSO- and PMSF-pretreated sections in glioma region of TAMRA-FP hotspot clusters. Note largely PMSF-resistant binding of both probes throughout these regions (**b**-**c**). Images were adjusted for brightness and contrast.

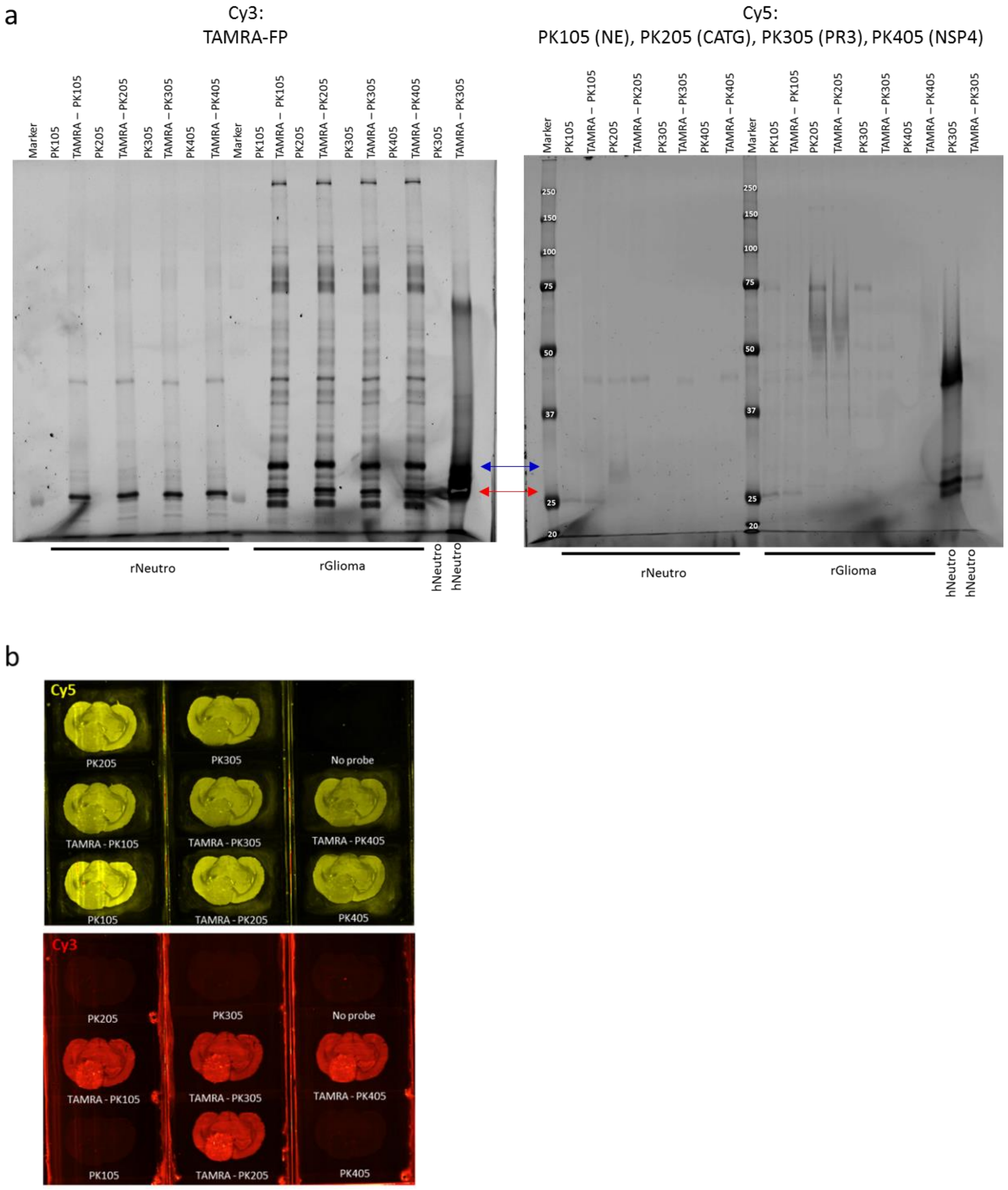


**Figure S31. ABPP of rat neutrophil and glioma samples using Cy5-labeled neutrophil serine protease (NSP) probes in combination with TAMRA-FP.** In **a**, rat neutrophil lysates (rNeutro, 0.1 mg/ml) or glioma homogenates (1 mg/ml) were treated for 1h with DMSO or with TAMRA-FP (1 µM), after which the proteomes were incubated with the indicated Cy5-probes for 1h (probe concentration 1 µM). Human neutrophil lysate (hNeutro, 0.1 mg/ml) was included as a positive control. The reaction was quenched, 0.7 or 7 µg protein (lysates and glioma homogenate, respectively) was loaded per lane, proteins separated by SDS-PAGE, followed by in-gel fluorescence imaging by BioRad gel scanner using Cy3-window for TAMRA-FP and Cy5-window for the NSP probes. Images of the same gel showing fluorescence in both windows with MW markers indicated in the Cy5-window. Note that TAMRA-FP strongly labels a ~25 kDa band in rat neutrophils (red double-headed arrow) and that band of similar size is also strongly labeled in glioma. Note that the neutrophil ~25 kDa band is not readily detected by any of the NPS probes. Note absence of the prominent ~30 kDa glioma band (blue double-headed arrow) from rat neutrophils. Note also that under the conditions employed, PK305 strongly labels several bands in the 25-30 kDa range in human neutrophils and that this labeling is sensitive to TAMRA-FP, indicating that the activity probes label the same SH targets. **b.** Tissue-ABPP of glioma sections using Cy5-labeled NSP probes in combination with TAMRA-FP. Sections were pretreated with DMSO or TAMRA-FP (1 µM) for 1h at RT, after which they went through the tissue-ABPP protocol using the indicated probes (0.2 µM final concentration) instead of TAMRA-FP. The upper and lower panels show Cy5- and Cy3-probe labeling pattern, respectively, throughout the coronal section plane, imaged using BioRad gel scanner. A control section undergoing identical protocol with no probe is shown for comparison. Note robust and largely TAMRA-FP-insensitive binding of the Cy5-probes throughout the section. Images were adjusted for brightness and contrast.

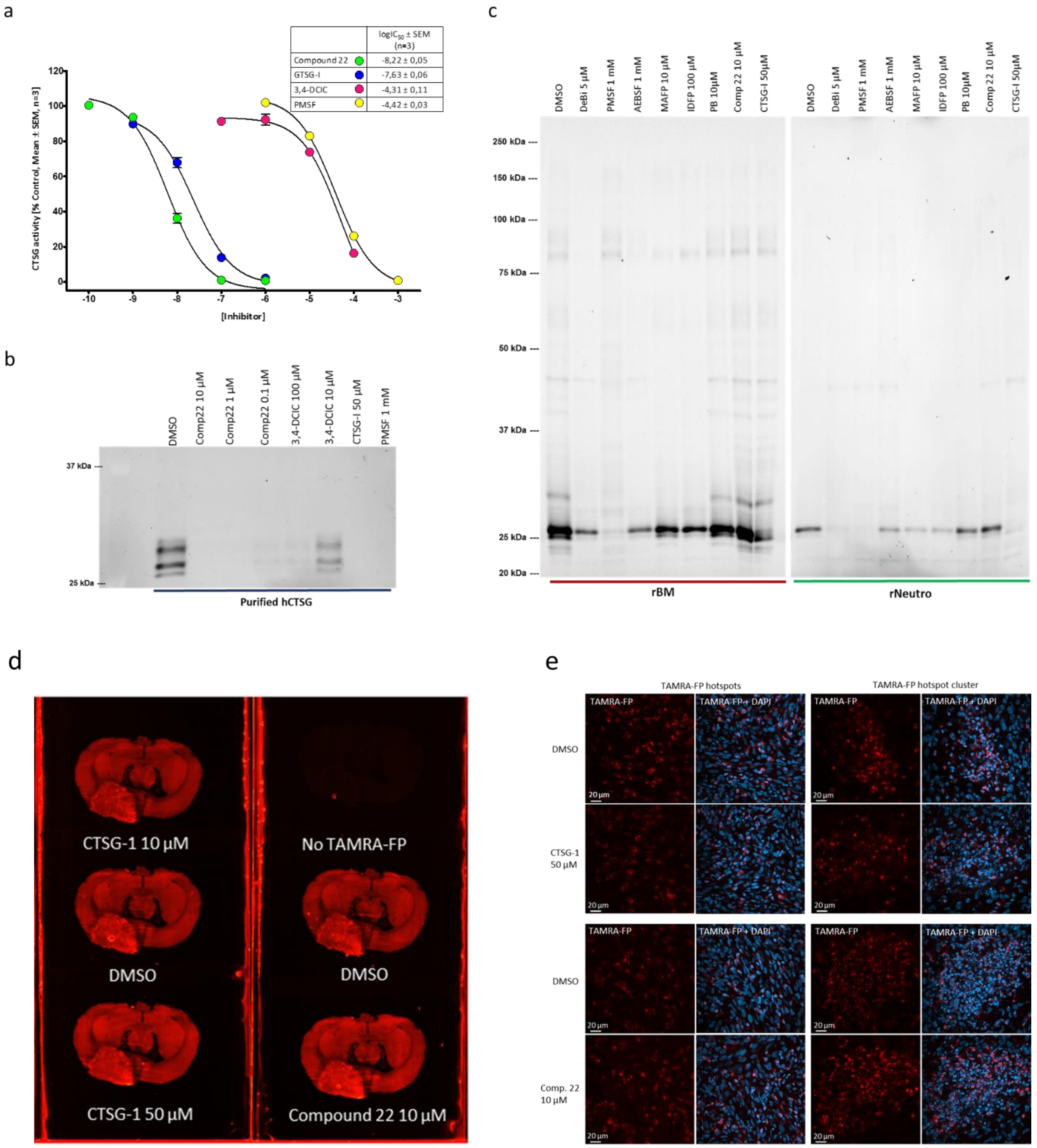


**Figure S32. Inhibitor profiles of human cathepsin G (hCTSG) and the prominent 25-30 kDa SH bands in rat bone-marrow-derived mononuclear cells and neutrophils.** **a**) Dose-response curves and potencies (log$IC_{50}$ values) for Compound 22 (Cyclo-GTCnXSDPPICFPN), GTSG-Inhibitor (GTSG-I), 3,4-dichloroisocoumarin (3,4-DCIC) and PMSF in inhibiting purified human neutrophil CTSG (Calbiochem Cat# 219373, 100 µU/well). hCTSG-mediated hydrolysis of the chromogenic substrate N-succinyl-Ala-Ala-Pro-Phe-pNA (100 µM) generates 4-nitroaniline whose absorbance was kinetically monitored at $\lambda_{405\ nm}$. The assay contained additionally 0.1 % (w/v) BSA and 1% (v/v) DMSO. **b)** In gel-ABPP, purified hCTSG migrates as four separate bands in the 25-30 kDa range. Note potent inhibition of hCTSG by Compound 22 in line with the outcome of the substrate-based activity assay. **c)** Comparative inhibitor profiling of rat bone-marrow-derived mononuclear cells (rBM) and neutrophils (rNeutro). Note that in contrast to PMSF and 3,4-DCIC, Compound 22 does not inhibit 25-30 kDa SH bands in rBM or neutrophils. Inhibitor profiles of the 25-30 kDa SH bands in rBM (1 mg/ml) and the 25 kDa band in neutrophils (0.1 mg/ml) show similar pharmacology, taken into account 10-fold difference in protein-inhibitor stoichiometry (only limited amounts of neutrophils were available for these experiments). **d-e)** Competitive tissue-ABPP of glioma sections pretreated with GTSG inhibitors. Sections were pretreated with DMSO or the indicated concentrations of the CTSG inhibitors for 1h at RT, after which they went through the tissue-ABPP protocol using TAMRA-FP (0.5 µM final concentration). Panel **d**) shows TAMRA-FP labeling pattern throughout the coronal section plane, imaged using BioRad gel scanner and panel **e)** shows confocal fluorescence images of regions with TAMRA-FP hotspots and TAMRA-FP hotspot clusters. Note insensitivity of TAMRA-FP signal throughout hotspots or hotspot clusters towards the CTSG inhibitors. Abbreviations: DeBi, desthiobiotin-FP; PB, palmostatin B. Images were adjusted for brightness and contrast.

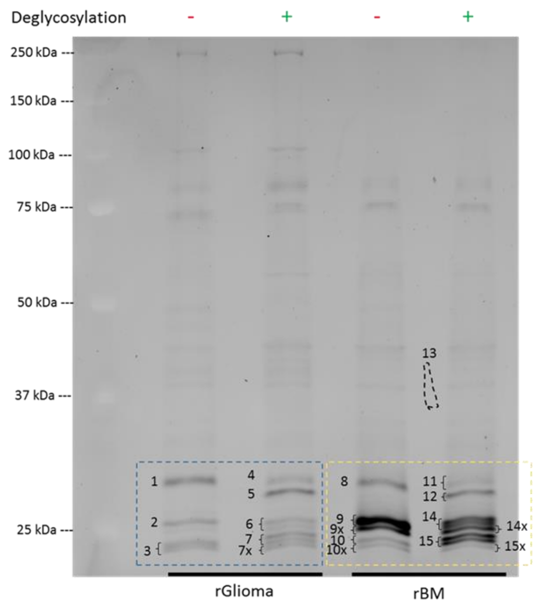


| Sample | rGlioma |
|---|---|
| 1 | Esd |
| 2 | Pafah1b3 , Mcpt1, Clpp, Ctsg, Elane, Prtn3 |
| 3 | Lypla1, Lypla2, Ctsg |
| 4 | Esd |
| 5 | Pafah1b2 |
| 6 | Olah, Pafah1b3, Clpp, Ctsg, Elane |
| 7 | Lypla1, Ctsg, Elane |
| 7x | Lypla1, Ctsg, Elane, Prtn3, Lypla2 |

| Sample | rBM |
|---|---|
| 8 | Mcpt10, Esd, Cma1, Elane, Ctsg |
| 9 | Iah1, Pafah1b3, Mcpt1, Elane, Ctsg, Prtn3 |
| 9x | Pafah1b3, Mcpt1, Elane, Ctsg, Prtn3 |
| 10 | Elane, Ctsg, Prtn3 |
| 10x | Esd, Ctsg, Prtn3, Lypla1, Lypla2 |
| 11 | Mcpt10, Esd, Ppt1 |
| 12 | Mcpt10, Iah1, Cma1, Ctsg |
| 14 | Cma1, Pafah1b3, Mcpt1, Clpp, Elane, Ctsg, Prtn3 |
| 14x | Pafah1b3, Mcpt1, Clpp, Elane, Ctsg, Prtn3 |
| 15 | Elane, Ctsg, Prtn3 |
| 15x | Cma1, Elane, Ctsg, Prtn3 |

Abbreviations: Clpp, ATP-dependent Clp protease proteolytic subunit; Cma1, Chymase; Ctsg, Cathepsin G; Elane, Elastase 2, neutrophil; Esd, S-formylglutathione hydrolase; Iah1, Isoamyl acetate-hydrolyzing esterase 1 homolog; Lypla1, Acyl-protein thioesterase 1; Lypla2, Acyl-protein thioesterase 2; Mcpt1, Mast cell protease 1; Mcpt10, Granzyme-like protein 2; Olah, S-acyl fatty acid synthase thioesterase, medium chain; Pafah1b2, Platelet-activating factor acetylhydrolase IB subunit beta; Pafah1b3, Platelet-activating factor acetylhydrolase IB subunit gamma; Ppt1, Palmitoyl-protein thioesterase 1; Prtn3, Proteinase 3

**Figure S33. LC-MS/MS-based identification of the ~25-30 kDa SH bands in rat glioma and bone marrow-derived mononuclear cells.** Gel-ABPP was conducted using rat glioma homogenates (rGlioma) or lysates of rat bone marrow-derived mononuclear cells (rBM) as detailed in Materials and Methods. To facilitate SDS-PAGE separation of proteins with similar size, the proteomes (4 mg/ml) were deglycosylated (+) or underwent control treatment (-) prior to TAMRA-FP labeling. For deglycosylation, samples were treated with Protein Deglycosylation Mix II (New England Biolabs, Cat#P6044S) for 1h at RT as per kit instructions, after which TAMRA-FP labeling was conducted using routine gel-ABPP protocol. Following in-gel fluorescence imaging, gel-pieces encompassing the SH bands of interest (numbered 1-15x) were cut and subjected to LC-MS/MS analysis, as described in Materials and Methods. Gel-pieces marked with x were cut as one sample containing also the numbered gel-piece and split thereafter, yielding two separate samples that were subjected to LC-MS/MS. The SHs identified from the glioma samples (1-7x) are listed in the middle (blue-white table) and those identified from rBM (8-15x) at right (yellow-white table). Note presence of the NSPs cathepsin G, elastase 2 and proteinase 3 in the ~25 kDa gel-pieces of both proteomes. Note also presence of platelet-activating factor acetylhydrolases 1b2 and 1b3 (PAFAH1b3 and PAFAH1b2) in the 25-30 kDa gel-pieces of both proteomes. The LC-MS/MS data as a whole showing all identified proteins and additionally data on BT4C glioma cells are available in Supplementary File 1.

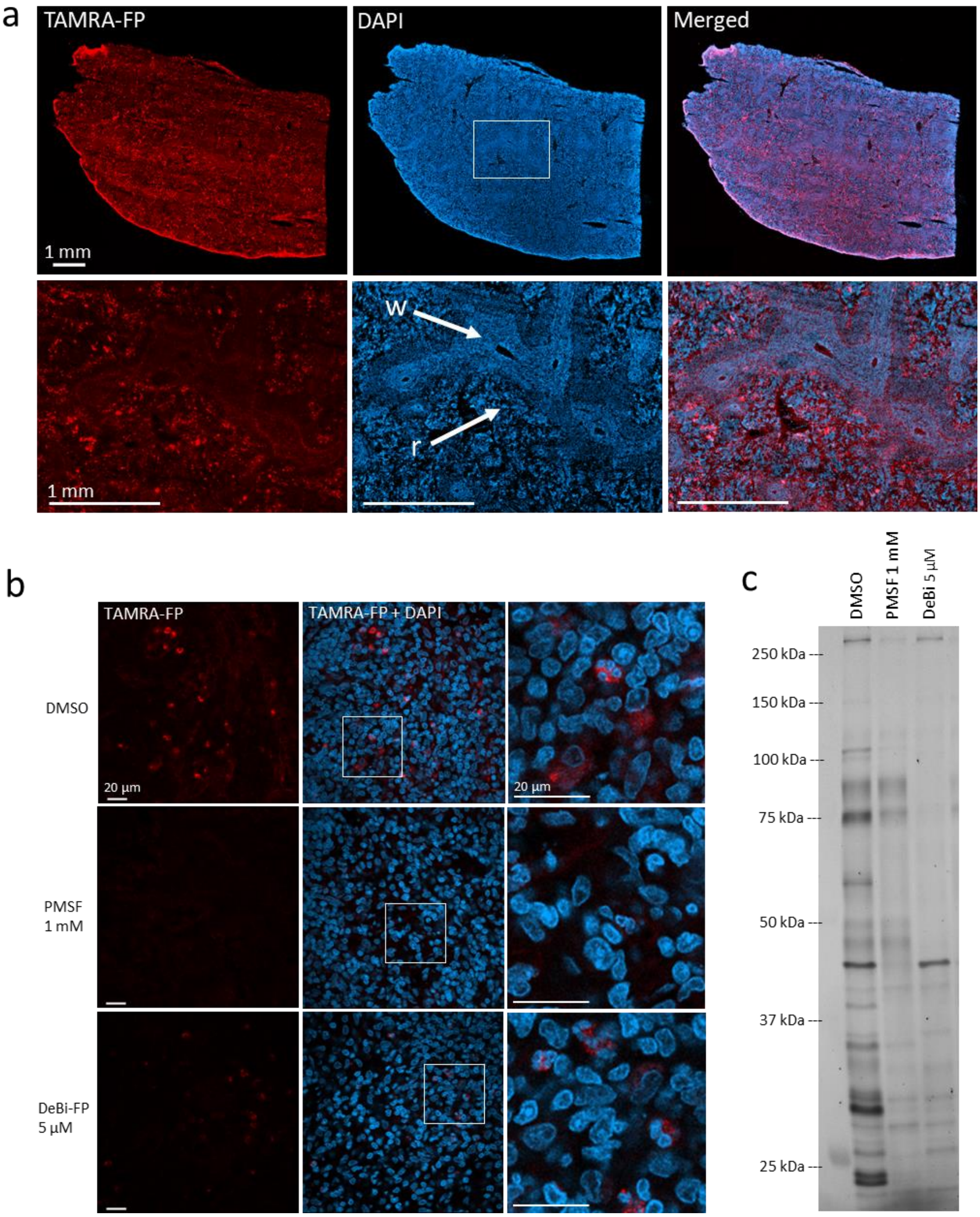


**Figure S34. High-resolution imaging of TAMRA-FP hotspots and their inhibitor sensitivity in rat spleen.** In **a**, sections went through the tissue-ABPP protocol to label SHs (red), followed by DAPI staining to visualize nuclei (blue). Note presence of TAMRA-FP hotspots in red pulp (r) and absence of TAMRA-FP hotspots from white pulp (w). In **b**, sections were pretreated with DMSO or with the SH inhibitors PMSF (1 mM) or deshiobiotin-FP (DeBi-FP, 5 µM) for 1h at RT, after which they went through the tissue-ABPP protocol to label SHs (red), followed by DAPI staining to visualize nuclei (blue). Red pulp regions of intense TAMRA-FP labeling were imaged. Note presence of multi-nucleated cells in regions of TAMRA-FP hotspots. Note that throughout the examined regions, TAMRA-FP labeling is sensitive to the inhibitors. Gel-ABPP (**c**) using spleen homogenate was run for comparison to visualize TAMRA-FP labeled SH bands and their sensitivity to inhibitors. Note that PMSF is more efficient than DeBi-FP in blocking TAMRA-FP labeling in both tissue- and gel-based ABPP. Scale bars 1 mm (**a**) and 20 µm (**b**). Images were adjusted for brightness and contrast.

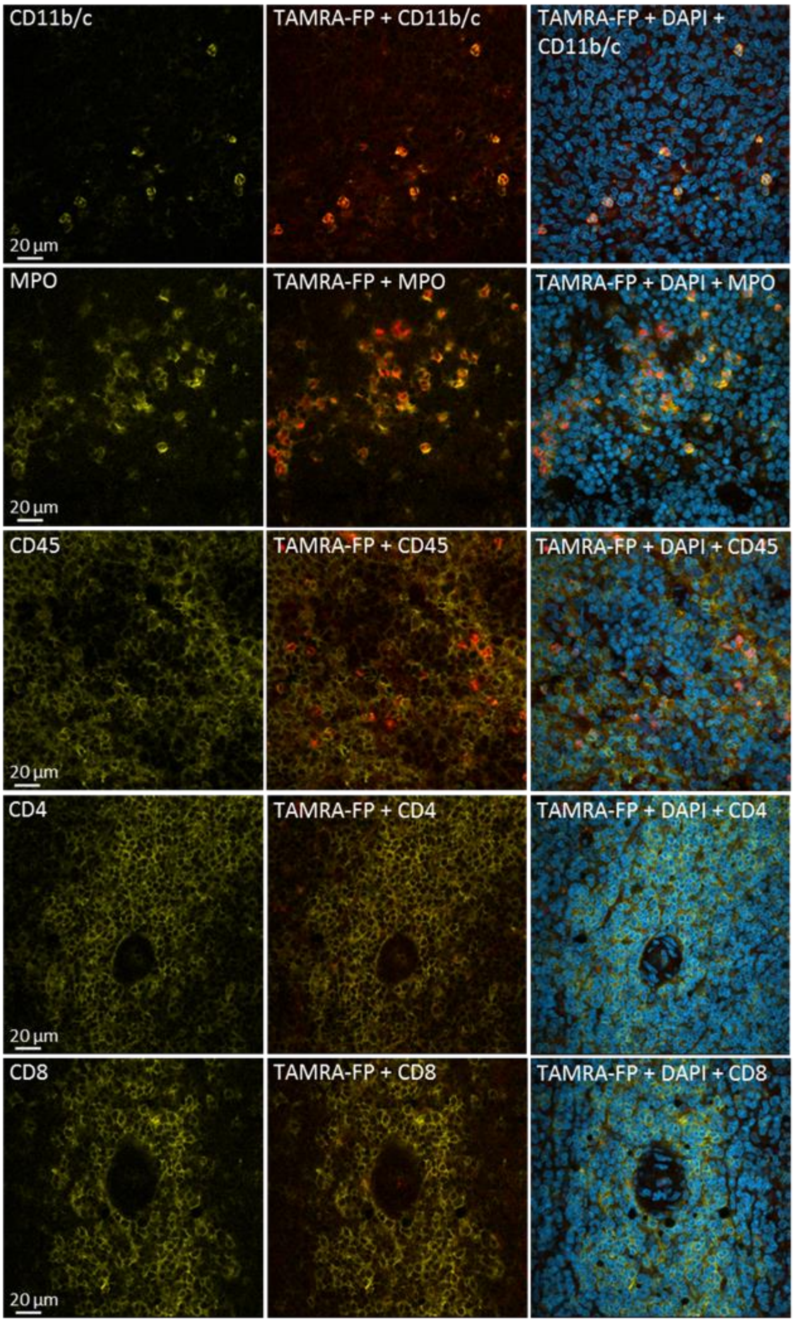


**Figure S35. Confocal imaging of SH activity in rat spleen sections in relation to selected immunomarkers.** Sections went through the tissue-ABPP protocol to label SHs (red) and were thereafter immunostained for phagocytes (CD11b/c), neutrophils (myeloperoxidase, MPO), nucleated hematopoietic cells (CD45), or T cells (CD4 and CD8) (yellow), followed by DAPI staining to visualize nuclei (blue). Note in particular CD11b/c-positive and MPO-positive cells in the red pulp region of TAMRA-FP hotspots and close match of CD11b/c- and MPO-positive cells with the TAMRA-FP signal. CD45-positive cells partially match with TAMRA-FP signal in red pulp region. CD4- and CD8-positive cells are enriched in white pulp regions with no TAMRA-FP hotspots. Primary antibody mouse anti-CD11b/c (OX42, Abcam, ab1211), dilution 1:500, secondary antibody Donkey anti-mouse IgG-Alexa Fluor 647 conjugate, dilution 1:500; rabbit anti-MPO (Abcam, cat# ab9535), dilution 1:25, secondary antibody Goat anti-rabbit IgG-Alexa Fluor 647 conjugate, dilution 1:500; mouse anti-CD45 (MRC-OX1, Abcam, ab33923, dilution 1:500, secondary antibody Donkey anti-mouse IgG-Alexa Fluor 647 conjugate, dilution 1:100; mouse anti-CD4 (OX-35, Abcam, cat# ab33775), dilution 1:100, secondary antibody Donkey anti-mouse IgG-Alexa Fluor 647 conjugate, dilution 1:100; and mouse anti-CD8 (OX-8, Abcam, cat# ab33786), dilution 1:500, secondary antibody Donkey anti-mouse IgG-Alexa Fluor 647 conjugate, dilution 1:100. Scale bars 20 µm. Images were adjusted for brightness and contrast.

### Materials and Methods

#### Ethical issues

All experimental procedures for animal studies have been approved by the Committee for the Welfare of Laboratory Animals of the University of Eastern Finland and by the National Animal Experiment Board (ELLA), and were performed in accordance with all national and local guidelanes and regulations. Blood for neutrophil isolation was donated by one volunteer member of the research group.

#### Chemicals and Reagents

The following activity probes were used: TAMRA-FP (ActivX TAMRA-FP serine hydrolase probe, cat# 88318) and DeBi-FP (ActivX Desthiobiotin-FP Serine Hydrolase Probe, cat# 88317) were purchased from Thermo Scientific. The Cy5-labeled serine protease probes PK-DPP and V-DPP bearing a diphenylphosphonate (DPP) warhead (Edgington-Mitchell et al., 2017) were used in selected experiments (Figures S29-30). Similarly, the Cy5-labeled NSP probes bearing a diphenylphosphonate (DPP) warhead (Kasperkiewicz et al., 2017) were used in selected experiments (Figure S31).

The following inhibitors were used: AEBSF (Sigma, cat#A8456), PMSF (Sigma, cat# P7626), IDFP (Cayman Chemical, cat# 10215), MAFP (Sigma, cat#M2939), CTSG-I (Calbiochem, cat# 219372). Compound 22 was obtained from Joakim Swedberg (at that time in the Institute for Molecular Bioscience, University of Queensland, Brisbane, Australia).

For immunohistochemistry, various antibodies from different sources were used. The hyaluronan binding probe (bHABR) was prepared in-house (Tammi et al., 1994) (Figure S13). The following secondary antibodies were used: Goat anti-rabbit IgG-Alexa Fluor 647 conjugate (Thermo Scientific, cat# A-21245), Donkey anti-mouse IgG-Alexa Fluor 647 conjugate (Thermo Scientific, cat# A-31571), Donkey anti-goat IgG-Alexa Fluor 647 conjugate (Thermo Scientific, cat# A-21447), and DyLight 488 Steptavidin (Vector Laboratories, cat# SA-5488). Primary antibody source, dilution used in immunostaining and choice of secondary antibodies are given in the captions of the images showing immunostained sections (Figures S8-28 and S34-35). Nuclei were stained with DAPI (4',6-diamidino-2-phenylindole) (Sigma, cat# D8417). The BSA used was essentially fatty acid free (Sigma cat#A0281). All other chemicals were of highest purity available.

#### Cell culture

BT4C cells were grown in Dulbecco's modified eagle's medium DMEM (EuroClone, high glucose, with stable L-Glutamine, cat# ECM0103L) supplemented with 10% FBS (Euroclone, cat# ECS0180L) in a humidified cell culture incubator with 5% $CO_2$ environment. The cells were maintained and sub-cultured two-three times a week at 1:6 splitting ratio. For preparation of the cell suspension for rat malignant glioma model, cells were trypsinized and collected in normal culture media (DMEM supplemented with 10% FBS) followed by a brief centrifugation at 1000rpm, for 4 min at 4 °C, and re-suspended in fresh OPTI-MEM®I reduced serum medium (Gibco, cat# 31985-062). Cells were then counted and diluted in the same medium to $2x10^6$ cells/ml so that each 5µl of cell suspension contained approximately 10 000 cells. For collecting cell pellets, cells were grown in several T-175 flasks until they become 95% confluent. On the day of collection, the cell culture media was aspirated from each flask, washed with cold 1xPBS, and cells were collected mechanically by using cell lifter/scraper in presence of 5 ml of cold 1x PBS. Cell fractions collected from different flasks were then combined into a 50 ml conical tube, washed with 1xPBS, centrifuged at low speed (300xg, for 10 min at 4 ˚C) to obtain opaque cell pellet. The pellet was stored at -80 ˚C.

**Animals**

*Malignant rat glioma model*

In total 34 BDIX rats (18 males and 16 females, Envigo, Huntingdon, UK) weighing 127-267 g were used. The rats were anesthetized i.p. with ketamine (Ketalar®, 60 mg/kg) and medetomidine (Domitor®, 0.4 mg/kg) and placed in a stereotactic apparatus. After skin incision, a hole was drilled 1 mm posterior to the bregma and to the 2 mm right of sagittal suture. 10 000 BT4C tumor cells in 5 µl of OPTI-MEM®I reduced serum medium were injected with a Hamilton syringe within right corpus callosum (depth 2.5-3.0 mm). To avoid backflow, the injection was done slowly over 2-3 minutes. The needle was left in place for 2 minutes and then slowly removed. Skin incision was closed with stiches followed by s.c. injections of antisedative (atipamezole, Antisedan®, 1 mg/kg) and analgesic (carprofen, Rimadyl®, 5 mg/kg). Control rats were processed similarly, except that cells were omitted and only OPTI-MEM®I reduced serum medium was inoculated.

*Tissue harvesting from glioma animals*

The rats were sacrificed 28-38 days after inoculation of the cells (the rats bearing tumors 28-31 days after inoculation). Animals were stunned with $CO_2$ and then transcardially perfused with PBS. From majority of the rats, the whole brain was removed, dipped briefly in cold (-80 °C) isopentane and stored on dry ice. From six rats in total, the tumor and corresponding brain piece from contralateral cerebral hemisphere were separated and were frozen on dry ice. For bone marrow collection, total of four bones (Tibia, Femur, Humerus, and Radius) were collected from each animal and stored in cold 1x PBS for further processing.

*Control rats used for harvesting immune cells and tissue*

For collecting rat spleens, cerebella, and fresh blood as the source of neutrophils, 10-12 week-old male Rcc:Han WIST rats (Laboratory Animal Centre, University of Eastern Finland) were used. Rats were decapitated and blood was collected with a funnel into a 50ml conical tube soaked with EDTA (1.5-1.8 mg/ml of fresh blood). The blood was then processed within 2 hours of collection. The cerebella were dissected, dipped briefly in cold (-80 °C) isopentane and stored on dry ice. The spleens were collected, frozen on dry ice and then stored at -80 °C.

*Mice used in preliminary tissue-ABPP experiments*

To test fixatives and TAMRA-FP concentrations during tissue-ABPP method optimization (Figures S4-5), 4-week-old male JAXC57BL/6J mice (Laboratory Animal Centre, University of Eastern Finland) were used. Mice were decapitated, the whole brain was removed, dipped briefly in cold (-80 °C) isopentane and frozen on dry ice and then stored at -80 °C.

**MRI**

Magnetic resonance imaging (MRI) was used to verify tumor existence 12-14 and 22-24 days after inoculation of the cells. Anesthesia was induced with 5 % isoflurane in a mixture of 70:30 % $N_2O$:$O_2$ and was maintained at 1.5 % isoflurane. MRI scanning was done with a 7T Bruker PharmaScan system and ParaVision® 5.1 software (Bruker, Billerica, MA, USA). Turbo-RARE imaging sequence with the following parameters was used: TE 12.5ms/50ms, TR 4000ms, RARE factor 8, FOV 2cm, 256x256 matrix, slice thickness 1 mm with 15 slices. The images were processed with a home-build Matlab program Aedes (University of Eastern Finland).

**Cryosectioning**

Coronal rat brain sections, horizontal mouse brain sections and rat spleen sections (20 µm thick) were cut at -19 °C to -21 °C using a Leica cryostat (Leica Biosystems, IL, USA). The sections (3 sections per slide) were thaw-mounted onto Superfrost®Plus slides (Menzel-Gläser, Germany), dried for 1-3 h at RT under a constant stream of air and stored thereafter at -80 °C. Rat and mouse brain sections were cut according to Rat Brain Atlas (Paxinos & Watson 1998).

**Tissue-ABPP**

Tissue-ABPP was used either for imaging SH activity alone or in combination with immunohistochemistry and nuclear staining, as detailed below.

*Protocol for imaging SH activity*

The assay was performed at RT (20-22 °C) unless otherwise stated. Generally, the fixing and all washing steps were performed in large volume (~200 ml) by dipping the slides into indicated buffer. Other steps of the protocol were performed by pipetting the indicated volume of buffer on the sections and by incubating the slides in a humidified chamber placed on a low-speed horizontal shaker.

The slides (3 tissue sections per slide) were thawed under a constant stream of air and thereafter each section was lined with a mini PAP pen (Fisher Scientific, cat# 10464573). The slides were fixed with 4 % paraformaldehyde (PFA) supplemented with 0.01 % glutaraldehyde in 0.1 M phosphate buffer (pH 7.4) for 10 min and rinsed thereafter 2x5 min in 1xPBS. The assay protocol consisted of preincubations 2x10 min in the assay buffer (250 µl/section) containing 50 mM Tris-HCl, pH 7.4; 1 mM EDTA; 100 mM NaCl; 5 mM $MgCl_2$ and 0.1 % (w/v) BSA, followed by 60 min incubation in the assay buffer (250 µl/section) in the presence of inhibitors or 1 % (v/v) DMSO as a solvent. After washes 3x1 min in the assay buffer (250 µl/section), the slides were incubated for 60 min with TAMRA-FP in the assay buffer (95 µl/section, routinely 0.5 µM final concentration) and thereafter washed 3x10 min in 0.1 M phosphate buffer (pH 7.4). In some experiments, Cy-5 labelled probes were used instead of TAMRA-FP as detailed in Figures S30 and S31. In preliminary experiments, presented in Figures S1 and S4-S5, the slides were dried under constant stream of air and imaged thereafter using a Fuji gel scanner. In subsequent experiments, the slides were further stained with nuclear stain DAPI (2 µg/ml in 0.1 M phosphate buffer, 300 µl/section) for 15 min at 37 °C and washed 2x5 min with 0.1 M phosphate buffer (pH 7.4). Finally, the slides were embedded with Vectashield® Antifade mounting medium (cat# H-1000, Vector laboratories, CA, USA).

*Tissue-ABPP combined with immunohistochemistry and nuclear staining*

When immune markers were used, the preceding protocol for TAMRA-FP labeling is the same as described above. The TAMRA-FP labeled sections were washed 5x5 min with 1xPBS supplemented with 1 % (w/v) BSA (210 µl/slide). The sections were incubated overnight with primary antibody (195 µl/slide) at 4 °C, washed 3x10 min with 0.1 M phosphate buffer (pH 7.4) followed by 60 min incubation with secondary antibody (195 µl/slide). Both primary and secondary antibodies were diluted in 1xPBS supplemented with 1 % (w/v) BSA. Finally, the slides were washed 3x5 min with 0.1 M phosphate buffer (pH 7.4), stained with DAPI (2 µg/ml, 300 µl/slide) for 15 min at 37 °C, washed 2x5 min with 0.1 M phosphate buffer (pH 7.4) and embedded with Vectashield® Antifade mounting medium.

### Histology

To visualize tumor morphology, the sections were fist fixed with 4 % PFA supplemented with 0.01 % glutaraldehyde and then stained with Mayer's hematoxylin and eosin. Imaging was performed with a Zeiss Axio Imager M2 microscope (Carl Zeiss Microimaging, Jena, Germany).

### Fluorescence imaging

*Fluorescence scanning*

For gel imaging and overview of tissue-ABPP slides, ChemiDoc™ MP imaging system (Bio-Rad, Hercules, CA, USA) was used. To visualize TAMRA-FP, Cy3 blot application (602/50, Green Epi) was used. For Cy5 probes or immunostained sections, Cy5 blot application (700/50, Red Epi) was used. In the early phase of this study (Figures S1, S4-S5), TAMRA-FP-labeled sections were imaged using Fuji gel scanner ($\lambda_{ex}$ 552 nm/$\lambda_{em}$ 575 nm).

*Confocal microscopy*

The confocal microscope images were obtained with a Zeiss Axio Observer inverted microscope (10x NA 1.3, 40x NA 1.3 oil or 63x NA 1.4 oil–objectives) equipped with LSM800 confocal module (Carl Zeiss Microimaging GmbH, Jena, Germany). TAMRA-FP, DAPI, and secondary antibodies were imaged with 561 nm ($\lambda_{ex}$ 543 nm/$\lambda_{em}$ 567 nm), 405 nm ($\lambda_{ex}$ 353 nm/$\lambda_{em}$ 465 nm), and 640 nm ($\lambda_{ex}$ 653 nm/$\lambda_{em}$ 668 nm) lasers, respectively. Secondary antibody used with bHABR (Figure S13) was imaged with 488 nm ($\lambda_{ex}$ 495 nm/$\lambda_{em}$ 519 nm) laser. ZEN 2.3 SP1 (black) and ZEN 2.3 lite (blue) softwares (Carl Zeiss Microimaging GmbH) were utilized for image processing and image analysis.

### Rat Bone Marrow Cell Isolation

For bone marrow collection, total four bones (Tibia, Femur, Humerus, and Radius) were collected from each animal removing extra materials covering these bones, and stored in cold 1xPBS. The tip of each bone was cut with scissors, and flushed into a 50ml conical tube with ice-cold 1xPBS using a 25G-needle, until no red color was visible. The sample was then homogenized using a pipette, filtered through Corning® cell strainer (CORNING, cat# 431750, 40 µm nylon), and centrifuged at 443xg for 4 minutes at 4 °C to pellet the cells, and suspended in 4 ml of 1xPBS. Four ml of ice cold ficoll-paque plus (GE Healthcare, 17-1440-02, density 1.077±0.001 g/ml) was pipetted into a 10 ml transparent tube, and bone marrow cells (4ml) were then overlaid on top of ficoll-paque plus carefully avoiding mixing of the two layers, then centrifuged at 443xg, 20 °C for 40 min without-brake in a swinging bucket centrifuge. The mononuclear cells were collected from the interface of PBS (top layer), and Ficoll-Red blood cells with granulocytes (two bottom layers). The cells were then washed by adding 3-6 volumes of 1xPBS followed by centrifugation at 1560xg for 4 min at 4 °C to remove extra Ficoll. The solid-white pellets were then stored at -80 °C.

**Human and rat blood neutrophil isolation**

For the isolation of neutrophils from human and rat blood, we used ficoll-paque plus (GE Healthcare, cat# 17-1440-02, Density 1.077±0.001 g/ml) density gradient protocol adapted from Kuhns et al., 2015 and Maqbool et al., 2011. Human and rat samples were handled separately. Briefly, fresh blood samples were collected from 5 rats (~33 ml) and a human donor (~12 ml) in sodium EDTA (1.5-1.8 mg/ml of fresh blood). Five ml of blood was overlaid carefully on 5.0 ml of ficoll-paque plus in a 15 ml conical tube, centrifuged at 500xg for 35 min at RT with brake off. The granulocyte-rich pellet was collected by removing three upper layers (plasma, mononuclear cells, and remaining ficoll-paque plus), and processed further by preparing 1x HBSS-buffer (LONZA, BioWhittaker, cat# 04-315Q) suspension, 3% (w/v) dextran (Dextran T500, Pharmacia, cat# 17-0320-01, 450,000-550,000 M.W.) sedimentation, collecting granulocyte-enriched supernatant to a new tube; sample was centrifuged (300xg for 10 min at RT) and after removing the supernatant, remaining erythrocytes were lysed using hypotonic buffer, re-equilibrated to restore isotonicity, followed by centrifugation (300xg for 10 min at RT). Final washing and centrifugation (300xg for 10 min at RT) was done using HBSS buffer, resulting to clear white pellet. The pellet was stored at -80 °C.

**Preparation of proteomes for gel-based ABPP**

Cell pellet was thawed in ice, and 100-500µl of 1xPBS was added to each tube depending on the size of the pellet (approximately 1 mm-sized pellet in height, was resuspended in 100µl of 1xPBS), mixed properly, spun, and went through freeze-thaw lysis procedure for at least five cycles (freezing at - 80 °C for 15 minutes or on dry ice for 5 min, thawing at 37 °C water bath for 1 min). Tissue samples were thawed in ice, weighted and required amount of 50 mM Tris-HCl, pH 7.4 + 150 mM NaCl buffer was added based on 1ml volume per 1g wet tissue-ratio, homogenized in glass-glass homogenizer in ice. The samples were collected into separate 2 ml Eppendorf tubes after washing with double volume of the Tris-buffer used initially. Finally, homogenized tissue samples and freeze-thaw-treated cell suspensions were sonicated on ice to obtain completely homogenized samples. Samples were then aliquoted, protein concentration was determined using BCA-200 Protein Assay Kit (PIERCE, 23226) as per company's protocol, and stored at -80 °C. Rat cerebellar membranes were prepared as previously described (Kurkinen et al., 1997).

**Competitive gel-based ABPP**

Competitive gel-ABPP was conducted using the Cy3-labeled probe TAMRA-FP or in selected experiments (Figures S29 and S31) various Cy5-labeled serine protease probes, following general outlines of our previous publication (Navia-Paldanius et al., 2012). Briefly, 25 µl of proteomes (5–10 µg protein) diluted in PBS were preincubated with the indicated inhibitors (50-fold desired final concentration) or a vehicle (DMSO) for 1h at RT, followed by addition of activity probe (in the case of TAMRA-FP, final concentration was 1 µM). The reaction was quenched by adding 2xSDS-loading buffer, followed by protein separation by SDS-PAGE (10%). For some lanes, Precision Plus Protein™ standards (BIO-RAD, cat# 161-0373) were included. In early phase of this study, TAMRA-FP fluorescence was imaged using Fujifilm FLA-3000 laser fluorescence scanner (Fujifilm, Tokyo, Japan) (Fluor. 532 nm; Filter: O580 nm). Thereafter, ChemiDoc™ MP imaging system (BIO-RAD, Hercules, California, USA) was used as follows: Cy3 blot application (602/50, Green Epi, Manual Exposure 10s-120s) was used, and Cy5 blot application (700/50, Red Epi, Manual Exposure 1s) was used to image MW markers and all Cy5-labelled probes.

**Sample preparation for LC-MS/MS**

Gel casting and ABPP were done with special care in order to minimize potential contamination by foreign proteins. Gel electrophoresis and imaging was as described above. As TAMRA-FP labelling is visible only in imager under specific wavelength, the cutting of intended bands was done by keeping gel in a new, clear square BioAssay Dish (Corning, cat# 431111, 245 mmx 245 mm) put on a horizontally positioned computer screen where the imaged picture was opened and positioned accurately (as accurate as possible). The gel pieces was cut using scalpel no. 11 and collected into labelled 1.5ml Eppendorf tube. The cut gel was then imaged again to check the accuracy of sample collection (Supplementary File 1_LC-MS-MS data). Collected samples were then wrapped with Parafilm®, stored in -80 °C freezer, and shipped to Helsinki proteomics lab in dry ice contained box.

**LC-MS/MS analysis**

Protein bands were cut out of the polyacrylamide gel (Bio-Rad, USA) and "in-gel" digested cystein bonds were reduced with 0,045M dithiothreitol (Sigma, cat# D0632) for 20 min at 37 °C and alkylated with 0,1M iodoacetamide (Sigma-Aldrich, cat# 57670) at room temperature. Samples were digested by adding 0,75 µg trypsin (Sequencing Grade Modified Trypsin, V5111, Promega). After digestion peptides were purified with C18 microspin columns (Harvard Apparatus) according to manufactures protocol and re-dissolved in 30 µl.

Liquid chromatography coupled to tandem mass spectrometry (LC-MS/MS) analysis was carried out on an EASY-nLC1000 (Thermo Fisher Scientific, Germany) connected to a Velos Pro-Orbitrap Elite hybrid mass spectrometer (Thermo Fisher Scientific, Germany) with nano electrospray ion source (Thermo Fisher Scientific, Germany). The LC-MS/MS samples were separated using a two-column setup consisting of a 2 cm C18-A1 trap column (Thermo Fisher Scientific, Germany), followed by 10 cm C18-A2 analytical column (Thermo Fisher Scientific, Germany). The linear separation gradient consisted of 5% buffer B in 5 min, 35% buffer B in 60 min, 80% buffer B in 5 min and 100% buffer B in 10 min at a flow rate of 0,3 µl/min (buffer A: 0,1% TFA in 1% acetonitrile; buffer B: 0,1% TFA acid in

98% acetonitrile). 6 µl of sample was injected per LC-MS/MS run and analyzed. Full MS scan was acquired with a resolution of 60 000 at normal mass range in the orbitrap analyzer the method was set to fragment the 20 most intense precursor ions with CID (energy 35). Data was acquired using LTQ Tune software.

Acquired MS2 scans were searched against rattus norvegicus protein data-base using the Sequest search algorithms in Thermo Proteome Discoverer. Allowed mass error for the precursor ions was 15 ppm. And for the fragment in 0,8 Da. A static modification parameter was set for carbamidomethyl +57,021 Da (C) of cysteine residue. Methionine oxidation (+15,995 Da (M)) and TAMRA-FP (+659,312 Da (S, Y)) were set as dynamic modifications. Only full-tryptic peptides were allowed for scoring maximum of 1 missed cleavages were considered.
