## Supplementary Video related to Fig. 1 for "Tissue-ABPP enables high-resolution confocal fluorescence imaging of serine hydrolase activity in cryosections – Application to glioma brain unveils activity hotspots originating from tumor-associated neutrophils"

### Slide 1
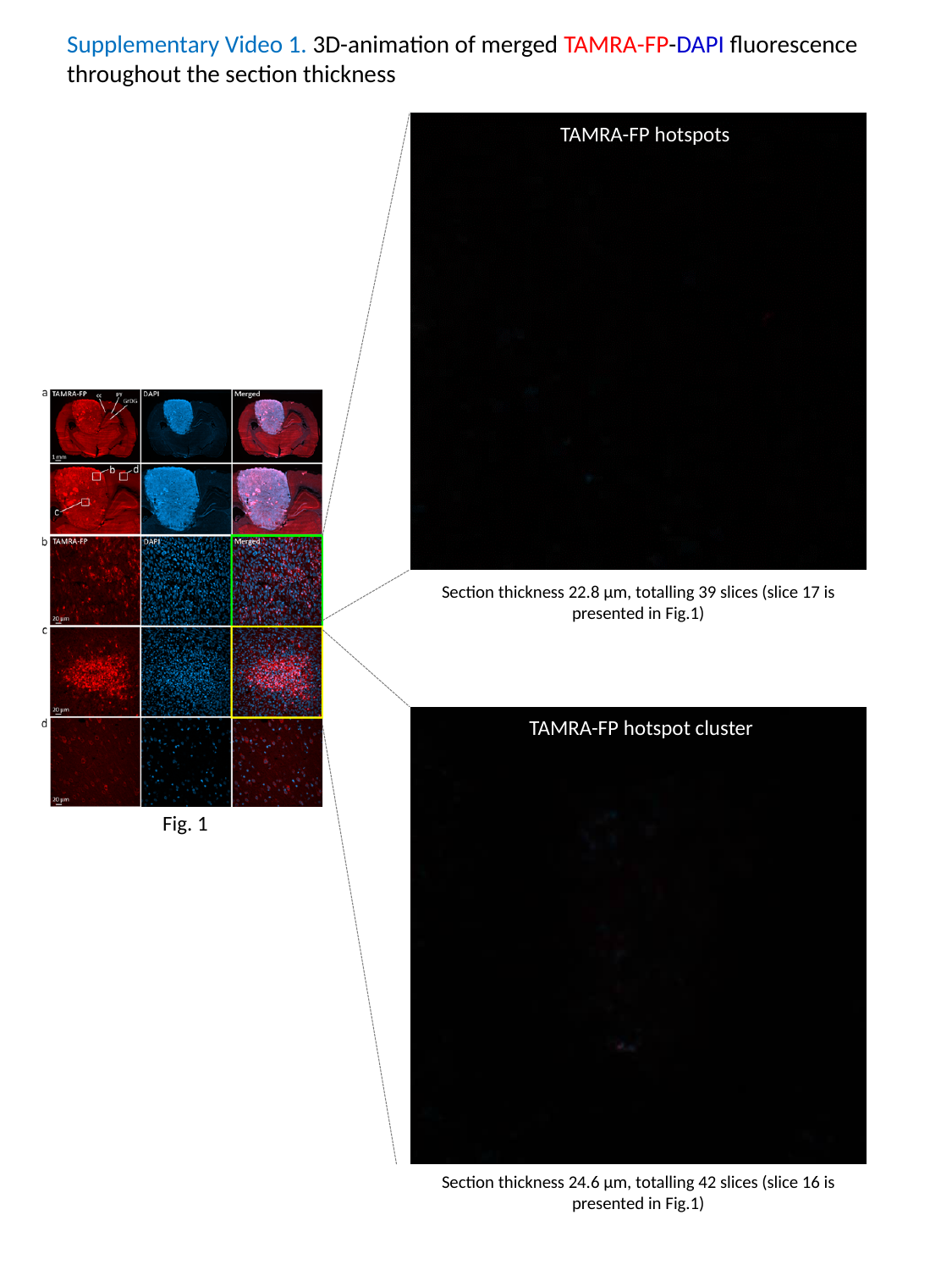

Supplementary Video 1. 3D-animation of merged TAMRA-FP-DAPI fluorescence throughout the section thickness
TAMRA-FP hotspots
Section thickness 22.8 µm, totalling 39 slices (slice 17 is presented in Fig.1)
TAMRA-FP hotspot cluster
Fig. 1
Section thickness 24.6 µm, totalling 42 slices (slice 16 is presented in Fig.1)
